# From field naturalism to Bayesian models: fog-frost interaction shapes growth form partitioning along Himalayan gradient

**DOI:** 10.64898/2026.06.25.733531

**Authors:** Pema Wangda, Melissa Whitman, Masahiko Ohsawa, Peter S. Ashton

**Affiliations:** Bhutan For Life Fund Secretariat/Executive Director, Box 1140, Thimphu, Bhutan; School of Earth, Environment and Society, Portland State University, United States; Laboratory of Biosphere Function, Graduate School of Frontier Sciences, University of Tokyo, Japan; Organismic and Evolutionary Biology, Harvard University, United States

**Keywords:** Bayesian beta-regression and Dirichlet compositional models, ephemeral cloud immersion, growth form partitioning, Bhutan Himalaya, tropical-temperate forests, uncertainty propagation (multiple imputation)

## Abstract

Mountain gradients facilitate our understanding of species’ range limits, competition dynamics, stress-resilience trade-offs, and determinants of vegetation zone boundaries. Forest compositional models often use altitude as the main predictor, a proxy for temperature that is defensible where floristic transitions are gradual and climate relationships are linear. However, mountains with distinct assemblages, representing tropical gradients or areas with complex biogeographic history, require a modeling framework that reflects non-linear dynamics or interactions between environmental factors, including outlier events (rather than mean conditions). Our study system encompasses both tropical and temperate forests along a broad (∼3000 m) altitudinal gradient, positioned within a narrow latitudinal band (< 1°) and composed of mature, continuous forest in the Bhutan Himalaya. To represent the breadth of climatic conditions experienced over a tree’s lifetime, we used a Bayesian modeling paradigm and integrated multi-generational field knowledge to develop *a priori* hypotheses and informed priors, with consideration of monsoon seasonality and possible ecophysiological thresholds. Our approach followed three stages (the Pattern, the Mechanism, the Test). Specifically, we interpolated microclimate data and derived custom metrics based on thermodynamics, propagating uncertainty into subsequent models to test whether climate posteriors outperformed altitude in explaining growth form partitioning. For spatial patterns, we identified six distinct vegetation zones (encompassing 145 species from 57 families), with a mid-gradient peak in richness at the tropical-temperate transition zone, and convergence of deciduousness at either end of the gradient. For individual growth forms, abundance was tied to different ecological mechanisms, explained by adaptations to climatic stressors and competition trade-offs. For instance, evergreen broad-leaved dominance was linked to ephemeral cloud immersion, whereas tropical deciduous species were affiliated with higher vapor pressure deficit at lower altitudes. Most importantly, compositional (between-group) models showed that the *interaction* between frost events and fog probability (air saturation prior to the dry season) governed growth form partitioning more than any single factor; temperate deciduous species, confined to a narrow altitudinal band, exemplified this finding. Our methodological approach is transferable to other data-sparse mountain systems, and our results highlight the vulnerability of unique habitat types and montane endemics under climate change scenarios that alter the fog-frost dynamics.

**Second abstract in Dzongkha:** To see the second abstract in Dzongkha, the official language of Bhutan, please visit our Zenodo site: https://doi.org/10.5281/zenodo.19081441.

## Introduction

Tropical to temperate floristic transitions occurring within a short spatial range are rare, generally restricted to mountain ranges or prominent landform formations within narrow latitudinal zones, or areas with complex biogeographic histories (Grubb 1971, Kitayama 1992, Mangen 1993, Li et al. 2013, González-Caro et al. 2020, Whitman et al. 2021, Ashton et al. 2022), but they provide natural experiments for identifying the ecological mechanisms that determine species’ range-limits and the abruptness of forest transitions. At broader temporal or spatial scales, tropical to temperate ecotones can emerge where plant lineages with distinct geographic origins converge, and interchange freely, following the removal of migration barriers, or via the formation of landform bridges, typically following tectonic movement, glaciation events, or shifts in climatic conditions (Brown et al. 1996, Morley 2000, Richardson et al. 2012, Joyce et al. 2021). At the local scale, compositional turnover, such as the transition from deciduous to evergreen canopy dominance, can indicate the abiotic filtering of species based on species’ ecophysiological tolerances, such as diurnal temperature range, freezing events, moisture or nutrient availability, or other edaphic properties, all of which can vary or shift in relative frequency along environmental gradients (Delcourt and Delcourt 1992, Tang and Ohsawa 1997, Givnish 1999, Russo et al. 2008, Mayor et al. 2017, Zhu and Ashton 2021, Whitman et al. 2021, Gallou et al. 2023), as well as context-dependent biotic interactions that range from competitive to facultative (Lynn et al. 2019). The exact mechanisms that determine where tropical forest transitions to temperate forest, and how abruptly, remain poorly resolved, in part because few mountains still support mature, late-successional forest across the full altitudinal transition due to increasing anthropogenic pressures; where intact gradients persist, detailed microclimate data are often sparse or unevenly distributed a subset of the gradient. Analyses of forest composition along mountains commonly use altitude as the sole predictor (Sundqvist et al. 2013, Whitman and Russo 2024), or rely on annual or seasonal climate means derived from broad scale gridded climate data, often due to logistical necessity. Such analyses exclude more specialized metrics, such as the diurnal or seasonal variation in growing conditions, or the probability of stochastic events. Thus, models based on generalist predictors can underrepresent the complexity of growing conditions that define observed assemblages, especially if floristic transitions correspond with non-linear climatic dynamics along the gradient. For instance, temperature, a climate variable that often linearly declines with increasing altitude, can display a subtle non-linear curve when accounting for interactions with humidity, which peaks during the monsoon season and can create a buffer from differences in lapse rate (i.e. how much temperature declines with altitude), relative to drier seasons when temperature differentiation is amplified (Wangda and Ohsawa 2006b). Ephemeral cloud cover, which seasonally shifts in size, position, and intensity, further influences moisture availability beyond what is directly attributed to precipitation. Thus, a more inclusive and nuanced approach for forest composition modeling is to use detailed site-specific examination of microclimatic conditions, which can reflect subtle interactions between factors, to understand spatial dynamics between floristic groups. However, gaining detailed habitat information can be labor intensive or lead to sparse data coverage, leading to either interpolation or reliance on simpler predictive metrics, which is why we opted for an alternative uncertainty-aware Bayesian approach to this analytical challenge (Ellison 2004, Clark 2005, Lynn et al. 2019).

Our study system represents a ∼3000 m mountain gradient with a narrow latitudinal range (< 1°) along the southern front range of Bhutan Himalaya, with mature late successional forests protected via national mandates for conservation. Bhutan is one of the few places on earth that supports a tropical to temperate transition zone, and is situated at the northernmost margin of Asian tropical climate influence (Ohsawa 1987, Wangda et al. 2010, Dorji et al. 2016, Khandu et al. 2022). The exact climate mechanisms structuring Himalayan forest mosaics remain understudied relative to the biodiversity of the region, which is why predicting forest response to climate change is especially pertinent (Leng et al. 2026). To better understand the climatic mechanisms that govern forest composition along this gradient we developed a comprehensive approach toward ecosystem evaluation and statistical modeling, where hypotheses were informed by decades of cumulative field observation across the mountains of tropical and temperate Asia, spanning multiple researcher generations (Appendix S1). Rather than relying on broadly available climate indices or altitude as a proxy, we deliberately constrained our analytical framework to predictions grounded in this cumulative natural history knowledge, such as specific observations about where dew forms and persists, which winters cause canopy dieback, and how cloud layers dynamically shift with the seasons. Translating our experiential understanding of forest function into formal statistical tests requires a novel approach, bridging historic to contemporary perspectives of ecology. Accordingly, this monograph is structured in three stages: first, *The Pattern*, where we characterize species’ relative abundance, compositional shifts, and vegetation zone boundaries along the gradient; second, *The Mechanism*, where we create a customized Bayesian pipeline for climate interpolation, utilizing sparse field station data while preserving uncertainty through to derived metrics and subsequent models; and third, *The Test*, where we use growth form partitioning as a response variable for compositional models, where we formally evaluate whether hypothesized climate interactions outperform the parsimony of altitude alone as a sole predictor. Our overarching paradigm is that the depth of the analytical approach should match the depth of the ecological system it aims to test.

Regarding forest composition in relation to climatic predictors, we hypothesized the following, informed by field observations and literature, with consideration of potential ecophysiological thresholds: 1) Tropical deciduous species would have a higher representation within areas experiencing greater thermal and moisture stress, where seasonal loss of leaves facilitates competitive success under drought. 2) Evergreen broad-leaved species would dominate habitats with more stable conditions, buffered by fog-derived moisture and reduced frost exposure. 3) Temperate deciduous species would display a narrow window of opportunistic dominance, where seasonal loss of leaves represents a stress-avoidance strategy, associated with frequent freezing events and less consistent moisture sources. 4) Conifers would dominate areas of greatest environmental stress from cold, aridity, and seasonal range of conditions. 5) Because the competitive outcome between growth forms depends on which stressors or sources of reprieve co-occur, we predicted that interactions between climate factors would outperform altitude or any single predictor in explaining stand-level dominance.

## Materials and Methods

### Regional context and study area

Unlike the Andes, where latitudinal migration pathways cross climatic zones, the Himalaya lies within a narrow latitudinal band, meaning that species interchange has occurred predominantly westwards or eastwards, along corridors defined by climatic similarity (Tang et al. 2018, White et al. 2019). The Himalaya were formed by the collision of the Indian and Eurasian tectonic plates ∼50-55 million years ago, when India, formerly part of the southern supercontinent Gondwana, drifted northeastwards and collided with the Eurasian plate margin, and through subduction and crustal thickening produced the tallest mountains on Earth. Backed by the Qinghai-Tibetan Plateau at more than 4,000 m a.s.l., which, as a major low-pressure zone and seasonal heat source, exerts climatic influence on Asia’s tropical climate to the south, the mountains draw equatorial moisture as summer rains off the Indian Ocean. With the uplift of the Himalaya, moist conditions returned to the region, after a dry Oligocene, and by the Miocene the progenitors of tropical lower montane and warm temperate forests dispersed across the continent and diversified (Morley 2000, Ashton 2014). Monsoon seasonality now characterizes extensive parts of tropical Asia, which extend beyond the Tropic of Cancer, with extremes in moisture availability and temperature thus shaping the adaptive strategies of the region’s flora. During the summer months, when rainfall is most intense and prolonged, moisture envelops the mountain slopes from central Nepal eastwards to the northern and coastal mountains of Burma (Myanmar). Our study took place within this zone of exceptional rainfall, along the humid southernmost outer slopes of the Bhutan Himalaya (i.e. front range, hereafter referred to as the “moist slope”), with annual precipitation often exceeding 4000 mm (Dorji et al. 2016).

### Surveys of woody flora and vegetation zonation

The study transect represents a near-continuous stretch of mature late successional forest (Fig. 1; Appendix S2), with vertical stratification of tropical to temperate forest types along a steep altitudinal axis, rather than spatially spread out along a continent as is more common for tropical-temperate comparisons (Bracewell et al. 2024). Floristically, the study area includes diverse broad-leaved forests with biogeographic ties to North-East India and South-West China (Ohsawa 1987), regionally managed for the collection of non-timber forest products and grazing (Norbu 2000). For the forest surveys along the gradient, we used a protocol established by Ohsawa (1984), using a series of 28 plots spanning from Rinchending (450 m a.s.l.; 26°51’30”N 89°23’23”E) to the top of Gedu (3370 m a.s.l.; 27°00’07"N 89°32’05"E). Most plots were ∼ 100 meters apart and 400 m^2^ in size, with larger plots in areas of abrupt topography; we determined plot size differences contributed to negligible variance (in part because the response variable used in models was proportional), so it was excluded as a random factor. At all sites (*n* = 28), we surveyed and identified all trees or woody shrubs ≥ 1.3 m in height, and estimated relative basal area (RBA), which is a unitless ratio (percentage) calculated as the basal area (cm^2^) of a species divided by the total basal area of the plot (Appendix S3). In addition, our other goal was to identify distinct vegetation zones, thus we replicated the methods used by Wangda & Ohsawa (2006a, 2006b), where zonation is inferred from major branches of a compositional dendrogram. Specifically, we calculated Bray-Curtis dissimilarity (Bray and Curtis 1957) between plots using relative basal area as a measure of species abundance, followed by Ward’s hierarchical clustering (Ward Jr. 1963) using the ward.D2 algorithm with squared dissimilarities (Murtagh and Legendre 2014).

**Figure 1.**
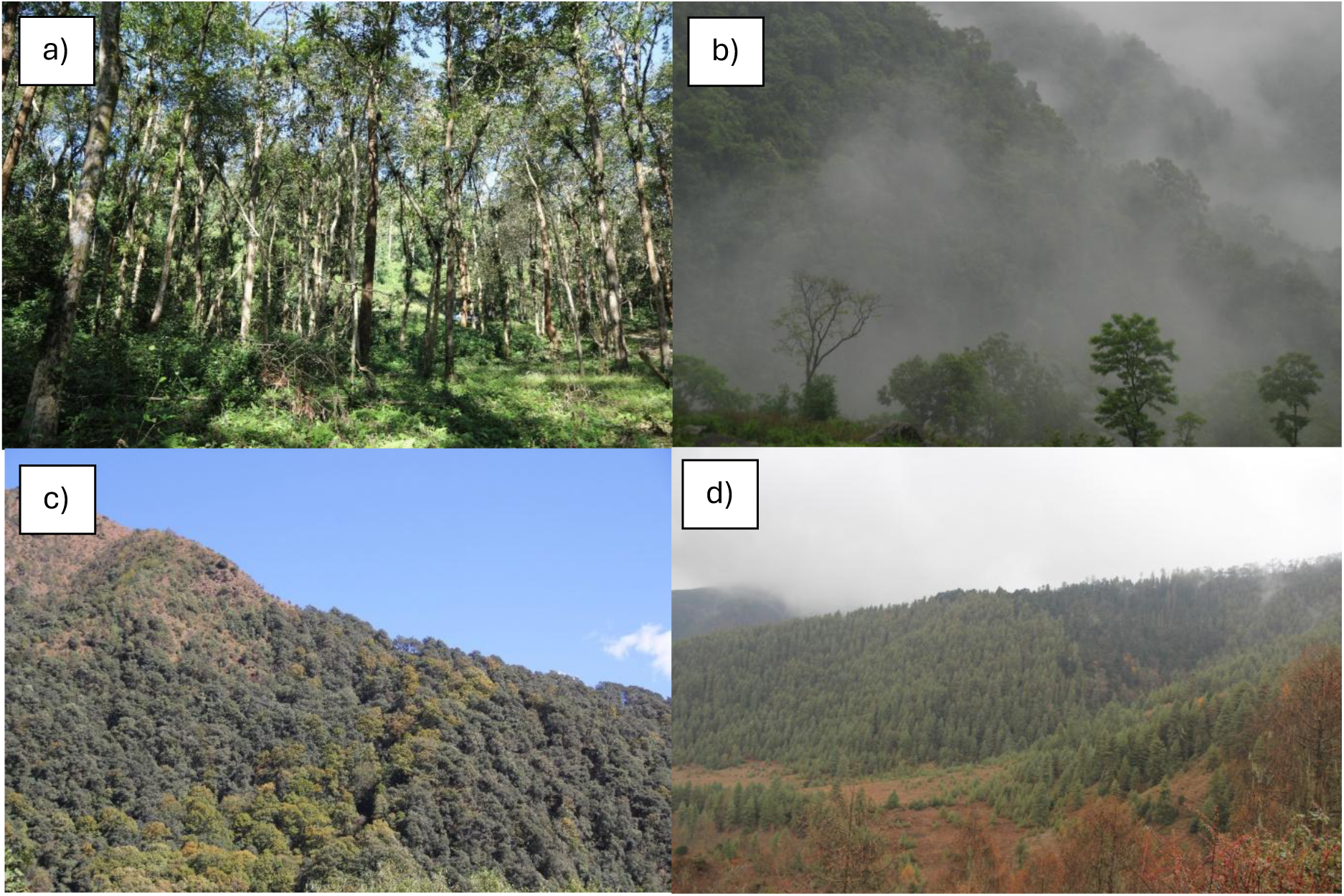
Forest types surveyed along the altitudinal gradient in Bhutan’s Eastern Himalaya. Forest types include the following: (a) Lowland forest with tropical deciduous species. (b) Mid-altitude forests, dominated by evergreen broad-leaved species, with the highest richness observed in the transition zone between tropical and temperate species, where the forest canopy is immersed in ephemeral clouds during the monsoon and transitional seasons. (c) Warm and cool temperate forest, showing a mixed canopy of evergreen broadleaf species (dark crowns) and deciduous species displaying autumn senescence (yellow–orange crowns). (d) High-altitude montane to alpine forest, with relative basal area (RBA) of the site dominated by coniferous species with more conical canopies. **Photos by Pema Wangda, Bhutan.**

### Interpolating climatic distributions along the gradient

At eight sites along the altitudinal gradient, we collected measurements of temperature (°C) and relative humidity (%) from daily data loggers, with information summarized as monthly averages and absolute minimum or maximum values, observed on any day of the month (see Appendix S4 for equipment and sampling details). For precipitation (mm), we collected data for three sites, further supplemented using weather station data summarized by Dorji et al. (2016). Collectively, these climate sampling locations spanned a slightly broader altitudinal gradient (∼210–3370 m a.s.l.), which facilitated interpolation (rather than extrapolation) of conditions across forest survey sites. In addition, we noted unusual outlier frost events from regional data spanning multiple years (Appendix S5).

Sparse or incomplete data present a recurring challenge in ecological field studies, thus to characterize climatic conditions at sites lacking direct microclimate instrumentation we used a Bayesian approach, analogous to data augmentation (Dumelle et al. 2025), informed by multiple imputation methods for propagating covariate measurement error (Wilson and Silander Jr 2014, Bartlett and Keogh 2018). Specifically, we generated full (1000 draw) posterior distributions, regarding individual draws as representative of plausible climatic conditions experienced over the lifespan of a mature tree. For intercept, we centered altitude at the transect median (1850 m a.s.l.), rather than sea-level, which minimizes extrapolation error and focuses on differences in habitat types relative to areas of the gradient noted to have the most climatic stability between seasons. Our climate interpolation model set included four levels of complexity, ranging from linear to natural spline with 4 degrees of freedom. For model selection, we used leave-one-out cross-validation (LOO-IC; Vehtari et al. 2017), combined with conditional logic to select richer non-linear relationships within reason (Appendix S6). Initial interpolations included 72 monthly models for temperature and relative humidity, and 4 seasonal precipitation models. Using interpolated posteriors and paired draws, we derived bioclimatic metrics including vapor pressure deficit, dew point depression (referred to as “fog gap”), and warmth index, with temporal aggregation. In total, we produced > 600 posterior files, designed to preserve the full distribution breadth of conditions experienced along the gradient.

Rather than using a data mining approach toward all possible posterior combinations as predictors, we instead referred to field observations and natural history knowledge as the means to identify factors and potential interactions of interest, and to hypothesize climate-forest dynamics and differences in forest response by growth form. We selected six predictors (deliberately selecting ∼1% of the data available) that conceptually matched our *a priori* hypotheses regarding the ecological mechanisms that determine species’ distributions and range-limits (Kira 1945, Holdridge 1967, Janzen 1967, Juvik and Nullet 1995, Still et al. 1999, Goldsmith et al. 2013, Grossiord et al. 2020, Feeley et al. 2020, Doughty et al. 2023), informed by decades of field research formalized as expert elicitation (Kaurila et al. 2026). These six climate predictors included metrics related to frost exposure, drought stress, fog proximity, cloud layer dynamism, thermal energy, and monsoon humidity amplitude, with the details and definitions of each metric shown in Appendix S6, and equation references in Appendix S9: Section S3. These factors reflect our prediction that deciduous trees, with the ability to lose their leaves during periods of stress, would have an advantage over evergreen species during high aridity events (e.g. conditions causing stomatal closure), such as high vapor pressure deficit scenarios during the spring transition from dry to wet season and warming temperatures. Alternatively, deciduousness can occur via evolutionary convergence but due to other environmental causes, such as within upper gradient areas where seasonal and diurnal stressors are more pronounced, where precipitation or moisture from cloud immersion is more infrequent but solar radiation and evaporation is high. In contrast, we predicted that evergreen broad-leaved species would often have a competitive advantage over other growth forms, facilitated by year-round growth and photosynthetic capacity, but this dominance would be conditional upon more stable growing conditions and higher moisture availability. Lastly, we anticipated the conifers would be most tolerant of extended periods below freezing, with the tapered canopy shape also better adapted to the weight of snow accumulation. As a method to evaluate these climate predictors prior to computationally-intensive forest compositional modeling, and to test whether climate explained compositional variation beyond altitude alone, we performed Mantel and partial Mantel tests, followed by distance-based redundancy analysis (dbRDA) and variance partitioning (Legendre and Legendre 2012). We regarded altitude a null predictor, representing neutral or random processes along the gradient, against which the explanatory power of climate mechanisms were compared (Tredennick et al. 2021).

### Modeling growth form partitioning along the gradient

To fully test whether observed forest composition was better explained by hypothesized climatic predictors, versus the simplicity of altitude-alone, we conducted a series of Bayesian models where relative basal area (RBA) was treated as a proportional value between 0 and 1 (Damgaard and Irvine 2019, Douma and Weedon 2019). Rather than using individual species RBA values, we simplified the response variable to represent growth form partitioning, categorizing species as broad-leaved evergreen, coniferous, or deciduous growth forms. Since the deciduous group displayed a bimodal diversity curve, we made tropical and temperate subgroups based on species’ average altitude of occurrence relative to the transect median (1850 m a.s.l.). First we tested individual growth form dominance as a function of climate using beta-regression models (i.e. each growth form is regarded as an independent unit), then we modeled dynamics between growth forms using Dirichlet compositional regression, where the gains by one group correspond to losses by others (Douma and Weedon 2019, Tremblay et al. 2021).

For the beta-regression models, we used partially informed priors and focused on parts of the gradient with species’ occurrence per growth form group to avoid zero-inflation. From a broader set of candidate models (with limited draws), we selected models for production runs utilizing all 1000 posterior draws, with results pooled using Rubin’s rules (Rubin 1987). We hypothesized directional response-predictor relationships *a priori*, reported as *P*(*β* in predicted direction), with complementary permutation-based one-tailed tests of observed distributions relative to spatially randomized null models (Manly and Manly 2018), see Appendix S7 and S9.

For the Dirichlet compositional models, which model all response groups simultaneously, we designated the evergreen broad-leaved group as the reference category because it is the dominant growth form. These Dirichlet models spanned the entire gradient, thus the larger sample size allowed us to incorporate more complex interactions between climate variables, elucidating potential stress-refuge trade-offs between groups. Similar to the beta-regression methods, we started analyses using a candidate model set, from which we carried forward a total of 10 climate models, and 2 altitude-only models (linear and quadratic) which served as null proxies, with full production runs across all 1000 draws and with results pooled per model. For model comparisons, we used a LOO-IC selection framework, prioritizing ecological plausibility over parsimony when results were similar (ΔLOO-IC < 2), see Appendix S8 and S9.

For all statistical analyses, we used R 4.5.3 (R Core Team 2026), with backend links to Stan, a probabilistic programming language compiled in C++, via R package “cmdstanr” (Carpenter et al. 2017, Gabry et al. 2026). All Bayesian models used an adapt_delta of 0.95 and maximum treedepth of 12 to ensure adequate posterior exploration given the complexity of some climate-altitude relationships; full convergence diagnostics are reported in Appendix S9. For tasks such as the development of checkpoint-based processing loops, memory management, or customized scaling of models or calculations, we used detailed prompts (Brown and Spillias 2026) to utilize AI-assisted coding tools (Claude Opus 4.6, Anthropic). All hypotheses, research design, analytical framework, and scientific interpretation were developed by the authors (see Appendix S9), with code and data provided for reproducibility (Appendix S10).

## Results

### Forest composition and vegetation zones along the altitudinal gradient

Across the gradient, spanning from 450 – 3370 m a.s.l., we recorded a total of 145 species from 57 families. The moist slope vegetation zones and dominant genera include: Zone 1, tropical lowland semi-evergreen *Terminalia-Duabanga-Schima* forest (450–850 m a.s.l.); Zone 2, tropical lower montane *Schima-Castanopsis* forest (850–1725 m a.s.l.); Zone 3, the tropical-temperate transition or *Quercus*-laurel forest (1725–2420 m a.s.l.), a diverse ecotone encompassing both the upper altitudinal limits of tropical lower montane species and the lower limits of warm temperate flora; Zone 4, warm temperate evergreen broad-leaved *Lithocarpus-Rhododendron* forest (2420–2850 m a.s.l.); Zone 5, cool temperate semi-deciduous *Sorbus-Rhododendron* forest (2850–3050 m a.s.l.); and Zone 6, cold temperate coniferous *Abies-Juniperus* forest (3050–3370 m a.s.l.). A more detailed list of indicator or endemic species for each zone is shown in Figure 2.

**Figure 2.**
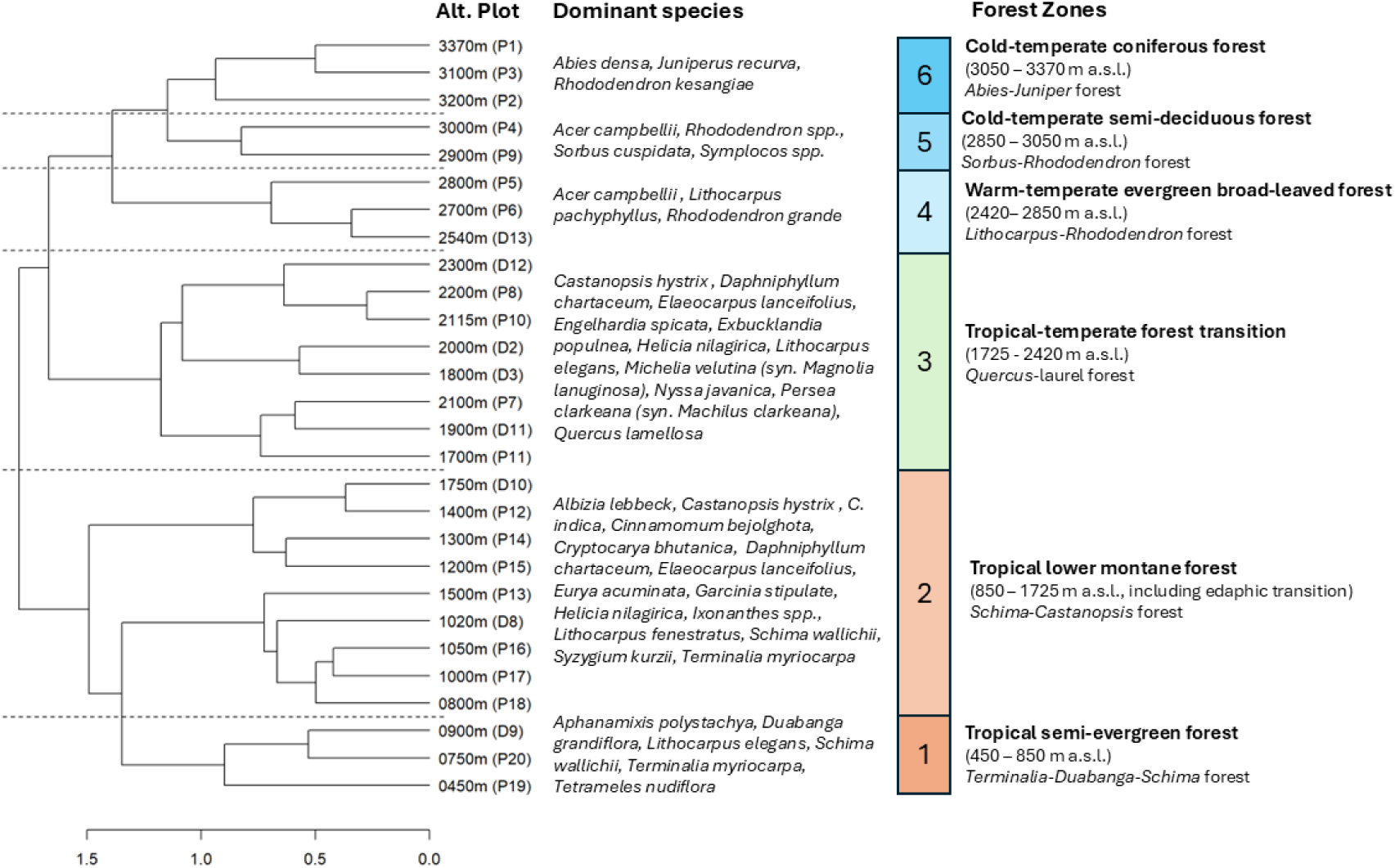
Vegetation zones identified using a hierarchical cluster analysis, with boundaries determined by the relative basal area (RBA) of species within each plot. The figure includes the altitude of each plot (m a.s.l), the proposed forest zones, and dominant tree species.

When categorizing species by growth form (Fig. 3), we found that the evergreen broad-leaved growth form was the most species-rich, representing 100 species from 43 families, concentrated within mid-altitude areas, peaking at 22 species at 1800 m a.s.l., with the greatest RBA (> 99%) occurring lower at 1050 m a.s.l.. Major evergreen families included Lauraceae (12 species, mostly trees from *Cinnamomum* or *Persea,* or shrubs from *Lindera*) and Fagaceae (8 species, including the genera *Castanopsis, Lithocarpus,* and *Quercus*), with Ericaceae as the main representative of montane richness (7 species, representing mostly *Rhododendron*). In contrast, the peak in tropical deciduous species richness occurred toward lower altitudes (8 out of 19 species occur at ∼ 800 m a.s.l., including genera *Albizia, Lagerstroemia,* and *Tetrameles*), which is where relative basal area was also highest (∼ 28%). The peak in temperate deciduous richness was broad, with 8 out of 25 species occurring at 3000 m a.s.l. and the largest RBA (∼65%) found at 2900 m a.s.l., representing species from *Acer, Gamblea, Magnolia,* and *Sorbus*. A notable outlier for the temperate deciduous group was *Betula alnoides,* occurring as low as 1020 m a.s.l. and representing ∼32% of the plot RBA. Conifers were restricted to alpine areas, becoming dominant above ∼3000 m a.s.l. (as high as 93% RBA), but displaying relatively low richness (only 2 species in total, *Abies densa and Juniperus recurva*).

**Figure 3.**
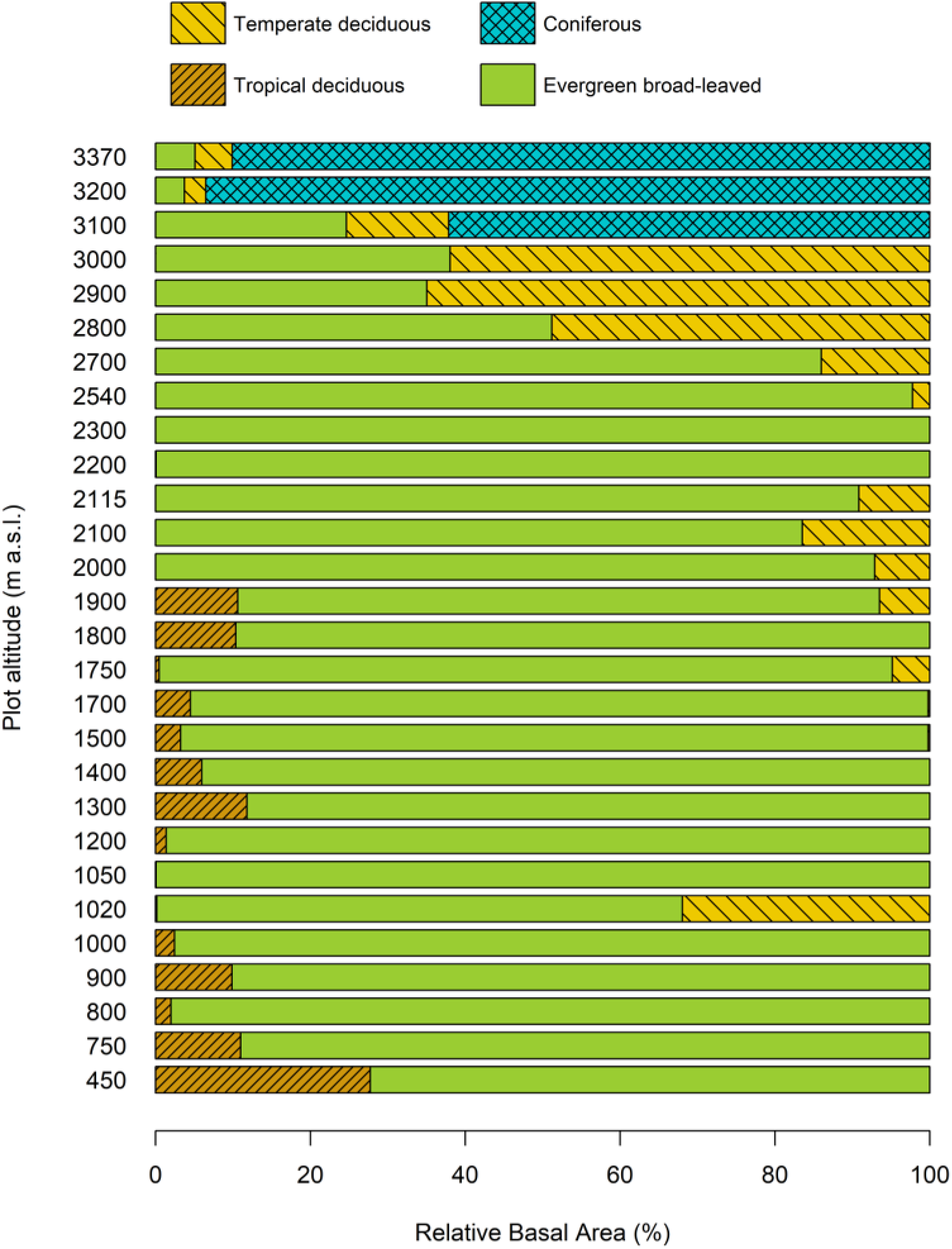
Relative basal area (%) of four growth form groups along the moist slope of the Bhutan Himalaya. All woody species taller than 1.3 m were measured for diameter at breast height (DBH) and converted to relative basal area per plot (*n* =28, spanning from 450 to 3370 m a.s.l.). Growth form groups: tropical deciduous (tan, right diagonal), evergreen broad-leaved (green, solid), temperate deciduous (yellow, left diagonal), and coniferous (cyan, diamond hatch).

### Climate as a predictor of community composition

After controlling for altitudinal distance between sites, we found that climate predictors explained more variation in forest composition than altitude alone, based on the partial Mantel test (*r* = 0.31, *P* < 0.001) followed by variance partitioning, where 25.1% of the explained variation was unique to climate as compared to 5.8% attributed to altitude (with 2.8% as shared). This finding occurred despite significant correlations between climate ∼ altitude (*r* = 0.86, *P* < 0.001), and composition ∼ climate distance (*r* = 0.605, *P* < 0.001) and ∼ altitudinal distance (*r* = 0.55, *P* < 0.001) respectively. The distance-based redundancy analysis (dbRDA) reinforced the importance of the six climate predictors (Appendix S6: Section S1-6), which explained 46.3% of the constrained variation (adj. R² = 0.31, *P* < 0.001), with each individual factor found to be significant in both the marginal and sequential permutation tests (Appendix S6: Section S7). The first dbRDA axis differentiated lower from mid-altitude sites (Fig. 4, 14.3% of the total variation), based predominantly on moisture metrics, and the second axis (11.0% of the total variation) was positively associated with sites experiencing freezing events or greater monsoon to dry season humidity range.

**Figure 4.**
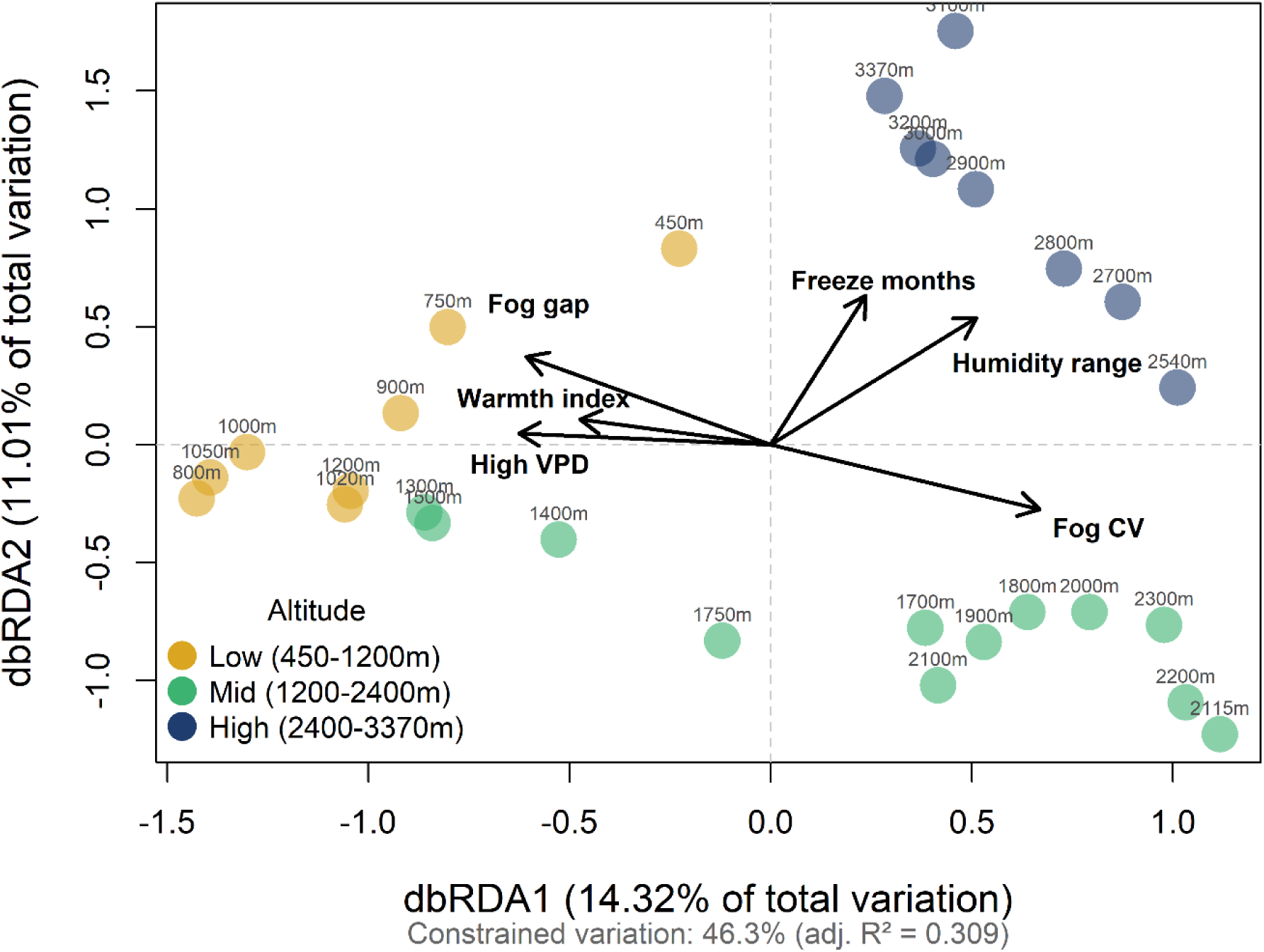
Distance-based redundancy analysis (dbRDA) of forest composition across the altitudinal gradient and in relation to predicted climate conditions of each site. To understand relationships between altitude, climate, and community composition of woody flora across sites (*n* = 28), we used site ordination, based on Bray-Curtis dissimilarity. Relative basal area (RBA) of woody species was used as the response variable, constrained by six climate predictors, selected a priori based on our hypotheses. Arrows indicate the direction and relative magnitude of climate predictor loadings, which are as follows: count of months at or below freezing (based on minimum temperature ≤ 0°C); count of months with high drought stress, described as VPD ≥ 1.5 kPa, as predicted at “noon” under maximum temperature and minimum humidity scenarios; fog gap, which is the difference between air temperature and the dew point, was estimated for the fall months at “dawn” under a scenario of minimum temperature and maximum relative humidity; warmth index, based on Kira 1945, which is the absolute lowest minimum temperature observed along the gradient; fog gap annual coefficient of variation (CV), based on capped values at “dawn” using monthly estimates; and average relative humidity range, based on the difference between monsoon and dry season conditions. The global model explained 46.3% of constrained variation (Adj. R² = 0.31, *F* = 2.979, *P* < 0.0001), and climate predictors explained 25.1% of unique variation as compared to 5.8% for elevation alone (variance partitioning). To visualize different parts of the gradient, sites are colorized and described as being: low (450–1200 m a.s.l., gold), mid (1200–2400 m a.s.l., green), or high altitude (2400–3370 m a.s.l., blue).

### Beta-regression models, with growth forms as independent units

Across all top beta-regression models, our *a priori* directional predictions were consistently supported, but growth forms with restricted distributions had wider credible intervals, sometimes crossing zero (Appendix S7: Table S1 and Section S2). Tropical deciduous species were prevalent at the lowest altitudinal areas of the gradient, with their RBA best predicted by months with more arid conditions (*n* = 19; *β* = 0.38, 95% *C.I.* -0.15–0.90; *P*[*β* > 0] = 0.92) with springtime extreme conditions exceeding the stomatal closure threshold (1.5 kPa). Tropical deciduous RBA was highly right-skewed, centered within plots from 450–900 m a.s.l. despite species occurring up to 2200 m a.s.l.. Evergreen broad-leafed species had a negative relationship with the seasonal range in relative humidity (*n* = 28; *β* = -1.19, 95% *C.I.* -1.88 to -0.50; *P*[*β* < 0] = 1.0), with greater RBA observed within stable habitats with consistent climatic conditions. For this metric, the lowest range in absolute relative humidity values occurred between 1050–1400 m a.s.l., with values ∼ 10.6% between the two seasons, as compared to values >20% within more montane areas at or above 2540 m a.s.l.. Similarly, evergreen species had higher RBA values in areas with higher fog gap CV (*β* = 0.63, *C.I.* -0.01–1.28; *P*[*β* > 0] = 0.98). The highest fog gap CV values (1.76) occurred at 2300 m a.s.l., which is the only plot where the minimum fog gap values average near zero annually (indicating saturated air), whereas areas that were consistently above or below the clouds had low fog gap CV values (Fig. 5). Based on climate posteriors, the ephemeral cloud layer spans from ∼2000–2800 m a.s.l., with the highest saturation probability during the monsoon, contracting to ∼2100–2400 m a.s.l. during transitional months and disappearing entirely in the dry winter. Temperate deciduous species also benefited from higher moisture availability, specifically with larger RBA values where the fog gap was lower during fall months, indicating supplemental moisture from cloud immersion (*n* = 23; *β* = -0.35, 95% *C.I.* -0.93–0.23; *P*[*β* < 0] = 0.92). Temperate deciduous RBA was also positively associated with frost occurrence, as indicated by the number of months below 0 °C (*β* = 0.61, *C.I.* 0.03–1.2; *P*[*β* > 0] = 0.95). Freezing events can occur as low as 2000 m, but they are more common at or above 2700 m a.s.l. where plots experience 1 to ≥4 months of freezing (with the highest frequency toward the summit). For conifers, instead of omitting them due to low sample size, we extended the analyses to areas of potential occurrence (Appendix S3: Section S1). We found that conifer RBA was positively associated with freezing months (*n* = 7; *β* = 1.35, *C.I.* -0.20–2.9; *P*[*β* > 0] = 0.99), albeit interpreted with caution.

**Figure 5.**
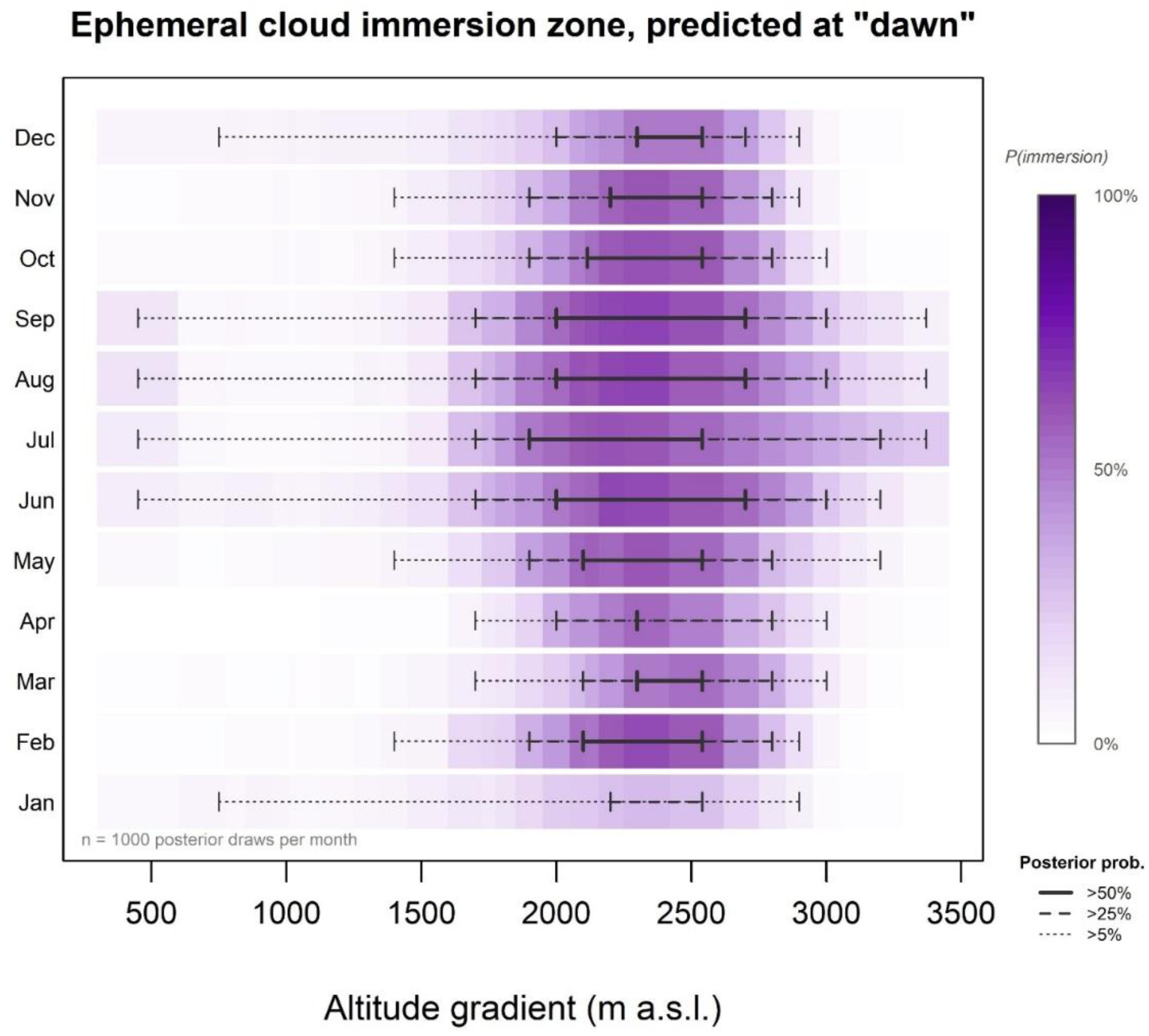
Ephemeral cloud immersion zone, as predicted along the altitudinal gradient. This figure illustrates “fog gap”, which is the difference between air temperature and the dew point on a monthly basis, under the “dawn” scenario with the combination of minimum temperature and maximum humidity observed for that time period. Each cell represents the posterior probability of cloud immersion along the gradient, calculated as the proportion of 1,000 Bayesian posterior draws in which the interpolated values were at levels associated with moisture saturation and the formation of dew on leaves (fog gap ≤ 0°C at dawn). Purple intensity reflects immersion probability from 0% (white) to 100% (deep purple). Horizontal bracket lines indicate the altitudinal range where immersion probability exceeded 50% (solid), 25% (dashed), or 5% (dotted) of posterior draws, estimated monthly. During the monsoon months (June to August, sometimes September), the cloud layer expands to lower altitudes, reaching to 450 m a.s.l. in some posterior draws. In contrast, during the dry season (December to February), the cloud layer contracts to a minimal presence, occurring primarily during nighttime and predawn hours, and is restricted to a narrow area ∼2100 – 2500 m a.s.l. This seasonal expansion and contraction of the cloud belt produces high coefficient of variation (CV) in annual fog gap at transitional altitudes and is one of the strongest predictors of the relative basal area of evergreen broad-leaved species.

### Dirichlet regression models, illustrating dynamics across growth forms

The best climate model outperformed the altitude-only model (ΔLOO-IC = 8.65 quadratic, 11.27 linear), indicating that compositional dynamics are better explained by ecological mechanisms than spatial turnover alone (Appendix S8: Table S1 and Figure S2). The top two climate models were very similar, with both showing an interaction between freezing event frequency and a fog refugia metric (Fig. 6), with fall fog gap scoring slightly better than annual fog gap CV (ΔLOO-IC = 2.99). Notably, these interaction-based models outperformed their additive equivalents using the same predictor pairs, indicating that the effect of frost exposure on growth form composition depends on moisture context, rather than operating independently. The winning model was selected based on overall compositional fit rather than individual coefficient significance. The one exceptional result for temperate deciduous species was their positive response to freezing events (*β* = 1.35, 95% *C.I*.: 0.60–2.11; *n* = 28), indicating reduced competition from evergreen broad-leaf species with greater frost exposure. Conifers responded similarly to freezing events (*β* = 0.85, 95% *C.I.:* 0.04–1.65), though with greater uncertainty due to extremely narrow range. Although individual fog coefficients had wide credible intervals, temperate deciduous and conifer species showed consistently opposing responses to fog metrics across models, with niche partitioning illustrated via the predicted effect sizes.

**Figure 6.**
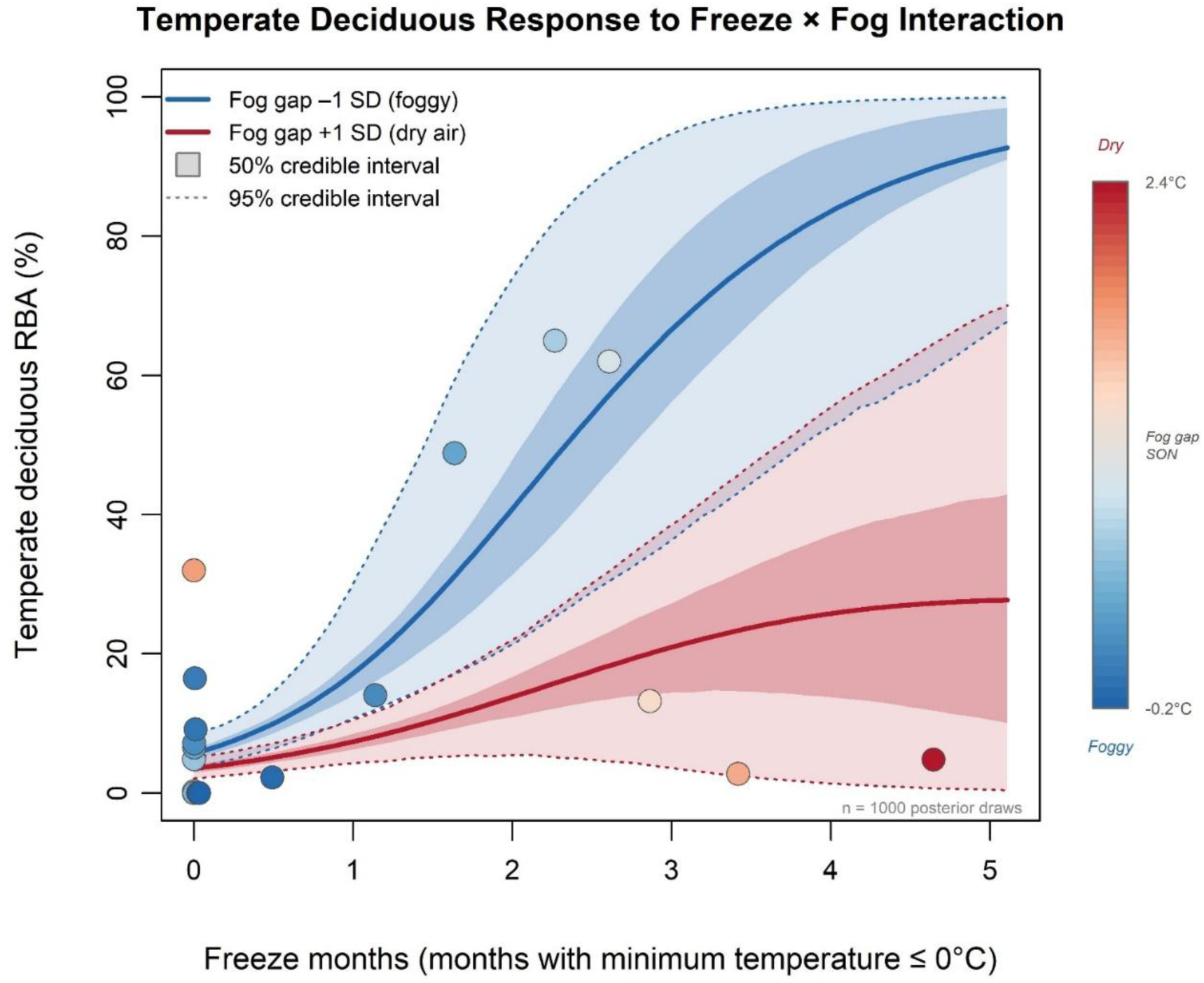
Temperate deciduous response to the interaction between freezing events and fog availability. Temperate deciduous response to the interaction between freezing events and fog availability. Predicted relative basal area (RBA) from the top Dirichlet compositional model, where freeze months (count of months with minimum temperature ≤ 0°C) interact with fall-season dew point depression (’fog gap’), a metric where values near zero indicate saturated air and cloud immersion conditions. The temperate deciduous growth form shows the most complex relationship with these two predictors, reflecting a trade-off between freezing exposure and moisture availability toward the upper edge of the ephemeral cloud layer. This climatic interaction may inhibit evergreen broad-leaved dominance within a narrow altitudinal zone, allowing higher proportions of temperate deciduous RBA before increasing abiotic stress leads to conifer dominance. Predictions are shown at fog gap conditions of +1 SD (dry air, red) and -1 SD (foggy, blue), with shaded ribbons representing 50% (darker) and 95% (lighter) credible intervals computed as pointwise quantiles across 1,000 posterior climate draws. Filled circles show observed RBA at the 28 survey sites, colored by mean fog gap value.

## Discussion

Relevant environmental conditions that limit taxa most can operate across different temporal scales or thresholds of exceedance (Janzen 1967, Still et al. 1999, Wang et al. 2010, Feeley et al. 2020, Aguirre-Gutiérrez et al. 2022, Doughty et al. 2023, Serafini et al. 2025), creating fitness trade-offs that are dependent upon overall life history strategies (Grime 1977, Givnish 2002, Givnish Submitted, Russo et al. 2008). Forest composition can be further enriched or reduced via competition between groups and compounding of stressors, either of which can shape forest mortality and generational composition (Ashton 2014, Davies et al. 2021, Ashton et al. 2022). Our results demonstrate that forest growth form partitioning along this gradient is governed by specific climate mechanisms rather than altitude per se, with the interaction between frost exposure and ephemeral cloud immersion emerging as the central finding across multiple independent analyses. Below we discuss the ecological interpretation of these mechanisms, their relationship to disturbance dynamics and the mid-elevation diversity peak, and their implications for montane research globally.

### The ecotone of Bhutan, blending tropical and temperate flora

Consistent with our three-stage framework, we begin with the Pattern: the floristic context needed to interpret the statistical models that follow. Within the Himalayan foothills, forests are often characterized by tall trees with large buttresses and mesophyll or macrophyll leaf sizes (Proctor et al. 1998). *Schima wallichii* is the dominant evergreen species characterizing lowland tropical forests. Emergent Dipterocarpaceae, characteristic of equatorial forests, were absent, although Sal forests (i.e. *Shorea robusta*) do occur in adjacent areas near the southern Bhutan-Indian border, affiliated with hilltop ridges or distinct soil types (Gyaltshen et al. 2014). The long-lived pioneer *Tetrameles nudiflora* was the only species to become emergent, the mature canopy otherwise being remarkably uniform. Other notable deciduous tropical genera sampled included *Pterospermum, Duabanga, Amoora (Aglaia), Terminalia,* and *Bombax*. Moisture stress tolerance further differentiates lowland assemblages, with tropical deciduous species affiliated with areas having freely draining siliceous soils (Ashton 2014, Zhu et al. 2021), or as we noted, high vapor pressure deficit values in the spring or in areas within the rain-shadow of the mountain face. Species’ response to disturbance events, namely fire intensity or frequency, also shapes forest composition of tropical lowland zones, with evergreen species showing higher fire-related mortality (Trouvé et al. 2020). Himalayan forests at comparable altitudes, with drier conditions and higher fire frequency, are typically dominated by *Pinus roxburghii* (Wangda and Ohsawa 2006b), a species that was absent from our transect.

At lower altitudes, the tropical lower montane forest (850–1725 m a.s.l.), sometimes referred to as “subtropical wet hill evergreen forest” by Champion (1936) or “middle hill forest” by Sharma et al. (2019), is a zone where lowland families with a more rugose canopy are replaced by a homogeneous main canopy dominated by Fagaceae, while Lauraceae characterize the subcanopy. Other taxa characterize the lower montane forest, notably Hamamelidaceae, and genera such as *Daphniphyllum, Engelhardia, Magnolia, Photinia, Schima,* and *Turpinia*. Belowground, the shallow organic horizon and termite-mediated decomposition characteristic of lowland forests gives way to earthworm-dominated soils within lower montane to warm temperate forests, where litter is ingested and carried to depth, resulting in darker humus-stained loam. Most flora within this altitudinal band are biogeographically associated with Indo-Burma or warm temperate forests of south Asia north of the Gangetic Plain (Morley 2000), with eastwards affiliation longitudinally via migration corridors that connect climatically similar habitats (Tang et al. 2018, Ashton and Zhu 2020). However, we noted minimal floristic connections to western Himalaya, which are closer to the dry Oligocene conditions of central Asia, with conditions that contrast with monsoon-dominated biomes. Similarly, most taxa sampled were not shared with the non-Himalayan portions of the Indian subcontinent, with the exception being a small subset of early successional or potentially invasive species colonizing landslide areas at lower altitudes (e.g. *Macaranga peltata*).

Mid gradient, at the tropical-temperate transition (1725–2420 m a.s.l.), the forest is dominated by broad-leaved evergreen species. Within this altitudinal band, the warm diurnal orographic updrafts keep clouds above the canopy, which cool and condense leading to canopy fog intrusion on upper slopes and ridges, most notable in late morning. The canopy then filters water as drops which leach and acidify the soil, podzolizing it and removing nutrient availability, yielding associated floristic and structural changes. A subtle compositional shift also occurs near ∼2100 m a.s.l., coinciding with peak species richness and marking the altitude where frost events begin to occur, though stochastically, with implications for the upper limits of tropical canopy species. The vegetation zone above (2420–2885 m a.s.l.) is a “warm-temperate evergreen forest” or simplified to “oak-laurel forest” based on the dominant canopy species (Grubb et al. 2013, Tang et al. 2018), transitioning into cool temperate forests with increasing altitude. This zone can also be referred to as “upper hill forest” (Sharma et al. 2019), “laurophyll forest”, or “lucidophyll forest” referring to the shiny or waxy leaf surfaces characteristic of the dominant species (Kira 1991). The growing conditions are analogous to the mountain plateau habitats of southern Yunnan, albeit Bhutan experiences relative floristic poverty because most tropical Asian species are at their westernmost range limits (Tang et al. 2018).

At the upper parts of the gradient, above ∼2900 m, we recognized three temperate forest zones that are more continuous and gradual in nature, transitioning from cool temperate with mixed evergreen-deciduous elements (*Quercus, Prunus, Sorbus, Betula*), to cold temperate conditions with conifers (*Abies*, *Juniperus, Tsuga*), which is why the area is sometimes referred to as “*Rhododendron*-conifer zone”. Toward the mountain summit, warm-temperate species are replaced by cold-temperate flora and Himalayan endemics such as *Rhododendron kesangiae*, with the uppermost forest zone populated by subalpine conifers (e.g. *Abies densa*) and scrub that are tolerant of extended periods below freezing.

Shifting from vegetation zone boundaries to the four growth form metrics used in subsequent models, evergreen broad-leaved species represent the dominant growth form in nearly every scenario, except at the stressful gradient edges where deciduous species and then conifers prevail. Our finding matches broader observations of forests worldwide, with deciduous species functioning as secondary dominants that gain proportional representation only where stressors reduce competitive exclusion (Grime 1977, Tang and Ohsawa 1997, Givnish 2002). However, this model operates on climate-mediated logic without consideration of disturbance mediated dynamics, in part because most of the forest is described as late successional mature stands. Landslide scars can lead to the establishment of early successional species, typically with *Alnus nepalensis* and *Macaranga peltata* at lower altitudes or *Betula alnoides* at higher altitudes, that persist as emergent canopy trees well beyond its more montane centers of occurrence (Ohsawa 1991). We flagged *B. alnoides* as an unusual outlier, representing 32% of RBA at 1020 m a.s.l., an altitude where temperate deciduous affiliation is unexpected, illustrating that natural disturbances can create opportunities for pioneer deciduous species that would otherwise be excluded.

### Climate mechanisms outperform altitude as predictors of forest composition

We turn now to the Mechanism: the climatic processes underlying growth form differentiation. Growing conditions in Bhutan Himalaya can range from hot summer temperatures and heavy rainfall to dry cold winters, yet the severity of these conditions can vary based on relative position of any given habitat along the altitudinal gradient. From field observations, microclimate extremes can be mitigated by ephemeral cloud layers that envelop the middle swaths of the mountain, shifting in position and breadth across seasons, leading to the transient presence of copious dew on leaves, heaviest before dawn yet disappearing with the day (Ashton 2014). The importance of higher moisture was apparent for evergreen-broad leaved species, with natural history knowledge reinforced via quantitative results. In contrast, habitat edges at the lowest altitudes are subject to greater drought or heat stress, thus are the domain of deciduous growth forms, whereas the highest summits have frost events and diurnal extremes or aridity above the clouds, marked by a sharp coniferous transition. From environmental extremes, convergent functional responses arise, such as the loss of leaves during times of stress, observed across unrelated lineages at opposite ends of the gradient (Ohsawa 1987, 1991). Our beta-regression models, where each growth form is regarded as independent unit, reaffirms that different climatic predictors can emerge as important depending on the growth form question, where one group’s bane may be the boon of another.

Evergreen broad-leaved species, representing the most dominant growth form, have an abundance peak that coincides with areas associated with cloud immersion, estimated via fog gap at “dawn” when atmospheric saturation is most likely (Fig. 5; Appendix S6: Section S3 and S5). Evergreen broad-leaves species are also affiliated with wet equatorial cloud forests, with their canopies enveloped in mist that supports a rich layering of moss and epiphytes (Edwards and Grubb 1977, Bruijnzeel and Veneklaas 1998, Still et al. 1999, Ashton 2014). However, the mountains of Bhutan are somewhat different from tropical mountain counterparts, in that monsoon seasonality leads to extremes where the cloud layer can either be all enveloping across the gradient, or near absent during the dry season, leading to a more ephemeral rather than persistent cloud belt (Ohsawa 1987, 1991). Our findings suggest that fog-derived moisture subsidizes plant water balance beyond what typical precipitation metrics (e.g. rainfall) indicate. Even though our conditional estimate of ephemeral cloud immersion is based on thermodynamics rather than directly via sensors, it is ecologically more important relative to other climate metrics examined. A more robust method worth investigation for future studies would be to test predictions of cloud immersion, as compared to ground-truthed surveys of the moisture provided from these events.

### The fog-frost interaction

Lastly, the Test. Our unified finding, that climate predictors, namely ones that reflect plant ecophysiological thresholds, perform better than altitude alone for explaining observed forest composition, was reinforced across models, including partial Mantel tests, beta-regression, and Dirichlet models. Of these analyses, the most meaningful result was that it is the *interaction* between fog and freezing events that matters, more than either factor in isolation. This concept is exemplified for the temperate deciduous growth form, where frost determines whether evergreen dominance gives way, but the buffering of additional moisture content via fog determines the outcome of their abundance, as well as their prevalence relative to conifers. Temperate deciduous species succeed in the narrow zone where frost and fog overlap, with seasonal leaf drop to mitigate frost damage, while fog-supplied moisture during the fall transition may support physiological preparation for the coming dormant season, such as via complex carbohydrate storage or prevention of cavitation as the air becomes drier. Similarly, during the other seasonal transition (the spring months) fog may also facilitate the rapid rebuilding of the canopy for deciduous species with a relatively short growing season.

Based on our top model, we predicted forest composition as 81.9% evergreen at the gradient median (1850 m a.s.l., with other growth forms in the minority), with 5.6% tropical deciduous, 8.5% temperate deciduous, and 4% conifer, with predicted values that differ slightly from observed proportions due to model smoothing, but sum to 1 by construction (Appendix S8). The top model provided useful effect size scenarios, such as when climatic conditions are colder and drier (+1 SD freeze, +1 SD fog gap), such as on a drier slope, then RBA for conifers increases to 19%, temperate deciduous to 13.3%, and tropical deciduous nearly the same at 7.3%, but with evergreen declining to 60.4%. However, under an extreme climate change scenario, where warmer and drier conditions are each increased by two standard deviations, compositional representation of montane growth forms is predicted to decline to less than 1% (Appendix S8: Figure S4).

### Other insights: diversity along the mountain gradient and conservation applications

Climate factors that influence the abruptness of vegetation zone boundaries or that differentiate growth form dominance along the gradient may also influence how diversity across species is distributed. The common "hump-shaped" diversity pattern reported on mountains is often attributed to mid-domain effects or the overlap of species ranges (Rahbek 1995, Colwell and Lees 2000, Bhattarai and Vetaas 2006, Grytnes et al. 2008), where the causes are geometric constraints rather than ecological mechanisms. Instead, we propose that the observed peak in richness (within the context of our study system) reflects two distinct filtering mechanisms operating at opposite ends of the gradient, with frost exposure at upper altitudes and heat stress at lower altitudes, both compounded by more arid conditions and elevated VPD above and below the cloud layer (Ashton et al. 2022). For instance, our beta-regression model that included stomatal closure threshold (i.e. VPD >1.5 kPa in springtime) directly predicted tropical deciduous dominance at the lowest elevations. Middle portions of the gradient can support more species because fog-derived moisture, rather than direct precipitation, buffers against the abiotic filtering that operates at both extremes (Juvik and Nullet 1995, Still et al. 1999, Goldsmith et al. 2013). A second scenario, which is not mutually exclusive but is worth further study, is that middle sections of the gradient support the transition from tropical to temperate floristic assemblages (Kluge et al. 2017, Thorne et al. 2022). Stochastic frost events, combined with niche partitioning under the context of habitat stability and specialization, would lead to a staggered arrangement of tropical to temperate species ratios, with areas supporting both groups having the highest diversity as the outcome (Ashton and Zhu 2020).

In Asian forests, the coldest mean temperature has long been correlated with the transition from broad-leaved to conifer-dominated zones, typically near -1°C at approximately 3200 m (Ohsawa 1990, 1992, Wangda and Ohsawa 2006a, 2006b), with the potential to disrupt tropical species more than temperate (Wang et al. 2010). Field observations along this gradient, specifically an outlier frost event at 2200 m a.s.l. (P.S. Ashton pers. obs., Appendix S1: Figure S2), reinforces our hypothesis that the upper altitude of tropical emergent and some canopy species in Bhutan is set by occasional heavy frosts which rarely penetrate the subcanopy, whereas the more gradual diminishing upper altitudinal limit, particularly of subcanopy species, is determined by individual species’ tolerance of increasingly frequent and intense nocturnal winter subcanopy frost. Events such as this can also cause mortality for seedlings or saplings, disproportionately impacting tropical species that are less tolerant of sudden temperature drops. Stochastic frost events may therefore cause cascading effects on recruitment, shaping forest composition for decades. While unusual, similar mid-altitude events have been observed throughout Bhutan (Appendix S5: Table S1), but these outliers can be missed if focusing on temperature mean values, either seasonally or annually. Similarly, the rate of temperature decline with increasing altitude (lapse rate) can be more dynamic than often assumed, shaped by relative humidity which fluctuates by season and location (Appendix S4: Figure S2). Bootstrap resampling of the Bayesian posteriors for extreme minimum temperatures during the coldest winter months revealed a sharp transition in frost probability along the gradient (Appendix S5: Section S2). The probability of freezing was near zero below 2000 m, increasing to 10% at 2300 m, 50% at 2540 m, and exceeding 99% above 2900 m, where temperate deciduous proportional representation (RBA) is highest.

Given the ecological complexity at the tropical-temperate transition, where species richness peaks and the ephemeral cloud layer most strongly mediates compositional outcomes, the establishment of a permanent forest dynamics plot(s) in partnership with the ForestGEO network (Davies et al. 2021), and long term microclimate sensors, would connect Bhutan to a global observatory for monitoring climate-driven change. As a world leader in sustainable landscape management, Bhutan can apply the findings of this study to several key areas. For instance, our research can be used to establish future protected areas, predict the response of species to changing environmental conditions, or to help develop landscape management strategies that meet the country’s constitutional forest-cover objectives. Other practical improvements would be the replication of this study globally, ideally contrasting the findings from Bhutan, representing an intermediary ecotone, with mountains situated within tropical and temperate latitudes respectively (Ashton et al. 2022).

### Broader methodological implications for montane research

Despite the importance of montane research, this field of study can be limited by habitat inaccessibility, lack of scientific infrastructure, and the need for specialized sensors (Juvik and Nullet 1995, Still et al. 1999, Fisher et al. 2013, Sundqvist et al. 2013, Mayor et al. 2017, Shih et al. 2025), which cannot be solved by remote sensing products whose resolution reflects the broader surrounding lowland landscape rather than steep montane gradients. The detailed microclimate data at some, but not all, of our research sites is what motivated our use of uncertainty propagation in subsequent models using climatic conditions as predictors. While multiple imputation (or similar Bayesian methods) most commonly address missing data for functional traits, phylogenies, or demographic information (Rubin 1987, Tremblay et al. 2021, Simmonds et al. 2024, Blomberg and Todorov 2025, Dumelle et al. 2025), this technique can also be applied toward interpolation of climatic conditions or spatial surfaces (Wilson and Silander Jr 2014). Unlike typical imputation scenarios, where data are assumed to be missing at random (Dumelle et al. 2025), sites that lack measurements represent structural design omissions, a distinction that is imperfect but not fatal, as the missingness is deterministic and fully explained by logistical constraints rather than probabilistic. Thus, when climate interpolations are treated as distributions, rather than trusting point estimates, then the result is a more honest reflection of site knowns and unknowns. Working with a limited number of sites with climate data did give us pause, but ultimately this limited information was still selected for over parsimonious yet less-informative options, being use of altitude only as the main predictor, retroactively validating our computationally intensive methods. Our methodological approach could be applied to similar mountain systems where station data are sparse, but the applied ecological stakes are high, as a step toward comprehensive forest management in the wake of climate change.

Our study reaffirms that microsite conditions are important for defining plant communities and their functional attributes (Givnish 1999, Camarero et al. 2021, Stark and Fridley 2022). Yet one limitation of our study is that we focused on modeling climatic conditions, rather than inclusion of edaphic characteristics or assessment of nutrient availability, which can strongly determine floristic assemblages and abruptness of compositional transitions along mountain gradients (Whitman et al. 2021). Similarly, this study inferred, but did not directly measure, metrics of biotic competition or facilitation, which would have better elucidated dynamics between growth form groups. We have confidence in our finding for evergreen broad-leaved species, which represent the most species and which occur across the entirety of the gradient, but our confidence in conifer results is reduced due to their relatively limited presence, even though the restricted range within areas of highest stress via freezing events is intuitive.

Under climate change, it can be hard to predict which taxa or habitat types are most at risk. Along mountain gradients, lower altitude communities can be threatened where heat exceeds ecophysiological limits (Aguirre-Gutiérrez et al. 2022, Doughty et al. 2023), or at the antipode, at summits where there is simply no more space to ascend upwards to cooler conditions (Sundqvist et al. 2013, Rehm and Feeley 2015, Mayor et al. 2017, Feeley et al. 2020, Camarero et al. 2021, Zu et al. 2021, Trethowan et al. 2023). However, it is safe to say that flora with limited distributions face higher odds of extinction, especially for assemblages with high endemism or unique trait characteristics (Steinbauer et al. 2016, Kidane et al. 2019, Gallou et al. 2023, Whitman and Russo 2024). Similarly, flora that are specific to more alpine conditions, including krummholz vegetation or *Rhododendron* shrublands with stunted growth forms which occur between the upper forests and the tree line can represent a unique set of environmental conditions that are near absent at lower altitudes (Korner et al. 1983, Ohsawa 1993, Mayor et al. 2017, Bader et al. 2021, Givnish Submitted). Within Bhutan, montane flora often have limited distributions, with notable species including *Rhododendron kesangiae* or the endangered *R. pogonophyllum* (pers. obs. R. Pradhan). Range restriction can also apply to unique habitat types, and as we found, applicable to the narrow zone characterized by temperate deciduous species, confined between contrasting influence of refugia and stress. From a modeling perspective, it makes sense then to include the tail ends of growing condition distributions, including the diurnal extremes rather than annual means (Zhang et al. 2022, Gallou et al. 2023, Shih et al. 2025), to best understand the habitat needs of these species.

## Conclusion

This paper represents the transfer of multiple generations of natural history knowledge, encompassing conversations between mentors and mentees regarding forest function, with the convergence of ideas into hypotheses, tested using modern Bayesian methods. The Himalayan gradient chosen as our study site is one of the few places on earth where the full tropical to temperate transition remains intact with predominantly mature forest stands, a circumstance attributable to Bhutan’s national conservation policy and commitment to future generations. If future conditions become warmer and drier, the frost line will move upward, the cloud layer will shift, and the climatic interaction that defines the narrow zone occupied by temperate deciduous species and montane endemics is likely to contract. Our quantitative framework is transferable to other data-sparse mountain systems, but the field knowledge and forests that informed these models can be more fleeting, ephemeral as the clouds which inspired this work.

## CRediT authorship contribution statement

**Pema Wangda:** Conceptualization, Data Curation, Formal Analysis, Funding Acquisition, Investigation, Methodology, Project Administration, Resources, Writing - Original Draft, Writing – Review & Editing.

**Melissa Whitman:** Conceptualization, Data Curation, Formal Analysis, Investigation, Methodology, Software, Validation, Visualization, Writing – Original Draft, Writing – Review & Editing.

**Masahiko Ohsawa**: Conceptualization, Funding Acquisition, Methodology, Project Administration, Resources, Supervision, Writing – Review & Editing.

**Peter S. Ashton:** Conceptualization, Funding Acquisition, Project Administration, Supervision, Writing – Original Draft, Writing – Review & Editing.

## Data availability

Curated data and code to fully replicate the findings of this study are openly available via Zenodo at https://doi.org/10.5281/zenodo.19081441.

## Acknowledgements

This paper is dedicated to the 70th Birth Anniversary of His Majesty Jigme Singye Wangchuck, fourth King of Bhutan, an international champion of forest conservation. We thank Rebecca Pradhan for sharing her knowledge of the plants and natural history of Bhutan, and the field research team, Sonam Tashi, Dorji Gyaltshen, Cheten Thinlay, Rinchen Dorji, Tshewang Norbu, and Damber K. Ghemiray for their valuable assistance with forest surveying. Dorji Gyaltshen, research officer, is duly acknowledged for his assistance in species identification. Peter S. Ashton thanks Virginia Medawar & Mary Oyeyemi for secretarial and computer assistance. Melissa Whitman thanks to Raymond Tremblay for Bayesian advice; Andrés Holz and Alex Fajardo for feedback; and Sabrina E. Russo and James O. Juvik for their encouragement to study tropical forests.

## Supporting Information

### Appendix S1: Section S1

#### Letter regarding the history of the study

I Peter S. Ashton, write in an editorial capacity, as explained in the following note at the end of this manuscript. I had the good fortune to meet Pema Wangda when Mary and I made our last visit to Bhutan, in November 2008 when we went to photograph trees and forests for my book, *On the Forests of Tropical Asia*. We saw and discussed his project at the site of his transect, especially at Gedu where the impact of severe frost on the crowns of some trees was something new for us to witness in the tropics. Four years later, I was puzzled that I had heard nothing of the expected publication, so I wrote to Pema who replied at once to explain that he was having difficulty in finding a journal that would accept his study. He then sent me his manuscript (which he had started in 2007), when I realized at once why: mainly that, as one might expect, it was too restricted to attract the interest of an international readership. But his achievements, inspired by methods which had been developed by his doctoral adviser Ohsawa Masahiko, are so remarkable that he and I have been working on the text off and on ever since. Fast forward to 2024, I ran into a long-time dipterocarp colleague, Sabrina E. Russo, who suggested that her first doctoral student, Melissa Whitman, would add a broader biogeographical component and more advanced statistical methods to the study. Her assistance with scientific statistics, writing, modeling framework, and data visualization helped the paper develop into what it is today.

### Appendix S1: Figure S2

#### Photos from Bhutan, by Peter S. Ashton

Image of severe crown damage from stochastic frost event at Gedu (∼2000 a.s.l.) in winter 2007-2008. from the transect at the tropical lower montane-warm temperate forest ecotone.

Originally published in *On the Forests of Tropical Asia* (2014), Plate 4.15a.

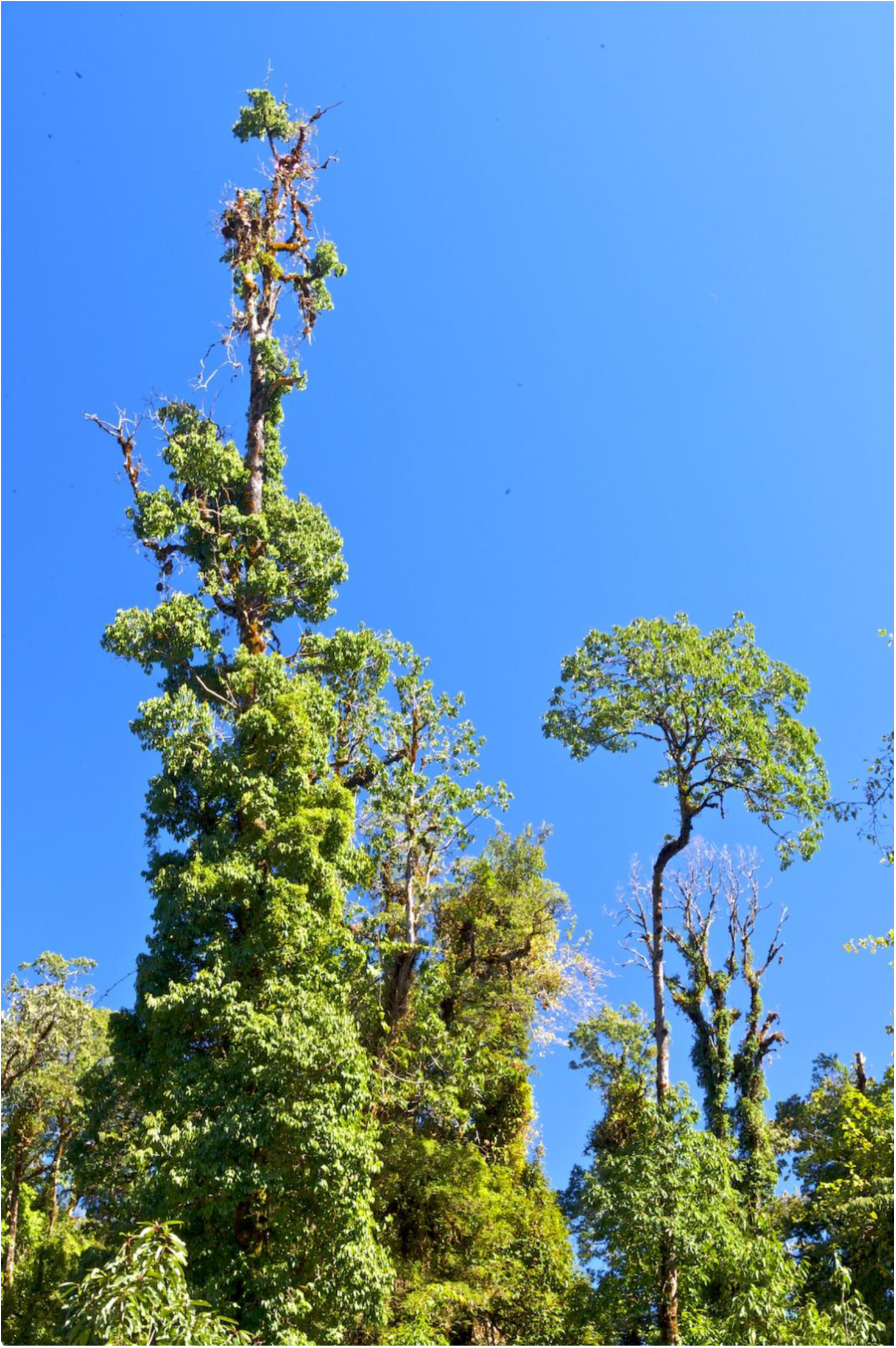

### Appendix S2: Figure S1

#### Map of sampling locations in Bhutan

Image includes: (a) Location of Bhutan within South Asia. (b) Schematic of Bhutan showing the humid outer slopes (study region, dashed line) and interior dry valleys. (c) Satellite imagery of the transect showing forest survey plots (P1–P20, D1–D13, yellow markers) and climate monitoring stations (M1–M8, blue markers) distributed along an elevational gradient from ∼450 m to ∼3370 m a.s.l. on the southern humid slopes between Phuntsholing and the upper ridgeline. Satellite imagery from Google Earth by Pema Wangda.

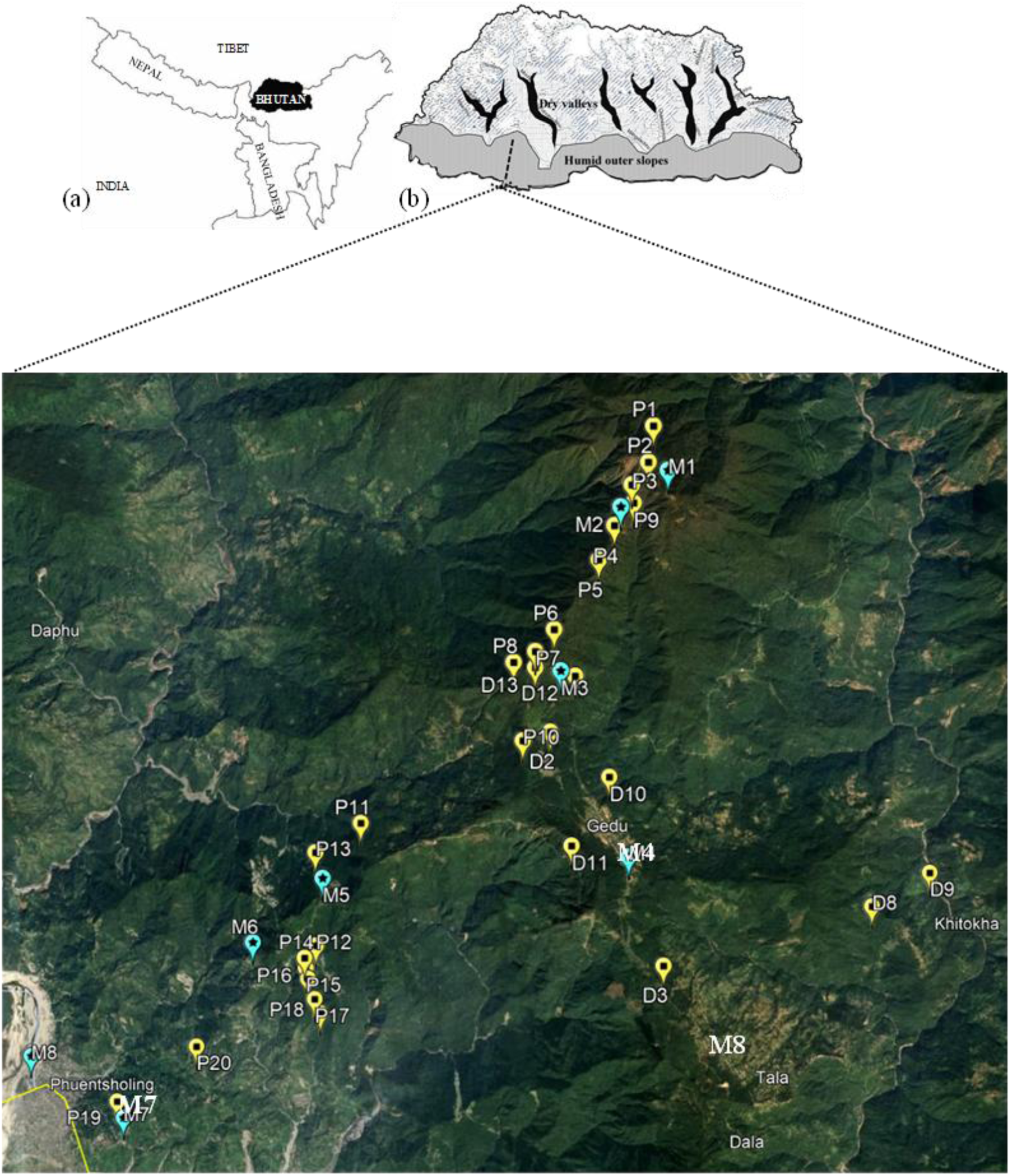

### Appendix S3: Section S1

#### Notes on field methods, sampling, and botanical identification

##### Dates of study

Fieldwork was carried out by Pema Wangda from December 2007 to February 2008, October 2010, June 2011, May 2012 and March 2015, under the guidance of Ohsawa Masahiko. Details on annual incidents of frost events were obtained by Pema Wangda from revisiting long-term data loggers, as well as the Paro airfield, for years spanning from 2009 to 2017. Additional visitation of the Gedu study area by Peter S. Ashton occurred in 2008. Statistical analyses performed by Melissa Whitman in 2025 and 2026.

##### Additional details on field methods used

1. Pema Wangda: Mountain Forest Pattern and Management of Natural Resources in the Bhutan Himalaya. PhD. 2006. University of Tokyo, Japan.
2. Ohsawa, M., 1984. Differentiation of vegetation zones and species strategies in the subalpine region of Mt. Fuji. Vegetatio 57, 15–52.

##### Plant samples were identified using the following literature

1. Brandis, D., 1907. Indian Trees. An account of the trees, shrubs, woody climbers, bamboos and palms indigenous or commonly cultivated in the British Indian Empire. Constable, London.
2. Hara, H., 1968. Photo-album of plants of Eastern Himalaya. Tokyo: Inoue Book Co.
3. Gardner, S., P. Sidisunthorn, Anusarnsunthorn, V., 2007. Field guide to Forest Trees of Northern Thailand. Kobfai Publishing Project, Bangkok, Thailand.
4. Grierson, A.J.C., Long DG., 1983-2000. Flora of Bhutan Vol. I. Part 1,2,3. Vol. II. Part 1,2,3 & Vol. III. Part 1,2. Edinburgh, UK.
5. Nakao S., Nishioka K., 1984. Flowers of Bhutan. Tokyo: Asahi Shimbun Publishing Co.
6. Noltie, H.J., 1994-2000. Flora of Bhutan: Part 1 & 2, vol. 3. Royal Botanic Garden, Edinburgh.
7. Polunin O., Stainton A., 1984. Flowers of the Himalaya. Oxford Univ. Press. New Delhi.
8. Pradhan, R. 1998. Wild Rhododendrons of Bhutan. Quality Printers, Kathmandu.
9. Stainton, A., 1988. Flowers of the Himalaya. A supp. Oxford Univ. Press. New Delhi.
10. Zhengyi, W., Raven, P.H., Deyuan, H. (editors), 1994-2013. Flora of China. Published by the Missouri Botanical Garden, St. Louis.

##### Nomenclature updates or synonyms

1. Boyle, B.L., Matasci, N., Mozzherin, D., Rees, T., Barbosa, G.C., Kumar Sajja, R., Enquist, B.J., 2021. Taxonomic Name Resolution Service, version 5.0.2 [WWW Document]. Botanical Information and Ecology Network. https://tnrs.biendata.org/. URL https://tnrs.biendata.org/ (accessed 8.3.21).
2. Plants of the World Online. Facilitated by the Royal Botanic Gardens, Kew. Published on the Internet; https://powo.science.kew.org/ Retrieved 14 May 2025.’

##### Conifer GBIF query

1. Records of *Abies densa* to estimate range of potential occurrence. GBIF.org (14 February 2026) GBIF Occurrence Download https://doi.org/10.15468/dl.vfqfhc

### Appendix S4: Section S1

#### Notes on meteorological sampling Equipment

For meteorological data, used for subsequent predictive models of climate conditions across sites, we installed HOBO Onset data loggers (Onset Computer Co. MA, USA), that were placed 1.3 m above the ground and enclosed in solar radiation shields. Along an altitudinal transect from 210 to 3370 m a.s.l., we recorded temperature (°C) and relative humidity (%) at one-hour intervals using eight data loggers. We downloaded data every three to six months using a data collector (Boxcar Pro for Windows, Version 4.3 Onset Computer Co.) and summarized information as monthly averages, and outlier events noted as absolute minimum and maximum conditions. Across sites we calculated lapse rate, defined as the linear decline in temperature (°C) per 100 m of altitude increase, which represents the slope of average temperature as a function of altitude using linear regression, calculated per month and annually. For rainfall metrics we calculated annual precipitation (PPT) in millimeters (mm), based on data collected from tipping-bucket rain gauges (HOBO Pendant Event Data Logger, RG3-M) with 0.2 mm resolution. These were installed at three locations (210, 1750, and 2000 m a.s.l). We supplemented our dataset with additional PPT data summarized by Dorji et al. (2016), selecting sites with the highest precipitation per altitudinal bin, spaced every 400 m spanning from sea-level to 4000 m a.s.l.. For instances where stations overlapped spatially with our transect range gauges (i.e. Phuntsholing and Tala) we averaged precipitation values across sites, yielding information on a total of 15 sites.

On a monthly basis, the warmest temperature recorded was 30.1 °C in September (at 210 m a.s.l.), while the coldest -5.5 °C was in January (at 3370 m a.s.l.). Monsoon events, which occurred from June to August, account for 60-65% of the total rainfall. During this period, mid-altitudinal areas experienced maximum relative humidity (RH) values up to 100%. The dry season spanned the winter months, with RH values typically ∼ 70%, but dropping to 13.7% on the mountain summit in December. Mean annual rainfall decreased with increasing altitude, with 4075.2 mm recorded at Phuntsholing (210 m a.s.l.) compared to 3773.0 mm at Darla (1750 m a.s.l.) and 3412.4 mm at Gedu (2000 m a.s.l.). See additional file for monthly data (dataset 1, raw_climate_monthly.csv) for details.

Overall, we found a strong linear relationship between altitude and temperature, with an annual mean lapse rate of 0.59 °C / 100 m^-1^. Average monthly lapse rates also changed seasonally, depending on the RH levels. The lowest lapse rate value (0.49 °C ·100 m^-1^) occurred in July, during the wet monsoon season, and the highest values occurred in March (0.71 °C ·100 m^-1^). Higher humidity during the monsoon season is associated with a reduced lapse rate, buffering temperature extremes along the gradient. See figure on the next page.

### Appendix S4: Figure S2

#### Monthly relative humidity as compared to lapse rate, which is the decrease in temperature (°C) per 100 m increase in altitude, along the moist slope transect

The grey ribbon represents maximum to minimum relative humidity (RH) values, which is highest during monsoon season (May to September), with an inverse trend noted for the monthly lapse rate.

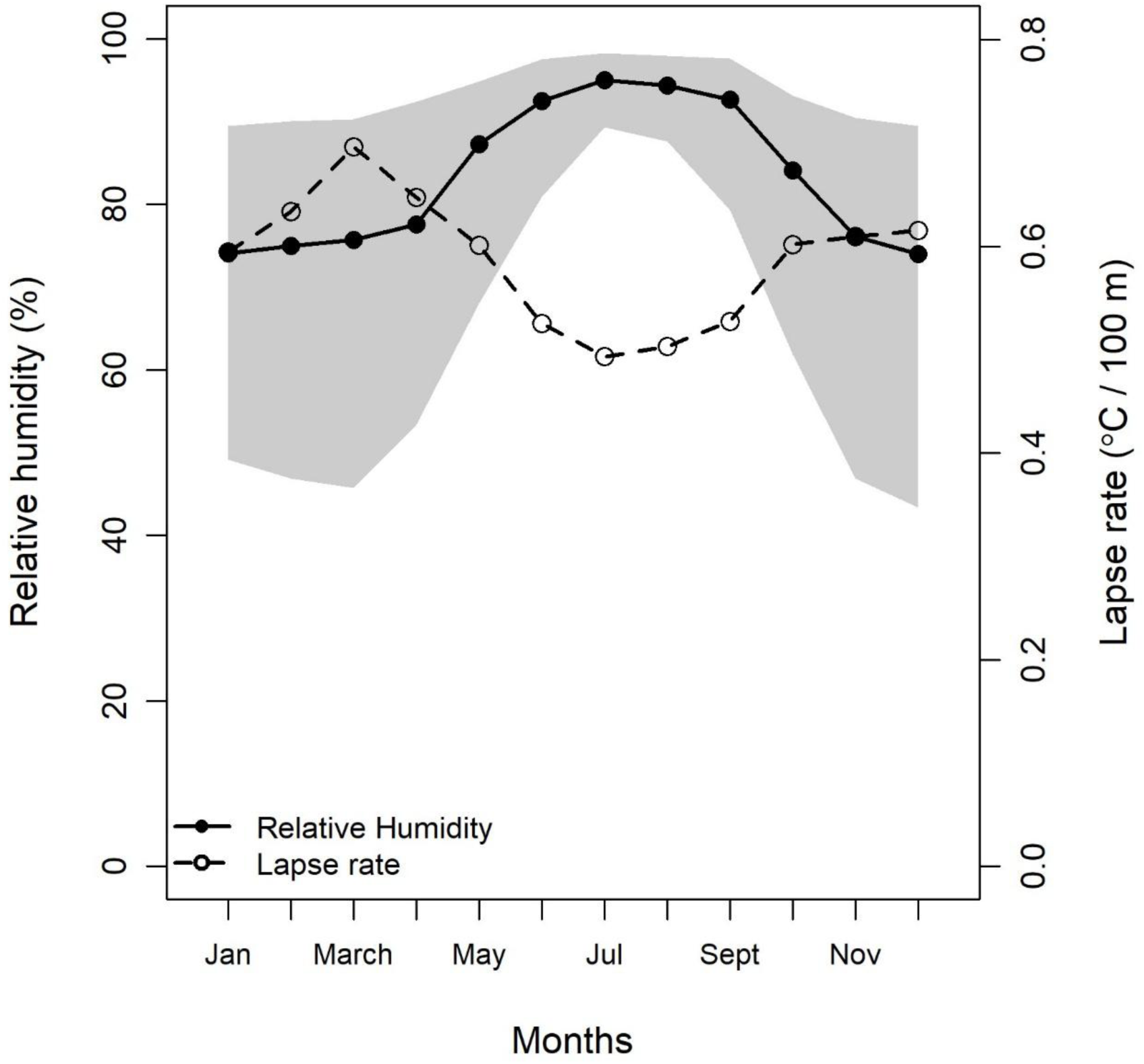

### Appendix S5: Table S1

#### Minimum January temperatures (°C)

Table to illustrate the occurrence of unusual frost events (shown in bold) at lower altitudes, collected based on information available at select locations and across years. Highlighted rows are the lowest altitudes (by moist or dry slope sites) with frost events, with moist slope being areas covered by this study, dry slope being areas included in study by Wangda and Ohsawa, 2006a, 2006b.

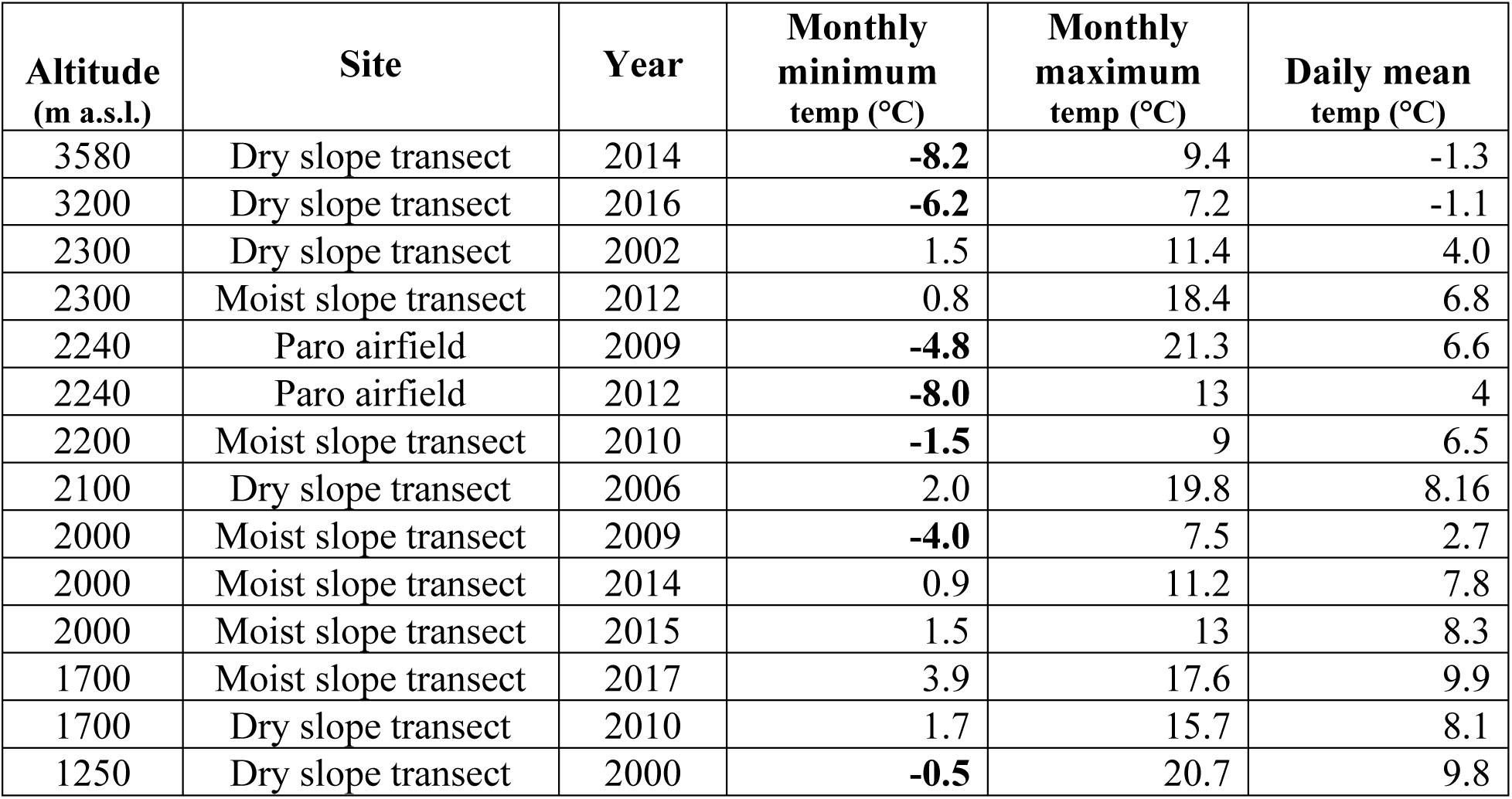

### Appendix S5: Section S2

#### Probability of stochastic frost events

Table and figure shows the probability of a stochastic frost event across the altitude gradient, based on minimum temperatures of December, January, and February, estimated using bootstrap resampling of 1000 posterior draws (1000 resamples). The dashed horizontal line marks the 50% probability threshold (∼2534 m a.s.l.). The dotted vertical line marks the lowest altitude with any predicted frost occurrence (1500 m a.s.l., 0.2%). Frost probability is essentially zero below 1400 m and approaches near certainty above 2800 m a.s.l..

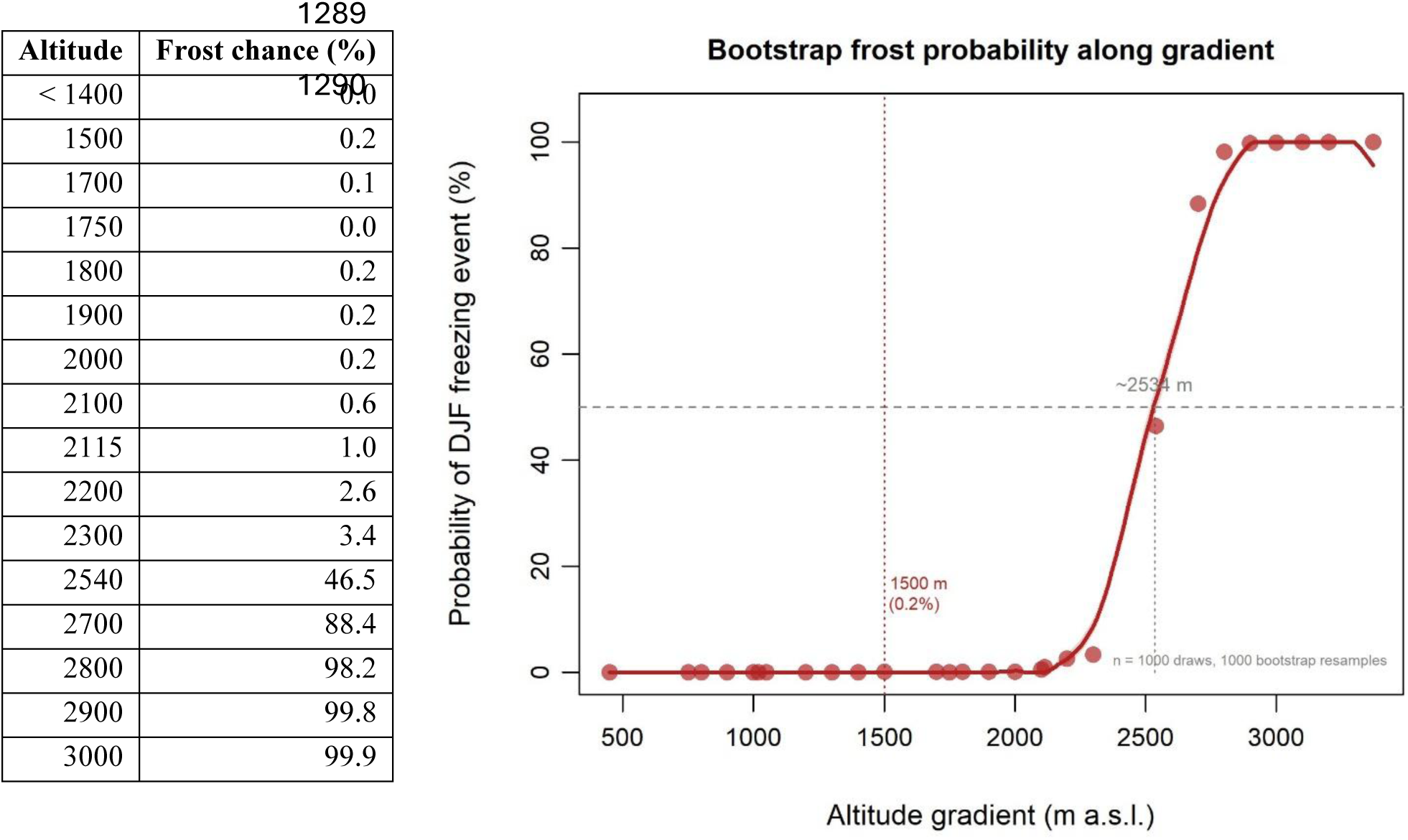

### Appendix S6: Sections S1-S7

#### Overview of climate predictors

We summarized the distributions of each of the six climate predictors, each representing 1,000 posterior draws from their respective interpolation model used to estimate conditions for the 28 forest survey sites, shown in tables and figures. Faint lines represent individual posterior draws (illustrating every draw), with the bold line representing the pooled mean; the outer dashed lines represent the 95% credible interval (2.5% and 97.5% percentiles). See additional methods notes regarding the original meteorological stations and model selection process. Community analyses follow, including distance-based redundancy analysis (dbRDA) and variance partitioning, which validated the explanatory power of these six predictors beyond altitude alone.

#### Here is the list of climate predictors

a. Freeze months. The estimated (integer) count of months where minimum temperature ≤ 0°C, with most values increasing sharply above ∼1800 m.
b. Vapor pressure deficit (VPD) “stop” count. The estimated count of months with the highest aridity stress (the combination of maximum temperatures and minimum humidity, referred to as “noon”), defined as VPD ≥ 1.5 kPa, a threshold associated with stomatal closure, abbreviated as “stop months”. VPD values are the highest at low altitudes during the early spring months.
c. Fog gap at “dawn” in the fall months. This metric refers to minimum dew point depression, the difference between air temperatures and dew point, which we refer to as “fog gap” at “dawn” indicating conditions where temperature is at a minimum and relative humidity is at a maximum, typically occurring predawn. This specific climate predictor represents the fall season, September to November (SON) prior to the dry winter. Lower values indicate areas with closer proximity to cloud immersion (typically noted mid-altitude), with values at zero indicating fog or mist. While negative values are physically implausible, we regard them as model error range and indicative of high certainty that the air is saturated and values are shown for transparency.
d. Warmth index, extreme annual minimum. This is a metric that represents the thermal energy floor for plant growth, which typically declines with increasing altitude. This metric can be used as an indicator of parts of the gradient that can be subject to stochastic frost events, with the possibility of occurrence as low as ∼2000 m a.s.l.
e. Fog gap CV annual. The coefficient of variation of dawn fog gap across 12 months. This metric peaks in the transitional cloud zone (∼2000–2500 m) where immersion is seasonal rather than permanent, with lower values in areas that are consistently beyond the cloud layer.
f. Relative humidity (%) monsoon range. This metric represents the difference of conditions when comparing the hot summer monsoon months (July-August) minus the dry winter season (Dec-Feb). The range in conditions is highest within alpine areas, and there is a dip at ∼1300 m a.s.l., possibly indicating the rain shadow on the leeward side, caused by orographic lifting.

### Appendix S6: Section S1

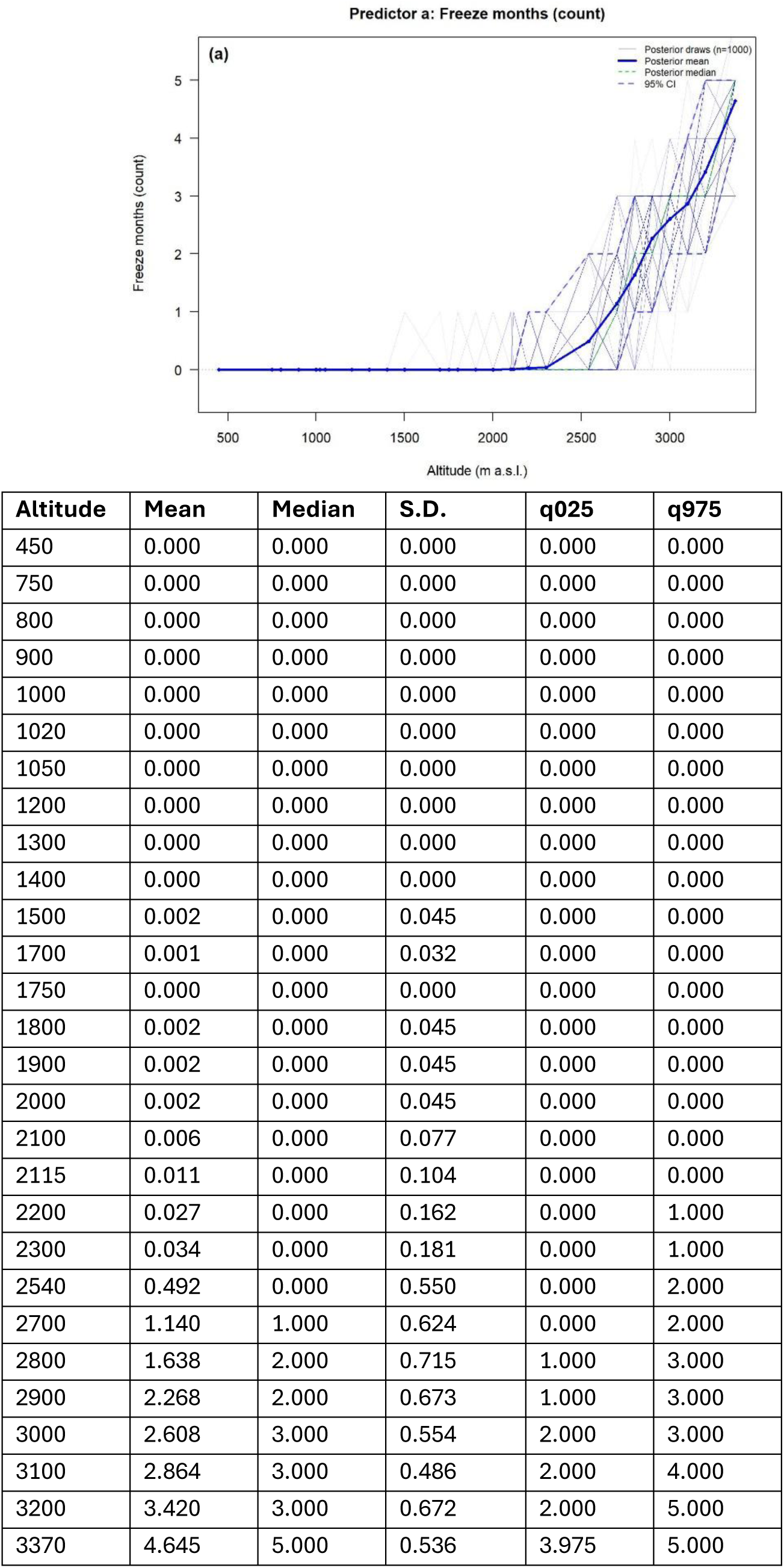

### Appendix S6: Section S2

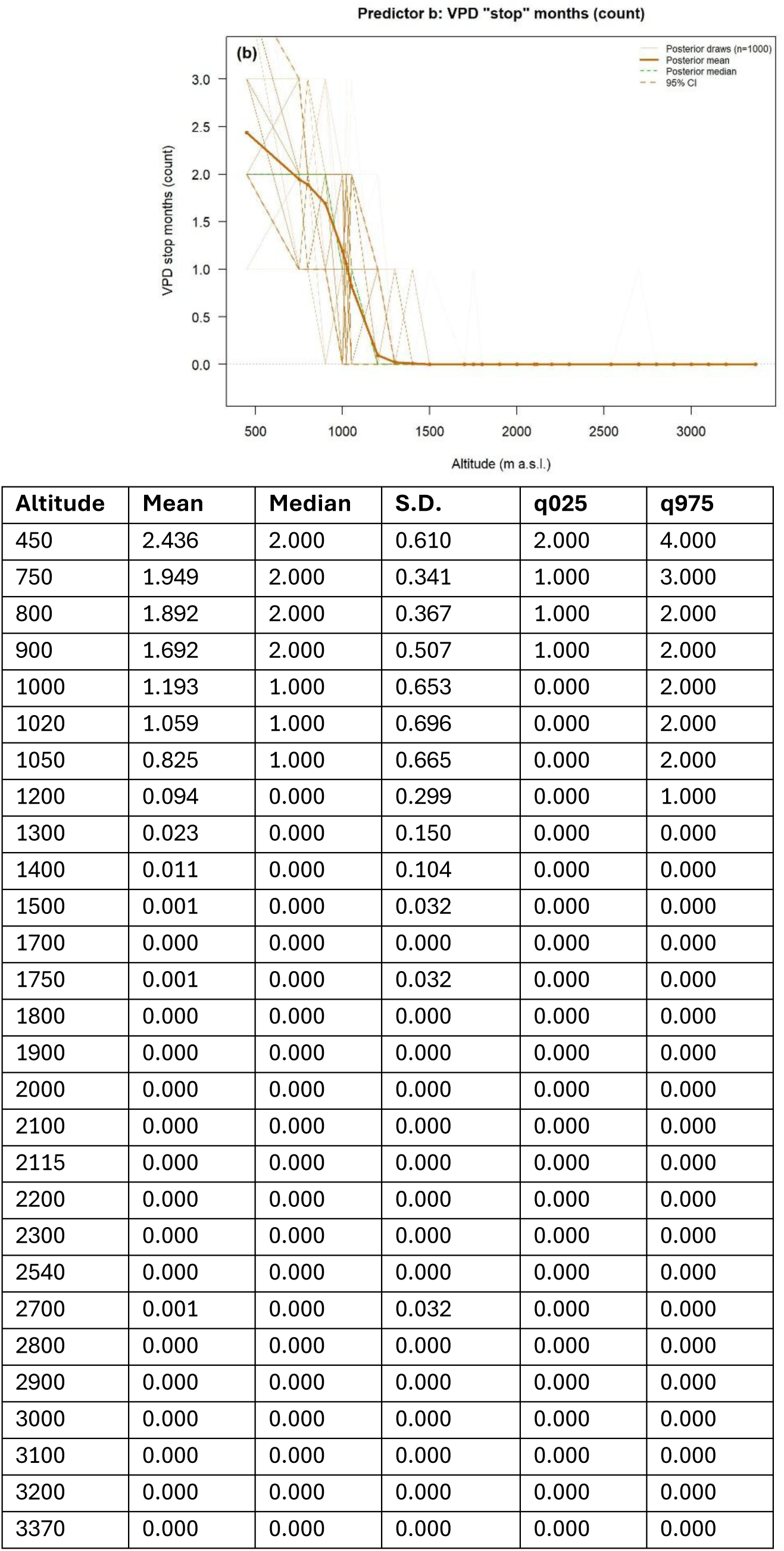

### Appendix S6: Section S3

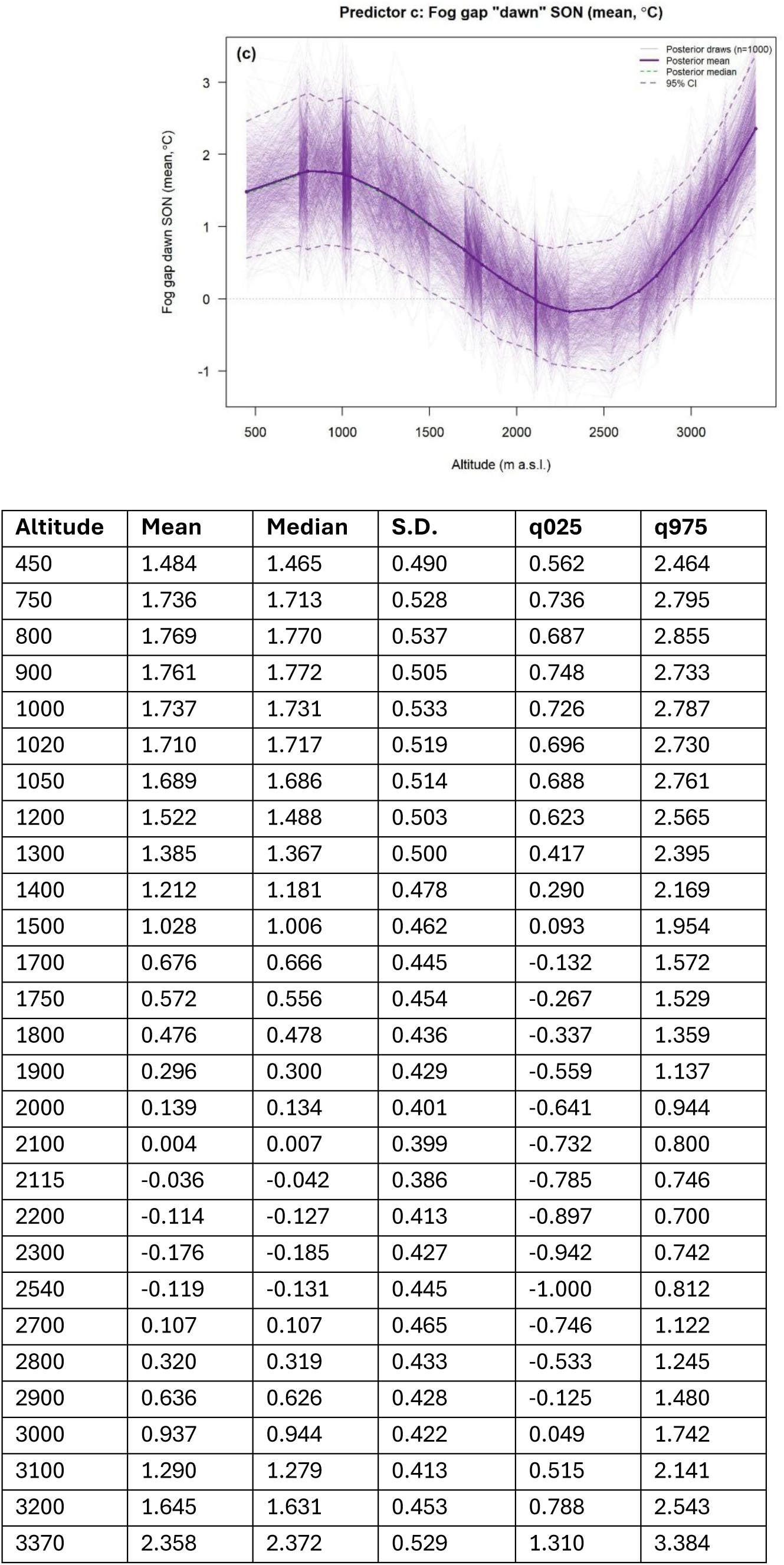

### Appendix S6: Section S4

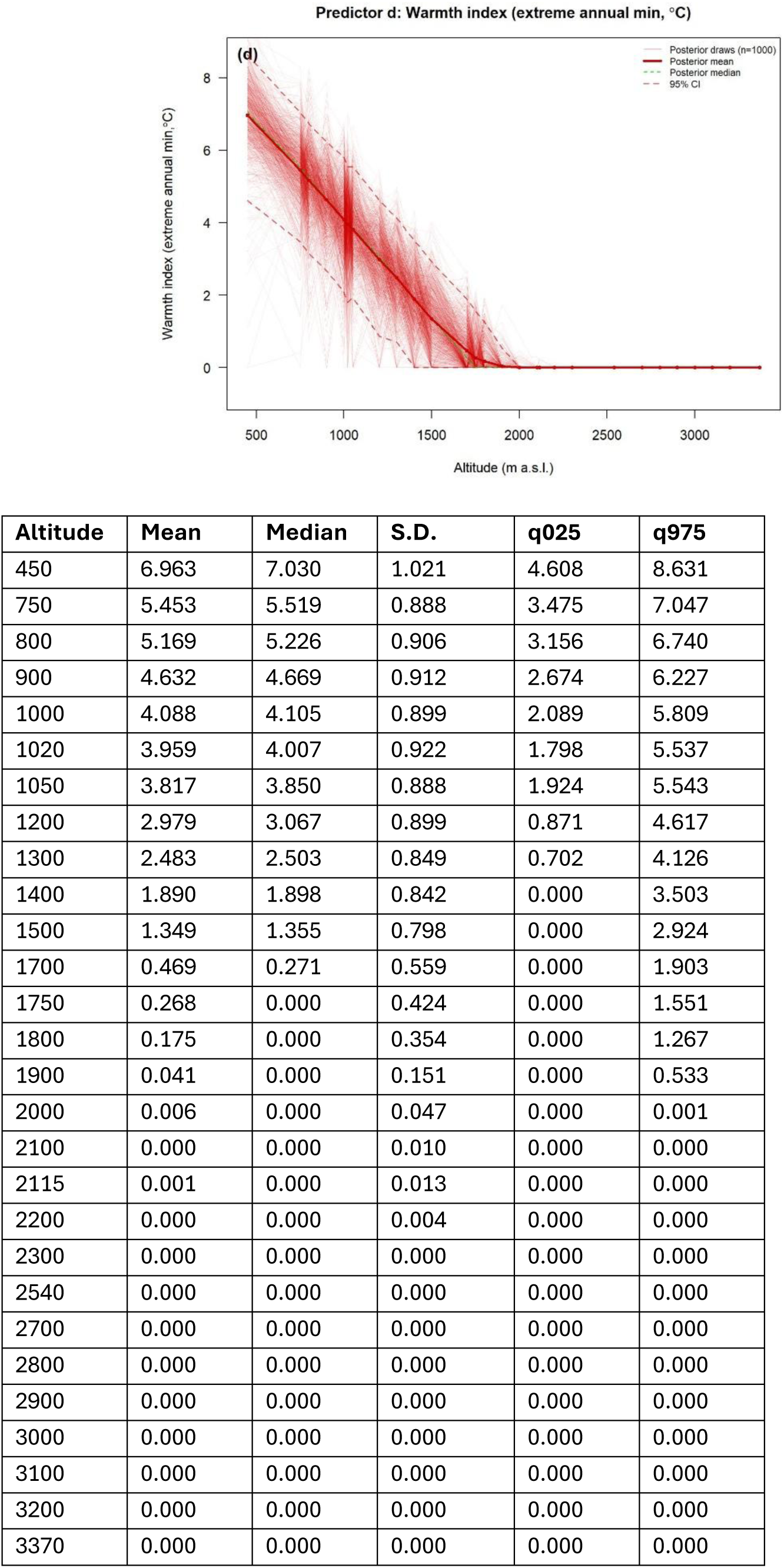

### Appendix S6: Section S5

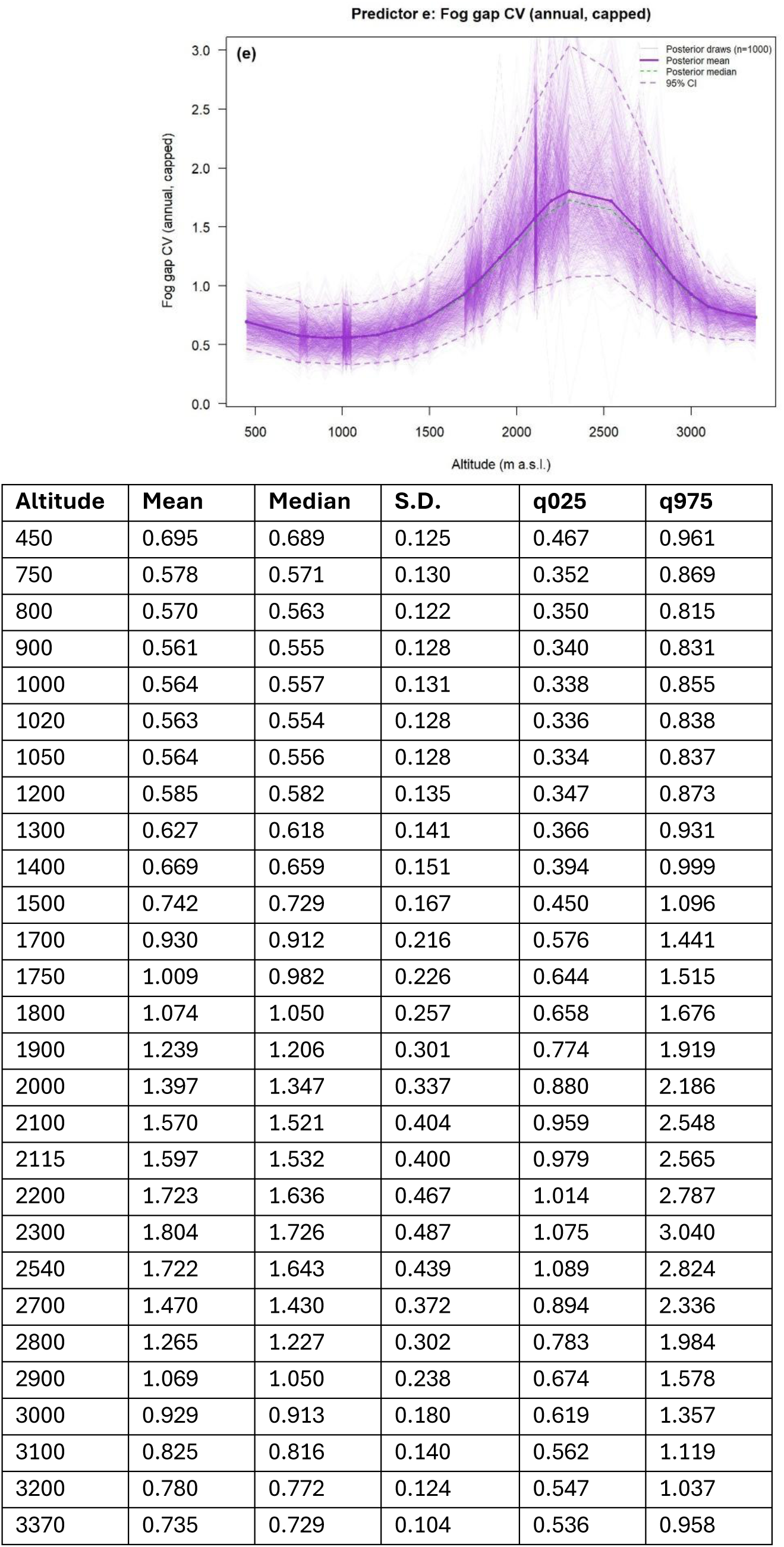

### Appendix S6: Section S6

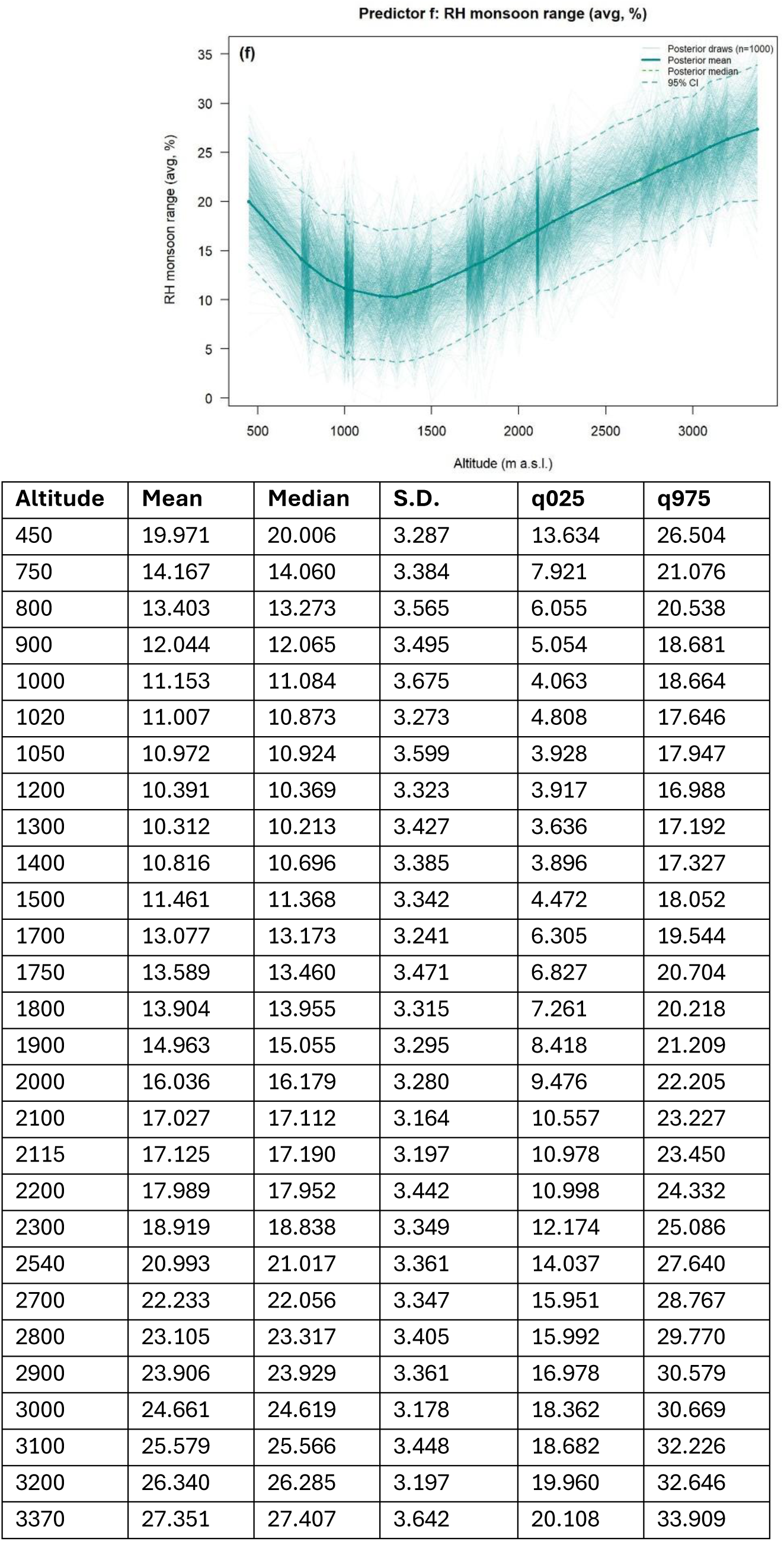

### Appendix S6: Section S7

#### Overview of climate predictors and community analysis

Marginal permutation tests from distance-based redundancy analysis (dbRDA) of four growth form groups across 28 sites. Each predictor was tested for its contribution to explained variation, after accounting for all other predictors. We found that all six climate predictors were individually significant, supporting their retention in subsequent ecological models (after also avoiding high collinearity pairing). *P*-values were based on permutation tests (*n* = 9999), comparing observed versus random metrics of Bray-Curtis dissimilarity using relative basal area as the response.

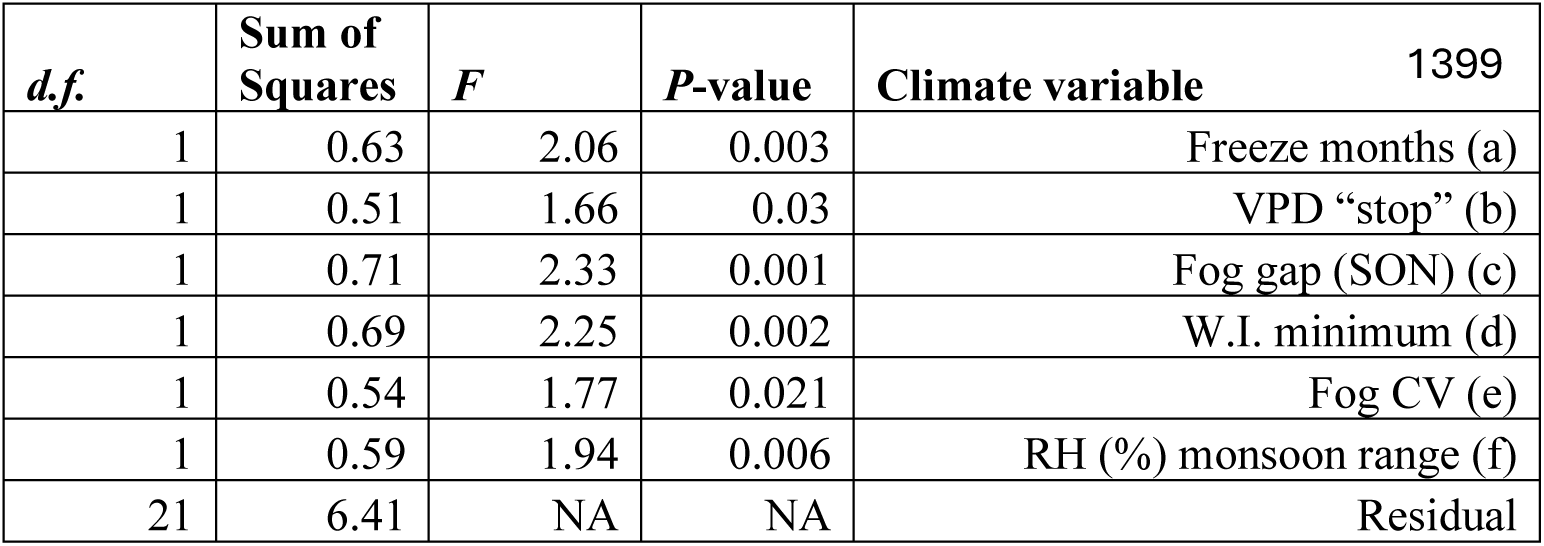

### Appendix S7: Table S1

#### Beta-regression model coefficients

Beta-regression model coefficients *(n* = 1,000 posterior draws per model), pooled via Rubin’s rules. Table includes observed relative basal area (RBA) relationships with climate, as compared to spatially randomized null models. The expected sign indicates the hypothesized direction of each predictor’s effect. Pooled means and 95% credible intervals (C.I.) are shown for both observed and null models. Credible intervals can be used to determine how different observed values are as compared to null models. Column "% agree" is the percentage of paired draws where the observed-minus-null difference was in the expected direction. P-values are based on a one-tailed permutation (proportion of deltas in the opposite direction). Support summarizes evidence strength: *** = full CI separation with expected direction; ** = partial separation; * = expected direction with 2:95% paired agreement; N.S. = non-significant.

| Growth form | Predictor | Hypothesized relationship | Real Mean | Real C.I. (lower) | Real C.I. (upper) | Null Mean | Null C.I. (lower) | Null C.I. (upper) | % Agree | P-value | Support |
| --- | --- | --- | --- | --- | --- | --- | --- | --- | --- | --- | --- |
| Tropical deciduous | Intercept |  | -3.25 | -3.99 | -2.51 | -2.96 | -3.69 | -2.24 | NA | NA | --- |
| Tropical deciduous | VPD stop | (+) | 0.38 | -0.15 | 0.9 | 0 | -0.63 | 0.62 | 92.1 | 0.079 | N.S. |
| Evergreen broadleaf | Intercept |  | 2.07 | 0.59 | 3.56 | 1.12 | 0.37 | 1.87 | NA | NA | --- |
| Evergreen broadleaf | Fog CV | (+) | 0.63 | -0.01 | 1.28 | 0 | -0.67 | 0.66 | 98 | 0.02 | * |
| Evergreen broadleaf | RH monsoon range | (-) | -1.19 | -1.88 | -0.5 | -0.01 | -0.65 | 0.63 | 100 | 0 | ** |
| Temperate deciduous | Intercept |  | -2.12 | -2.91 | -1.33 | -2.05 | -2.94 | -1.15 | NA | NA | --- |
| Temperate deciduous | Freeze months | (+) | 0.61 | 0.03 | 1.2 | -0.02 | -0.71 | 0.68 | 98.2 | 0.018 | * |
| Temperate deciduous | Fog gap SON | (-) | -0.35 | -0.93 | 0.23 | 0.01 | -0.67 | 0.7 | 91.9 | 0.081 | N.S. |
| Conifer | Intercept |  | -1.52 | -3.1 | 0.06 | -1.33 | -2.54 | -0.12 | NA | NA | --- |
| Conifer | Freeze months | (+) | 1.35 | -0.2 | 2.9 | 0 | -1.62 | 1.61 | 95.4 | 0.046 | * |

### Appendix S7: Section S2

#### Beta-regression, observed as compared to null distribution figures

Observed versus null coefficient distributions for relative basal area (RBA) as a function of climate relationships, replicated by growth form and analyzed using Bayesian beta-regression. The figures illustrate the distribution of the standardized regression coefficients (from the top model for each growth form, with one or two climate predictors used), as calculated from 1,000 posterior draws, with colored points representing observed values as compared to null values, which are shown in grey. Climate interpolation uncertainty was propagated by fitting each draw independently, then pooling results via Rubin’s rules. *A priori*, we hypothesized the directional response of each response and predictor pairing, as indicated as (+ or -), with these predictions we tested the paired delta methods (real minus null values per draw), where the *P*-value is the proportion of deltas in the opposite direction of the hypothesized effect. Filled circles indicate the pooled mean of each distribution; vertical dashed line marks a value of zero (which means no effect).

The top models for each growth form (as plotted on the next page) are as follows: (a) Tropical deciduous RBA increases with more VPD “stop” months (months with noon VPD ≥ 1.5 kPa), consistent with drought-deciduousness as an adaptive strategy in the seasonally dry lowlands. (b) Evergreen broadleaf RBA increases with fog CV (annual coefficient of variation of “dawn” fog gap), where higher values indicate ephemeral cloud immersion that provides moisture, and where lower values represent areas of the gradient that are consistently cloudless, and (c) RBA for this group decreases with a higher monsoon humidity range, indicating large seasonal swings in moisture availability. (d) Temperate deciduous RBA increases with freeze months, consistent with cold-deciduousness as a frost-avoidance strategy. (e) Temperate deciduous RBA decreases with fog gap values that are farther from zero, indicating drier air farther from saturation. (f) Conifer RBA increases with freeze months, reflecting cold-tolerance at the highest altitudes.

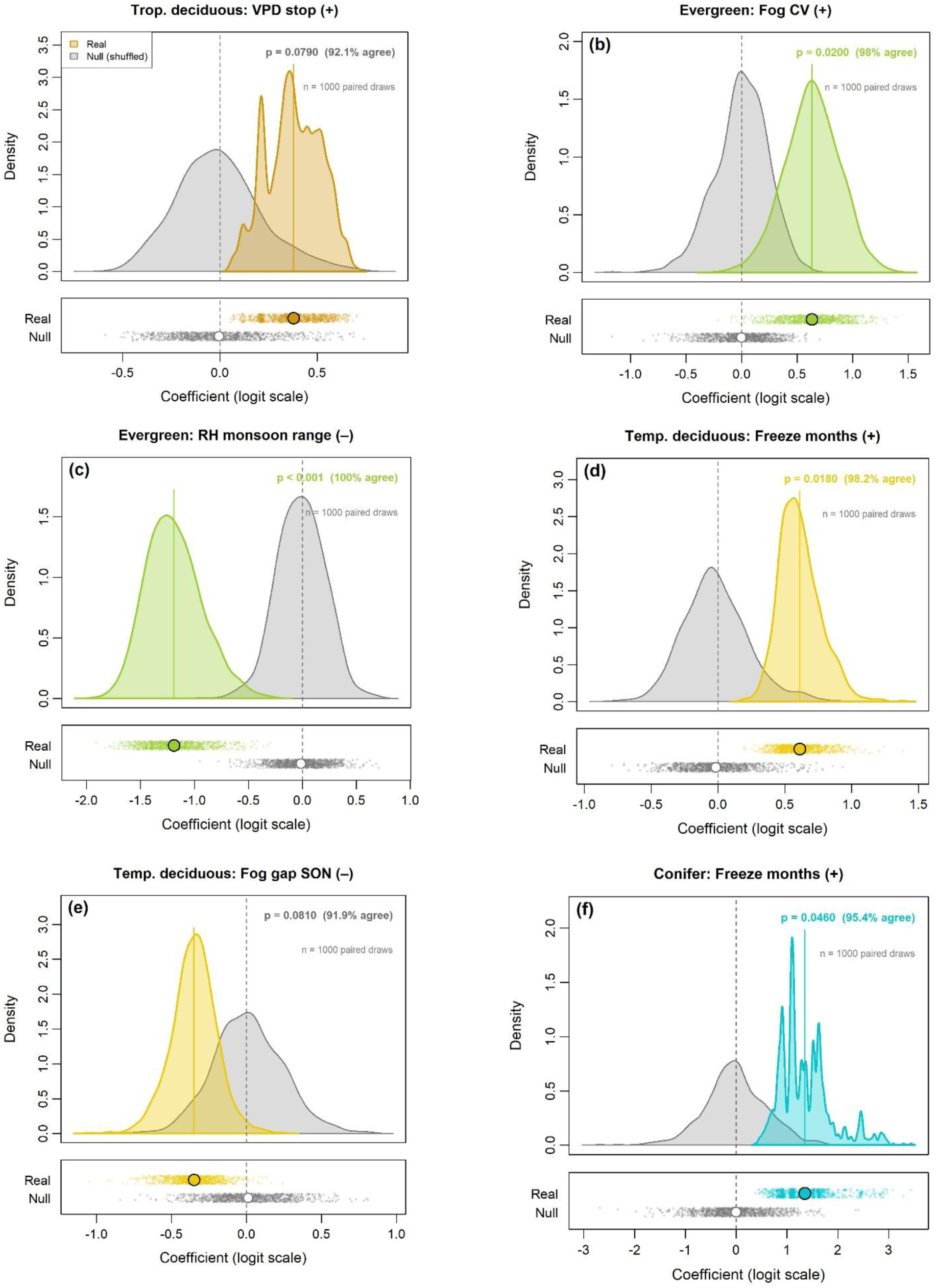

### Appendix S8: Table S1

#### Table of Dirichlet compositional top models, ranked via LOO-IC

These models all used relative basal area (RBA) as the response, representing the composition dynamics between four growth forms (tropical deciduous, evergreen broadleaf, temperate deciduous, conifers), with predictors set either to climatic factors, or altitude in either linear or quadratic form, which is representative of the null model where the ecological mechanism could not be identified. Lower LOO-IC values indicate better predictive performance. The top climate model (a*c, freeze × fog interaction) outperformed both altitude-only null models by a delta LOO-IC of 7.8–10.6. All results are pooled across 1,000 posterior draws via Rubin’s rules. Predictor codes: a = freeze months, b = VPD stop months, c = fog gap SON, d = warmth index min, e = fog CV, f = RH monsoon range, with “*” as an interaction and “+” as an additive term.

| Rank | Model | LOO-IC | Delta LOO-IC | Note |
| --- | --- | --- | --- | --- |
| 1 | a*c | -651.13 | 0.00 | Top model |
| 2 | a*e | -648.14 | 2.99 |  |
| 3 | a*c + b | -647.99 | 3.14 |  |
| 4 | a + c | -646.76 | 4.36 |  |
| 5 | a*c + a*f + d*f | -646.31 | 4.82 |  |
| 6 | a*c + b + d | -645.85 | 5.27 |  |
| 7 | a*f | -645.82 | 5.31 |  |
| 8 | a + c + d | -644.93 | 6.20 |  |
| 9 | a + c + e | -644.92 | 6.20 |  |
| 10 | a*f + d*f | -644.00 | 7.13 |  |
| 11 | altitude (quadratic) | -642.47 | 8.65 | Null 2 |
| 12 | altitude (linear) | -639.86 | 11.27 | Null 1 |

### Appendix S8: Figure S2

#### The top Dirichlet regression compositional model

Illustration of growth form RBA predictions from the top Dirichlet model (with lines indicating per-draw values), which includes interacting climate predictors (count of freezing months x fog gap values for the fall months). For each of 1,000 posterior draws, site-specific climate values were extracted from the interpolated posteriors, globally scaled, and combined with the corresponding fixed-effect coefficients from that draw to compute predicted RBA via softmax transformation.

Each faint line represents a single posterior draw; bold lines show the pooled mean across all draws; filled circles indicate observed RBA at the 28 survey sites. Colors are as follows the four growth form groups: tropical deciduous (dark goldenrod), evergreen broadleaf (green), temperate deciduous (yellow), and conifer (cyan). The “spaghetti” spread reflects the combined uncertainty from both climate interpolation and model coefficient estimation, propagated simultaneously through the prediction. The four growth form groups show distinct altitudinal distributions, with tropical deciduous species concentrated at the lowest sites, evergreen broadleaf dominating most of the gradient, and temperate deciduous and conifer groups increasing at higher altitudes.

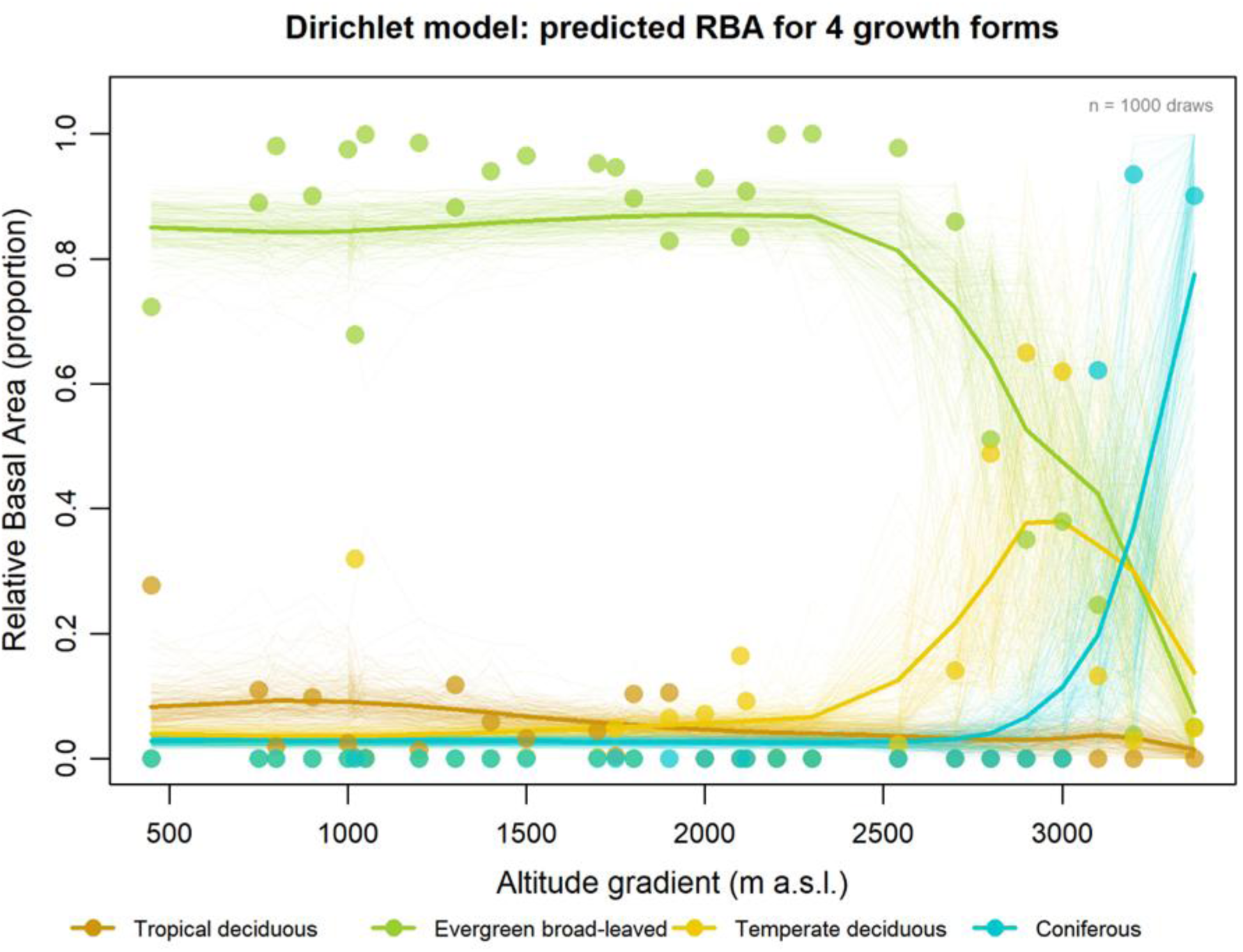

### Appendix S8: Table S3

#### Pooled coefficients from the top Dirichlet compositional model

Top model (freeze months × fog gap interaction), estimated across 1,000 posterior draws and pooled via Rubin’s rules. All coefficients are on the log-ratio scale, relative to evergreen broadleaf growth form group at the reference altitude (1850 m a.s.l.), with negative intercepts indicating a lower expected RBA proportion relative to this point of the gradient. Significance (***) indicates the 95% credible interval excludes zero.

| Growth form | Parameter | Mean | SE | C.I. lower | C.I. upper | Sig. |
| --- | --- | --- | --- | --- | --- | --- |
| Tropical deciduous | Intercept | -2.68 | 0.34 | -3.35 | -2.01 | *** |
| Tropical deciduous | Freeze (a) | -0.10 | 0.33 | -0.75 | 0.55 |  |
| Tropical deciduous | Fog (c) | 0.48 | 0.27 | -0.05 | 1.00 |  |
| Tropical deciduous | Freeze x Fog (a*c) | 0.18 | 0.34 | -0.48 | 0.84 |  |
| Temperate deciduous | Intercept | -2.27 | 0.31 | -2.88 | -1.66 | *** |
| Temperate deciduous | Freeze (a) | 1.35 | 0.39 | 0.60 | 2.11 | *** |
| Temperate deciduous | Fog (c) | -0.35 | 0.28 | -0.91 | 0.20 |  |
| Temperate deciduous | Freeze x Fog (a*c) | -0.24 | 0.36 | -0.95 | 0.46 |  |
| Conifer | Intercept | -3.02 | 0.37 | -3.73 | -2.30 | *** |
| Conifer | Freeze (a) | 0.85 | 0.41 | 0.04 | 1.65 | *** |
| Conifer | Fog (c) | 0.38 | 0.29 | -0.19 | 0.95 |  |
| Conifer | Freeze x Fog (a*c) | 0.63 | 0.42 | -0.20 | 1.46 |  |

### Appendix S8: Figure S4

#### Predicted relative basal area (RBA) under different climate scenarios, based on the top model selected (freezing x fog interaction)

Each scenario (± 1-2 SD for each variable) estimated via the multinomial logit link function (aka softmax), with each cell representing the predicted RBA (%) for a given combination and the departure from reference conditions of the gradient median (set to 1850 m a.s.l., with 0 SD for both predictors, indicated by the black box). All four panels sum to 100% at each cell, and coefficients were pooled across 1,000 posterior draws via Rubin’s rules. Negative fog gap values indicate conditions closer to atmospheric saturation (more cloud immersion), while positive values indicate drier air farther from the cloud layer. The interaction between frost and fog is visible in the opposing responses of temperate deciduous species (which peak under cold, foggy conditions) and conifers (which peak under cold, dry conditions), illustrating how moisture context determines the compositional outcome of frost exposure.

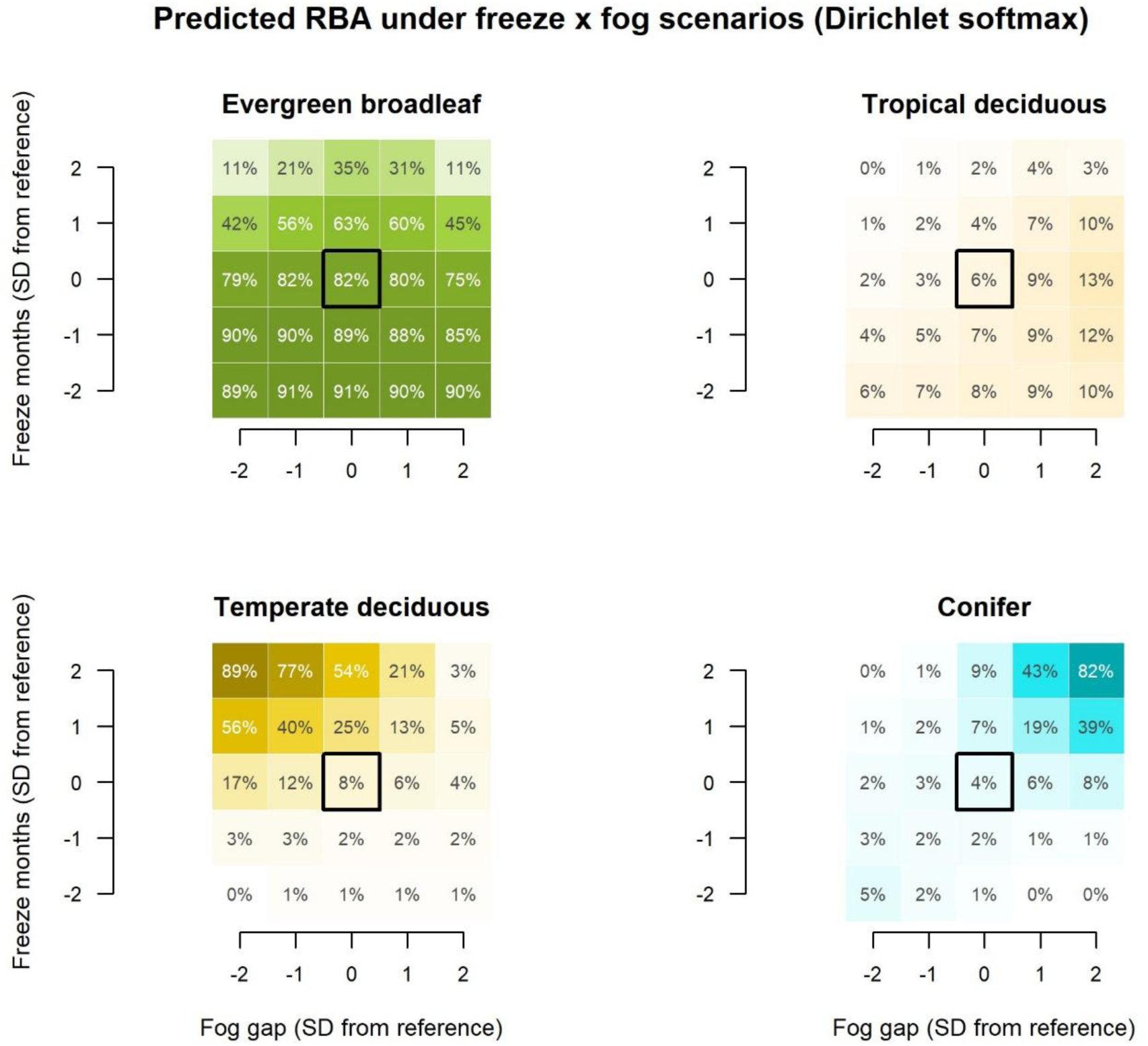

### Appendix S9: Section S1

#### Additional notes on Bayesian Modeling and overarching philosophy

The following notes are provided to aid in the transparency and reproducibility of the statistical methods used in this study. They outline various details that there simply was not space for within the main manuscript, encompassing nuances such as the logic of why various decisions were made, the protocol applied within and across the modeling pipeline, and miscellaneous technical details that may be of interest to a subset of readers (e.g., software and hardware used, modeling parameters, quality control procedures).

The overall Bayesian models used in this research fall into two camps of thought. The first modeling set is predictive, with the goal of climate interpolation, which at its heart is more of an exploratory process to understand conditions along the altitudinal gradient. This first modeling set involved daydreaming about how clouds work when meeting the mountain side or being shaped by sometimes conflicting forces of temperature and humidity, then seeing if said ephemeral forms could take shape via the numbers. In total, more than six hundred climate predictions were made, but only few (the select ∼1%) made it into our final modeling round, with prioritization informed by the thoughtful musings of those with decades of field observations. This then led us to the second Bayesian model set, which is distinctly inference based, to test ideas regarding the mechanisms that shape forest composition and growth forms.

Throughout the Bayesian modeling process, we prioritized the tracking of uncertainty, favoring a slower but more deliberate process in which each individual step and output could be evaluated independently. By decomposing the analysis into discrete, auditable stages (i.e., interpolation, derivation, aggregation, metric calculation, and ecological modeling) we ensured that intermediate outputs could be inspected, validated against field knowledge, and traced back to their source data. This modular approach was chosen over alternatives such as Gaussian process regression or joint hierarchical models that estimate climate and ecological responses simultaneously, where the propagation of uncertainty may be handled internally, but at the cost of less scrutiny or accessibility to the intermediate stages. While such approaches are statistically valid, the staged pipeline that we opted for makes every analytical decision visible and defensible, albeit after considerable computation time and tedious notetaking.

Our Bayesian framework was adopted after an extensive preliminary analysis using ordinary least squares (OLS) and weighted least squares (WLS) regression, which are best described as unsatisfactory. The original OLS/WLS models had more of a data-mining approach, produced upwards of 14,000 univariate models, using a broader set of response variable and easily compiled predictors. While many of these models were significant, they simply did not tell an interesting story; we were also dissatisfied with the collinearity among derived climate predictors, and substantial underestimation of error for derivatives or aggregates. Simple models produced simple results, failing to capture the essence of a system supporting both tropical and temperate assemblages. These concerns motivated us to do a philosophical pivot in real time, embracing a Bayesian approach centered on two principles: hypothesis-driven model design and the propagation of uncertainty as the central premise underlying all analytical choices to differentiate the known from the unknown. Thus, the Bayesian methods that we used were developed through experimentation and necessity, rather than following a pre-existing template. Many of the methods were further refined via the observation and correction of errors, with many small victories eventually culminating to become the study as it is today. The following notes are intended to help with lingering questions, or for replication of modeling methods used.

#### Bayesian climate interpolation and derivation

We processed climate data as part of a four-stage pipeline that transforms raw station measurements into ecologically meaningful posterior distributions, predicted for each of the 28 forest survey sites. This pipeline is strictly unidirectional, each stage reads only from the outputs of the preceding stage and never modifies earlier files, ensuring auditability of the steps taken.

Our set of climate models included many minor adjustments. First is that we set the maximum treedepth parameter to 12, rather than the Stan default of 10, which allowed the sampler more room to explore complex posteriors (i.e. wander freely) at the slight cost of additional computation time. Similarly, we set the target average proposal acceptance probability (adapt_delta) to 0.95 rather than the default of 0.80, causing the sampler to take smaller, more cautious steps during the leapfrog integration, as is recommended when dealing with complex or correlated posteriors and in turn reduces the occurrence of divergent transitions.

#### Reproducibility

We use a fixed random seed (i.e. 12345) for all associated scripts in R, to ensure reproducibility of results. To verify that findings were not sensitive to seed choice, we ran the full pipeline twice, once using a randomly selected seed and once using a set seed, with consistent overall results. In both instances, the same top model was selected, coefficient estimates varied within rounding precision of the reported values, and no qualitative conclusions changed, which was quite reassuring. All source files and code are available on Zenodo.

#### Stage 1 (Interpolation with model selection)

Monthly temperature and relative humidity from eight data logger locations, summarized to daily average as well as monthly absolute minimum and maximum, were interpolated to the 28 forest survey sites using Bayesian parametric regression. For each of the variable and month combinations (72 in total) we fit four candidate models (i.e., linear, natural spline df = 2, 3, and 4). All candidate model outputs were retained for review if needed. In addition, we also interpolated precipitation, but using a mix of data from three sites (that we collected) and additional sites in Bhutan (from summaries by Dorji et al. 2016), representing information for 15 sites along the gradient, but with resolution at the seasonal, rather than monthly, basis. Each selected model produced 1,000 posterior predictive draws at all 28 target sites.

Regarding the process of deciding on a “best model”, we expected that several climate variables, particularly anything related to relative humidity, would exhibit highly non-linear trends along the gradient, reflecting the complex interplay between cloud positioning, wind patterns, and mountain topography. Rather than using strict parsimony for best model selection (i.e. LOO-IC), we instead opted for a conditional logic-based selection process. We opted for this change after noting that the LOO-IC model selection process was more conservative than Akaike Information Criterion (AIC), when comparing identical models performed using OLS/WLS. Instead, we opted for picking the most complex model “within reason” out of a competitive set (LOO-IC delta < 2 from the best model), selecting for more complex models as long as the ratio of delta LOO-IC to the standard error of the expected log pointwise predictive density difference (SE_diff) was also less than 2. Under this criterion, slightly more complex spline models for climate were often selected during transitional months outside the monsoon season.

#### Stage 2 (Monthly derivatives)

For the next stage, we focused on derivatives based on temperature and humidity, using element-wise matrix calculations that preserved the joint posterior structure (e.g., draw 1 of temperature paired with draw 1 of humidity, etc.). Derived metrics include vapor pressure deficit (VPD), dewpoint temperature, fog gap (temperature depression to dewpoint), biotemperature (ABT), and warmth index (WI). For moisture-sensitive metrics, three diurnal variants were calculated (i.e., "dawn", "avg", and "noon"): "dawn" (temperature minimum × humidity maximum, representing pre-dawn saturation conditions), "avg" (daily averages), and "noon" (temperature maximum × humidity minimum, representing midday aridity). Calculations were performed at the monthly level before temporal aggregation to preserve information about within-month covariance and extreme conditions.

#### Stage 3 (Temporal aggregation)

The goal of stage 3 was temporal aggregation (monthly to seasonal or annual time scales) as well as additional derivatives using precipitation data, that was based on seasonal information. Aggregation types included seasonal means, absolute extremes (i.e., element-wise minimum or maximum across all variants and months within a period), and precipitation-derived metrics including potential evapotranspiration (PET, Hargreaves-Samani method), aridity index, and potential evapotranspiration ratio. Seasonal abbreviations include the cold dry season December to February (DJF), March to May (MAM), the monsoon peak from June to August (JJA), and September to November (SON).

#### Stage 4 (Custom ecological metrics)

We calculated metrics based ecophysiological thresholds (e.g. frost events) or temporal variability of the climate regime, using posteriors that were generated in Stages 1-3. These metrics included threshold-based counts and durations (e.g., months below freezing, months with VPD above stomatal closure thresholds), annual coefficient of variation (inter-monthly climate stability), monsoon range (JJA minus DJF amplitude), and intra-seasonal range (within-season variability across diurnal variants).

#### Consolidated modeling archive and naming convention

In total, the pipeline produced >600 posterior files as potential climate predictors. Even though the vast majority were not used, these posteriors were still quite informative, and when plotted they provided perspective on the complexity of the Bhutan’s monsoon climate and the range of conditions along the altitudinal gradient. All posterior files used as potential climate predictors followed a strict five-component naming convention that functions as embedded metadata, encoding the climate variable, variant or diurnal period, temporal resolution, time period, and derived statistic (e.g., fog_dawn_seasonal_son_mean_posterior.csv denotes the mean dawn fog gap in the fall season. This convention ensures that every file is self-documenting and enables programmatic filtering across the library; for example, all fog-related posteriors can be identified by the prefix fog_*, and all seasonal DJF files by the pattern *_seasonal_djf_*. All posterior files from Stages 1–4 were copied to a single flat directory (the modeling archive for big posterior files, named mixalot_data) to serve as the input for Stage 5 ecological models.

#### Bayesian proportional models (beta-regression and Dirichlet)

The next set of Bayesian models both used growth form relative basal area (RBA) as the response, with these proportional values contained within a 0 to 1 space, and climate variables as predictors, but differed in that the beta-regression regarded each growth form individually as separate models whereas the Dirichlet compositional model also captured dynamics between the four groups simultaneously. The beta-regression models also differed in that they included only portions of the gradient relevant to a given response group’s locations of occurrence (or predicted range of potential occurrence in the instance of conifers), avoiding zero-inflation. Gradient subsets were as follows: tropical deciduous (450 to 2200 m a.s.l., 19 sites), evergreen broadleaf (450 to 3370 m a.s.l., 28 sites), temperate deciduous (1020 to 3370 m a.s.l., 23 sites), and conifer (2700 to 3370 m a.s.l., 7 sites, based on GBIF-derived altitudinal range). The Dirichlet model spanned the full 28 plot gradient, from 450 to 3370 m a.s.l.. The same six climate predictors were used for both model sets, but the beta-regression included simpler model design. Similarly, both model sets centered the reference-altitude to median of each respective gradient, such that the intercept can be interpreted as the expected logit-transformed RBA under the climate conditions of that reference location and the coefficient represents the change in logit (RBA) per one standard deviation departure from these reference conditions. One detail of the Dirichlet model, is that because it is proportional and all values must add to 1, sites where a given growth form was absent (RBA = 0) were adjusted by adding a small constant (0.0001) followed by renormalization to sum to 1. For the ecological model set, each model was fitted separately for each of the 1,000 climate posterior draws, with results pooled across draws using Rubin’s rules (Rubin, 1987). This is distinct from the climate interpolation stage, where 1,000 posterior draws are produced directly from a single model fit.

Selection of the best model for both Bayesian sets was a multi-step process. Both model sets included a preliminary decision round of models, each fitted using 10 posterior draws rather than the full 1000. For the beta-regression models, we examined 14 relatively simple models across 4 growth forms, which was then narrowed down to a single full model, whereas the Dirichlet model included a 75-model decision round (encompassing simple to highly complex models) which was narrowed down to a set of 12 full models. For the null model design of the beta-regression set, each production model (RBA of a given growth form ∼ climate) was paired with a null model using identical climate posteriors and model design, but with shuffling of RBA spatial association for each draw. This process preserved the overall distribution of each growth forms’ RBA values, while breaking the spatial co-occurrence between growth form dominance and climate, with separate pooling of coefficients via Rubin’s rules. We assessed the separation between credible intervals for null and observed ranges, and additional comparisons of paired deltas (i.e. values per permutation draw) for the proportion of indices where real coefficients exceeded the null in the hypothesized direction (e.g. larger tropical deciduous RBA in more arid areas with higher VPD values), with p-values based on one-tailed permutations. The null model comparison for the Dirichlet compositional model used a simpler design, being the comparison of the top 10 climate-based models relative to 2 altitude-only models (representing linear or quadratic forms), with best model(s) chosen based on LOO-IC and general assessment of highest ranked factors. The relative rank of the top climate model, as compared to the top altitude model, addresses the question of whether hypothesized relationships have more explanatory power, even after incorporating uncertainty of the climate interpolation process (a.k.a. was it worth the effort).

#### Quality control

Across, all models, we assessed model quality via four diagnostics for each of the 1,000 individual model fits, by evaluating the following: (1) convergence success rate, defined as the proportion of draws where all parameters met diagnostic criteria; 2) maximum R-hat < 1.01 for all parameters, a convergence diagnostic based on the potential scale reduction factor by Gelman and Rubin (1992); 3) an effective sample size (ESS) somewhere in the realm of at least 400, but higher numbers are better, which is done via estimating the number of independent posterior samples after accounting for autocorrelation, calculated using the posterior package and filtered to model parameters to exclude auxiliary parameters with artificially low ESS values; and 4) count the number of divergent transitions and hope it is zero (if not then adjust treedepth, set a higher adapt_delta, or toss the entire model out as being more trouble than worth). We also recorded model run time per draw, which gave us the opportunity to identify fits with unusually slow convergence and was used to justify building an awesome computer to begin with. For each beta-regression model fit, we also extracted the precision parameter (phi) and its standard error, which quantifies the concentration of observed RBA values around the predicted mean, with higher values indicating tighter clustering.

As a quality control step for climate predictions prior to ecological modeling, we conducted both numerical and visual checks on all interpolated values (i.e. does it match what is reasonable to expect for this corner of the globe, based on the knowledge available). Some of the tedious number gazing included numerical checks of posterior mean as compared to median values in wide-format summary files. When we encountered numbers that stuck out like a sore thumb, such as a large mean–median divergence or impossibly huge numbers, we would investigate to determine if it was a calculation error or if it was just the misunderstood offspring of two unusual events occurring. Mildly problematic edge cases, such as negative VPD or fog gap values (which are physically impossible but can happen with matrix derived data) were intentionally retained in early pipeline stages, more as diagnostic indicators of interpolation uncertainty at gradient extremes, and then optionally capped if used as a predictor. We also inspected extreme outlier values, that can indicate errors in the order of operations or computation (e.g. dividing by zero). The visual check of the diagnostic plots were honestly the nicest of the quality control steps, spending time simply looking at monthly and seasonal predictions across the full elevational gradient, with observed station data overlaid for validation. These colorful plots were assessed against natural history expectations and physical constraints. For example, verifying that temperature lapse rates were consistent with known adiabatic gradients (or better yet were similar to numbers recorded via independent studies conducted on nearby mountains), that relative humidity approached but did not systematically exceed 100%, and that seasonal patterns reflected expected monsoon dynamics. Diagnostic plots were additionally reviewed by co-authors with extensive field experience along the gradient (Pema Wangda) or overall deep knowledge of Asian mountains and their forests (Ohsawa Masahiko, Peter S. Ashton) to verify that interpolated climate patterns were consistent with their field observations of conditions across the gradient or by seasons.

#### Model diagnostic summary

Across the full study, a total of 20,994 individual Bayesian model fits were performed, comprising 87.8 million Markov chain Monte Carlo (MCMC) iterations and approximately 2.5 billion likelihood evaluations. This total includes 304 candidate models for Stage 1 climate interpolation, 140 beta-regression decision round fits (14 valid models across 4 growth forms, 10 posterior draws each), 8,000 beta-regression production fits (8 models across 1,000 posterior draws), 750 Dirichlet decision round fits (75 candidate models, 10 posterior draws each), and 11,800 Dirichlet production fits (12 models across up to 1,000 posterior draws). Computation time totaled approximately 54 to 61 hours across all stages. Only 2 divergent transitions were recorded across all 20,994 fits (both in Stage 1 climate interpolation), indicating that the sampler settings were appropriate for the model complexity at every stage. As a minor side note, if we had tested every single climate posterior available (∼600 total) using the full 1000 draw option, then we estimated it would have taken approximately half a year of continuous computation time to complete; opting instead for a smaller hypothesis-based set of predictors was the obvious choice.

For Stage 1 climate interpolation, the 76 selected models (one per unique variable-month combination) had a mean R-hat of 1.004 (range 1.001 to 1.013), with 96% of models meeting the strict R-hat < 1.01 criterion. Minimum bulk effective sample size across selected models ranged from 778 to 3,626 (mean 2,138). Non-linear models (natural splines) were selected for 84% of comparisons, confirming that climate-elevation relationships along this gradient are predominantly non-linear. No linear model was selected for any relative humidity variable. For Stage 5 ecological models, the 12 Dirichlet production models achieved 97.5% to 99.5% convergence rates across 1,000 posterior draws, with zero divergent transitions across all 11,800 fits. The 8 beta-regression production models achieved 100% convergence with zero divergences across 8,000 fits. Full per-model diagnostics, including convergence rates, R-hat distributions, effective sample sizes, and model selection reasoning, can be provided upon request.

#### Computational resources and memory management

All analyses were run on a custom-built PC desktop workstation with the following specifications: AMD Ryzen 9 7900X 12-core processor (4.7 GHz), 32 GB DDR5 RAM (6000 MT/s), 16 GB dedicated graphics card, and a 1 TB NVMe SSD, which was partitioned into separate drives for software (C:) and data storage (D:), with approximately 1 TB of available working space. The operating system was Windows 11.

Memory management was a significant practical concern given the volume of repeated model fits (e.g., 12 models × 1,000 draws for the Dirichlet production runs, each requiring compilation, sampling, and LOO-IC calculation). Typical memory usage was ∼60%, but it could often spike to 99% usage which required manually stopping analyses and restarting of the computer. We took several steps to maximize throughput and reduce failures. First, for all Bayesian ecological models we used the “cmdstanr” backend rather than “rstan” R package (which was tried as a part of preliminary runs and tended to take 4-5x longer to run), because we found that “cmdstanr” compiles Stan models to standalone executables with lower memory overhead per fit and more efficient garbage collection between runs. Second, we re-routed temporary files that were generated during model compilation and sampling to a dedicated directory on the C: drive, outside of the default Windows %TEMP% location. This avoided two recurring issues, the first being the Windows path character limits that occasionally can cause failures with deeply nested temporary directories, and the second being antimalware software that flagged the rapid creation and deletion of temporary files during sequential model fitting. This temp directory also provided a simple method to check that the model was running as intended, with purging of temporary files between individual model fits to prevent excess accumulation.

For the most computationally intensive production runs, we typically ran two R Studio sessions simultaneously rather than using a multi-chain or multi-core parallelization within a single session (which was also tried during preliminary runs with less success). We found that multi-core approaches introduced more complexity than they resolved (aka too many cooks in the metaphorical kitchen), with the act of coordinating parallel chains across cores creating a massive overhead that offset the theoretical speed gains and overall output tracking became more error prone. Two independent sessions, each focusing on a different subset of models, provided us with a more reliable workflow. Additional memory usage best practices included minimizing the use of other programs during extended runs (especially anything internet related); disabling automatic backups and scheduled system updates; and lastly, retaining only the pooled summary statistics in checkpoint files (i.e., saved in increments of 50 draws) rather than full model objects, except for the overall “best” models where the full output is used for figures or as additional quality diagnostics estimates.

#### Prior specifications

Prior distributions for all Bayesian models are summarized in the section below, organized by model stage. Stage 1 was for climate interpolation, with models that used data-informed intercept priors centered on the monthly (or seasonal) mean of observed station values. The prior standard deviation was scaled by variable type to reflect expected variance of each factor, such as narrower standard deviation for variables with natural bounds (e.g., relative humidity maximum, which approaches 100%) and wider for variables with greater spatial variability (e.g., relative humidity minimum, precipitation). Slope priors were based on weakly informative Student-t distributions allowing for more flexibility in the potential altitude-climate relationship space, with the residual standard deviation (sigma) using an exponential prior. For precipitation models, with a greater overall variability of seasonal precipitation totals (due to monsoon as compared to dry season extremes), we used a wider slope and sigma priors.

Stages 2-4 were devoted to temporal aggregation (e.g. monthly to seasonal), calculation of complex climate derivatives (e.g. vapor pressure deficit), and custom metrics (e.g. threshold counts for freezing). Stage 5 was the second major Bayesian modeling stage, divided into two analysis types, beta-regression (for individual growth forms) and Dirichlet-regression (for dynamics between growth forms), both using climate predictors that were generated as a part of Stage 1. The intercept priors for Stage 5 ecological models were centered on the logit-transformed (beta-regression) or log-ratio-transformed (Dirichlet) mean proportion of each growth form across the respective ranges of each analysis. Slope priors were centered at zero, with standard deviations of 1.5, which is wide enough to allow substantial effects on the standardized predictor scale while providing regularization against extreme values. For the Dirichlet intercept prior for the conifer group, we used a slightly wider standard deviation (3.0) to allow adequate posterior exploration given the sparseness of conifer occurrence followed by rapid proportional increase toward the ridge summit. The beta-regression precision parameter (phi) used a gamma prior (2, 0.1), placing most mass on moderate to high precision values consistent with the observed concentration of RBA proportions. The Dirichlet model used the brms default prior for the precision parameter.

### Appendix S9: Table S2

#### Prior specifications for all Bayesian models

##### Stage 1: Climate interpolation (temperature and relative humidity)

| Parameter | Distribution | Notes |
| --- | --- | --- |
| Intercept (temp_min) | Normal (monthly mean, 12) | Wider SD: temperature inversions increase low-elevation variability |
| Intercept (temp_avg) | Normal (monthly mean, 10) | Moderate SD |
| Intercept (temp_max) | Normal (monthly mean, 12) | Wider SD: solar radiation effects |
| Intercept (rh_min) | Normal (monthly mean, 20) | Widest SD: dry extremes are highly variable across the gradient |
| Intercept (rh_avg) | Normal (monthly mean, 15) | Moderate SD |
| Intercept (rh_max) | Normal (monthly mean, 10) | Tightest SD: values approach 100% ceiling |
| Slopes (all variables) | Student's t (3, 0, 10) | Weakly informative; heavy tails allow non-linear altitude effects |
| Sigma (all variables) | Exponential (1) | Weakly informative; constrains residual SD to positive values |

*Monthly intercept prior means for each variable are provided upon request, or can be calculated using* *the file "raw\_climate\_monthly.csv". These observed climate values ranged from 4.6 to 17.6 °C for* *temp\_min, 8.1 to 19.0 °C for temp\_avg, 11.4 to 20.5 °C for temp\_max, 43.4 to 89.3% for rh\_min, 74.0 to* *95.0% for rh\_avg, and 89.4 to 98.3% for rh\_max.*

##### Stage 1: Climate interpolation (precipitation)

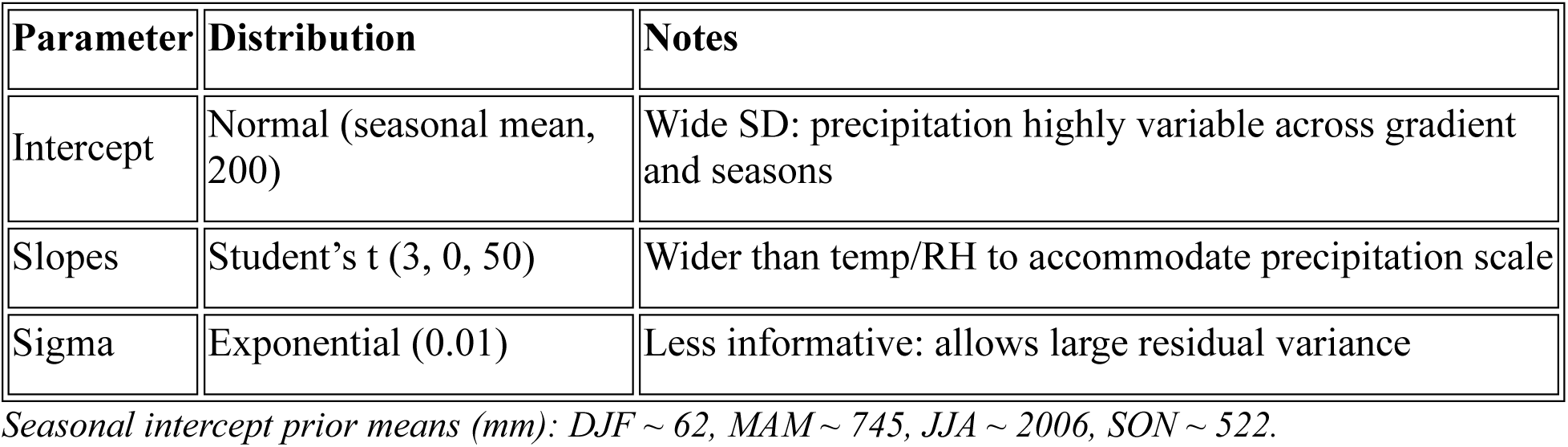

##### Stage 5: Beta-regression (individual growth form models)

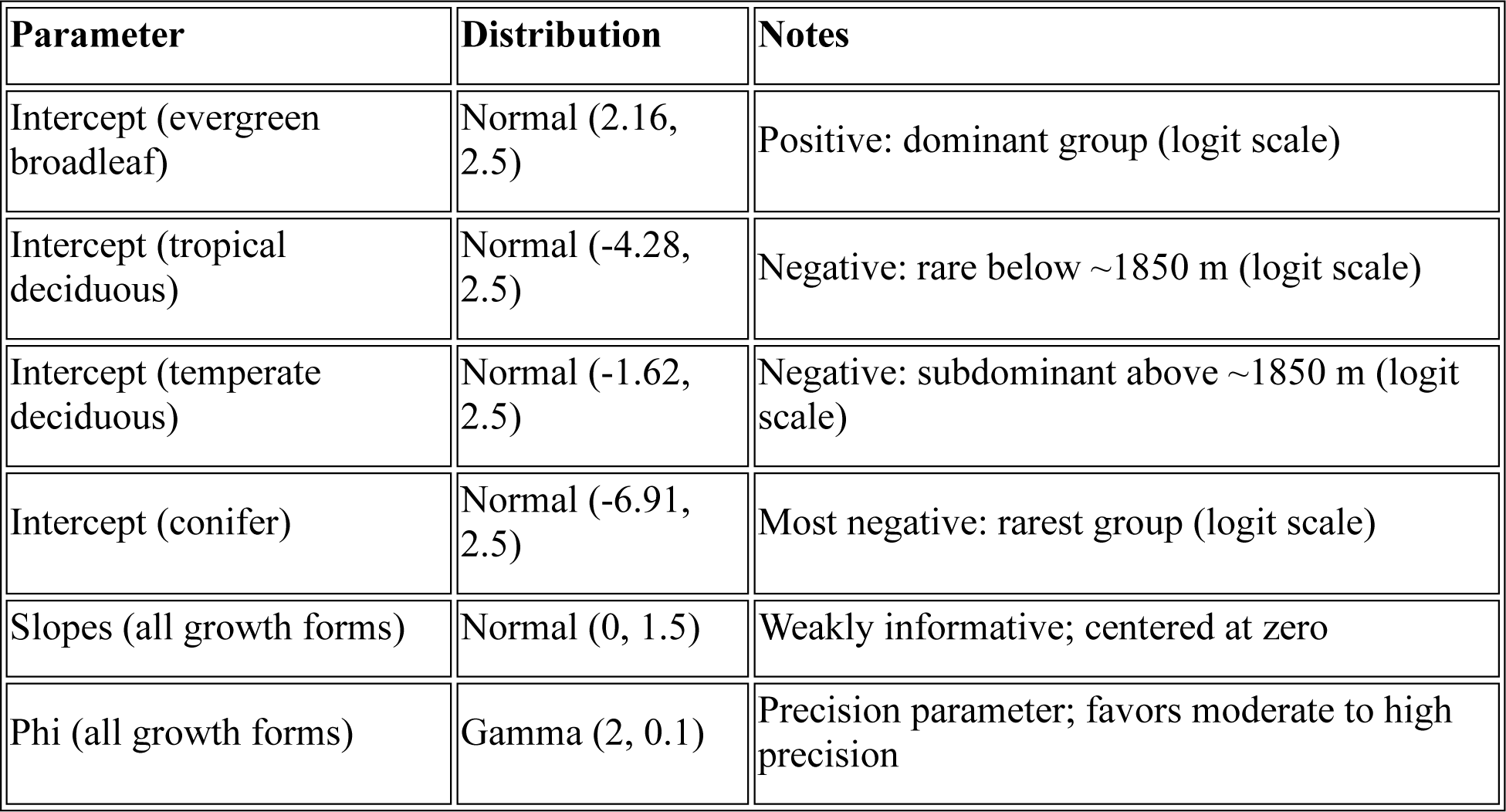

| Parameter | Distribution | Notes |
| --- | --- | --- |
| Intercept (evergreen broadleaf) | Normal (2.16, 2.5) | Positive: dominant group (logit scale) |
| Intercept (tropical deciduous) | Normal (-4.28, 2.5) | Negative: rare below ~1850 m (logit scale) |
| Intercept (temperate deciduous) | Normal (-1.62, 2.5) | Negative: subdominant above ~1850 m (logit scale) |
| Intercept (conifer) | Normal (-6.91, 2.5) | Most negative: rarest group (logit scale) |
| Slopes (all growth forms) | Normal (0, 1.5) | Weakly informative; centered at zero |
| Phi (all growth forms) | Gamma (2, 0.1) | Precision parameter; favors moderate to high precision |

##### Stage 5: Dirichlet compositional model (4-group RBA)

| Parameter | Distribution | Notes |
| --- | --- | --- |
| Intercept (conifer vs evergreen) | Normal (-2.5, 3.0) | Wider SD for sparse group |
| Intercept (tropical deciduous vs evergreen) | Normal (-3.2, 2.5) | Log-ratio scale relative to reference |
| Intercept (temperate deciduous vs evergreen) | Normal (-2.2, 2.5) | Log-ratio scale relative to reference |
| Slopes (conifer) | Normal (0, 1.5) | Consistent across all modeled categories |
| Slopes (tropical deciduous) | Normal (0, 1.5) | Widened from initial value of 1.0 for sparse group |
| Slopes (temperate deciduous) | Normal (0, 1.5) | Widened from initial value of 1.0 for sparse group |
| Phi (precision) | Gamma (0.01, 0.01), brms default | Gamma prior on Dirichlet precision parameter |

*Note: Evergreen broadleaf serves as the reference category in the Dirichlet model; its intercept and* *slopes are implicitly zero. All intercept priors are on the log-ratio scale relative to the reference category.* *Negative values indicate lower expected proportion relative to evergreen broadleaf.*

### Appendix S9: Section S3

#### Statistical literature, software programs, or packages used

##### Software and Programming Environment

1. **R.** Programming environment for all statistical analyses, data processing, and figure generation. *Citation:* R Core Team, 2026. R: A language and environment for statistical computing. R Foundation for Statistical Computing, Vienna, Austria. https://www.R-project.org/

##### Bayesian Modeling

2. **R package "brms".** Used for all Bayesian regression models across Stages 1–5: climate interpolation (linear and natural spline candidates), beta regressions (null comparison tests), and Dirichlet compositional models (4-group RBA). *Citation:* Bürkner, P.-C., 2017. brms: An R package for Bayesian multilevel models using Stan. Journal of Statistical Software, 80(1), 1–28. https://doi.org/10.18637/jss.v080.i01
3. **R package "brms"**. Extended capabilities for non-standard response families (Dirichlet, beta) and complex formula syntax used in Stage 5. *Citation:* Bürkner, P.-C., 2018. Advanced Bayesian multilevel modeling with the R package brms. The R Journal, 10(1), 395–411. https://doi.org/10.32614/RJ-2018-017
4. **R package "parallel".** Base R package used for parallel execution of MCMC chains across multiple CPU cores, reducing computation time for the 8-chain Stage 1 models and repeated Stage 5 model fits. *Citation:* Included in R base (R Core Team, 2024).
5. **R package "cmdstanr" version 0.8**. Lightweight R interface to CmdStan, the command-line interface to Stan’s C++ implementation. Used as the backend for all Bayesian ecological models, providing faster compilation and lower memory overhead than rstan for the high volume of repeated Dirichlet model fits (12 models × 1,000 draws). *Citation:* Gabry, J., Češnovar, R., Johnson, A., Bronder, S., 2026. cmdstanr: R Interface to CmdStan. R package. https://mc-stan.org/cmdstanr/
6. **Stan software (written in C++).** Probabilistic programming language providing the Hamiltonian Monte Carlo (HMC) sampling engine underlying all brms models. For preliminary we used the rstan backend, but shifted to cmdstanr for improved computational efficiency with repeated model fits. *Citation:* Stan Development Team, 2024. Stan modeling language users guide and reference manual, version 2.38. https://mc-stan.org/

##### Model Evaluation and Diagnostics

7. **R package "loo".** Used for model selection via LOO-IC (leave-one-out information criterion) at all stages, including the conditional complexity algorithm and pooled model comparison under Rubin’s rules. Implements Pareto-smoothed importance sampling (PSIS-LOO). *Citation:* Vehtari, A., Gelman, A., Gabry, J., 2017. Practical Bayesian model evaluation using leave-one-out cross-validation and WAIC. Statistics and Computing, 27(5), 1413–1432. https://doi.org/10.1007/s11222-016-9696-4 *Software citation:* Vehtari, A., Gabry, J., Magnusson, M., Yao, Y., Bürkner, P.-C., Paananen, T., Gelman, A., 2024. loo: Efficient leave-one-out cross-validation and WAIC for Bayesian models. R package. https://mc-stan.org/loo/
8. **R package "posterior".** Used for convergence diagnostics: ESS calculation filtered to model parameters matching ^b_|sigma|phi, excluding auxiliary parameters with artificially low ESS. *Citation:* Posterior Developers, 2024. posterior: Tools for working with posterior distributions. R package. https://mc-stan.org/posterior/
9. **Gelman-Rubin diagnostic (R-hat).** Used to assess MCMC convergence with threshold R-hat < 1.01 for all model parameters. *Citation:* Gelman, A., Rubin, D.B., 1992. Inference from iterative simulation using multiple sequences. Statistical Science, 7(4), 457–472. https://doi.org/10.1214/ss/1177011136

##### Uncertainty Propagation

10. **Rubin’s rules.** Used to pool parameter estimates, standard errors, and LOO-IC values across 1,000 climate posterior draws, treating each draw as one plausible reality; analogous to multiple imputation for covariate measurement error. *Citation:* Rubin, D.B., 1987. Multiple Imputation for Nonresponse in Surveys. John Wiley & Sons, New York.

##### Community Analysis

11. **R package "vegan".** Used for Mantel tests (matrix correlation between community and climate distances), partial Mantel tests (controlling for elevation), distance-based redundancy analysis (dbRDA on Bray-Curtis dissimilarity of 4-group RBA), and variance partitioning (climate-unique vs elevation-unique explained variation). *Citation:* Oksanen, J., Simpson, G.L., Blanchet, F.G., Kindt, R., Legendre, P., Minchin, P.R., O’Hara, R.B., Solymos, P., Stevens, M.H.H., Szoecs, E., Wagner, H., et al., 2024. vegan: Community Ecology Package. R package. https://CRAN.R-project.org/package=vegan
12. **dbRDA methodology.** *Citation:* Legendre, P., Anderson, M.J., 1999. Distance-based redundancy analysis: testing multispecies responses in multifactorial ecological experiments. Ecological Monographs, 69(1), 1–24. https://doi.org/10.1890/0012-9615(1999)069[0001:DBRATM]2.0.CO;2

##### Climate Derivation Methods

13. **Tetens equation.** Used for saturation vapor pressure calculation in VPD and dewpoint derivations (Stage 2). *Citation:* Tetens, O., 1930. Über einige meteorologische Begriffe. Zeitschrift für Geophysik, 6, 297–309. *See also:* Murray, F.W., 1967. On the computation of saturation vapor pressure. Journal of Applied Meteorology, 6(1), 203–204.
14. **Inverse Tetens equation (dewpoint)**. Used for dewpoint temperature and fog gap derivations (Stage 2). *Citation:* Lawrence, M.G., 2005. The relationship between relative humidity and the dewpoint temperature in moist air: A simple conversion and applications. Bulletin of the American Meteorological Society, 86(2), 225–234. https://doi.org/10.1175/BAMS-86-2-225
15. **Hargreaves-Samani method.** Used for potential evapotranspiration (PET) estimation from temperature data (Stage 3). *Citation:* Hargreaves, G.H., Samani, Z.A., 1985. Reference crop evapotranspiration from temperature. Applied Engineering in Agriculture, 1(2), 96–99. https://doi.org/10.13031/2013.26773
16. **FAO 56 reference framework.** Standard reference for evapotranspiration methodology; provides context for the Hargreaves-Samani simplified approach used here. *Citation:* Allen, R.G., Pereira, L.S., Raes, D., Smith, M., 1998. Crop Evapotranspiration: Guidelines for Computing Crop Water Requirements. FAO Irrigation and Drainage Paper No. 56. Food and Agriculture Organization of the United Nations, Rome.
17. **Holdridge life zone system.** Used for biotemperature (ABT) calculation and classic potential evapotranspiration ratio (PER_class) in Stages 2–3. *Citation:* Holdridge, L.R., 1967. Life Zone Ecology. Tropical Science Center, San José, Costa Rica.
18. **Kira warmth index.** Used for thermal vegetation zonation; cumulative thermal energy above 5°C base temperature (Stage 2). *Citation:* Kira, T., 1945. A New Classification of Climate in Eastern Asia as the Basis for Agricultural Geography. Horticultural Institute, Kyoto University, Kyoto, 1–23.

##### AI Assistance

19. **Claude (Anthropic).** Used as needed, under direct supervision of the authors (mostly for scaling of analyses and troubleshooting). All analytical decisions, ecological interpretations, and scientific conclusions are the responsibility of the authors. *Citation:* Anthropic, 2026. Claude Opus 4.6 [Large language model]. https://www.anthropic.com

### Appendix S10: Section S1

#### Descriptions of additional files and data

Files 1 (.md) and 2 (.pdf) are included as Supporting information. All code and datasets (1-4) for analyses are publicly available at Zenodo at https://doi.org/10.5281/zenodo.19081441. For questions about field data or sampling protocol, contact Pema Wangda.

For questions about statistics, R code, contact Melissa Whitman.

#### File 1. Reproducible analysis pipeline (aka all code used for this study)

**File name:** Full code archive (see README.md)

**Description:** The Zenodo archive contains all R code and intermediate data needed to replicate the Bayesian modeling pipeline described in this paper. The pipeline spans 24 scripts organized in 6 stages: Stage 1 (climate interpolation from 8 stations to 28 sites), Stage 2 (monthly derivatives including VPD, fog gap, and warmth index), Stage 3 (temporal aggregation and precipitation-derived metrics), Stage 4 (ecological threshold metrics), Stage 5 (community analysis, beta regression with null model comparisons, and Dirichlet compositional regression), and Stage 6 (all manuscript and Supporting Information figures). The archive includes a detailed README documenting every script, its inputs and outputs, runtime estimates, prior specifications, and methodological notes. All scripts use base R with brms and cmdstanr for Bayesian models. A test mode (10 posterior draws) runs in approximately 4–6 hours; the full 1,000-draw production version requires approximately 54–61 hours total, inclusive of all stages and accounting for practical considerations such as installation of the cmdstanr backend and C++ toolchain (which requires additional setup beyond standard R package installation), configuring a dedicated temporary file directory, drive partitioning for data storage, troubleshooting, and hardware variability. All scripts use a fixed random seed (12345) for exact reproducibility.

#### File 2. Monthly plots of interpolated climate data and derivatives

**File name:** monthly_diagnostics_Bayesian.pdf

**Description:** This file provides example plots of climate model output for quality control evaluation purposes, performed using monthly data prior to seasonal aggregation or inclusion in forest compositional models. Additional plots used in the methodological evaluation stage, including seasonal plots or calculation of custom metrics, available upon request. Known site values are shown as points outlined in black, interpolated values for plots along the gradient shown as points with neutral outline, and credible intervals as ribbons. Transparent boxes are used to illustrate when conditions cross a key threshold of interest, such as temperatures below freezing. Red lines and ribbons indicate the estimated maximum values over the given month, and blue the minimum, with black as monthly average; orange values are “noon” values, and purple are “dawn”, based on thermodynamic estimates.

#### Dataset 1. Monthly climate data

**File name:** raw_climate_monthly.csv

**Description:** File name: raw_climate_monthly.csv Description: Dataset of temperature (°C) and relative humidity (%) data, collected at eight sites along an altitudinal gradient, as well as rainfall precipitation (mm) for three sites, with additional precipitation data from weather stations (Dorji, 2016). Data were collected daily but summarized as monthly averages and absolute minimums and maximums. This is the raw data used as the foundation of the climate interpolation models.

#### Dataset 2. Relative Basal Area by growth form

**File name:** growthform_rba.csv

**Description:** Relative basal area and distributions of four growth forms (tropical deciduous, evergreen broadleaf, temperate deciduous, and conifer). Cell values represent the proportion of total basal area (cm²) sampled per plot, with altitude as rows and growth form groups as columns. Data are based on a moist slope survey of woody trees and shrubs more than 1.3 m tall. This file also contains site identifiers and altitude values used throughout the modeling pipeline.

#### Dataset 3. Seasonal precipitation data

**File name:** Dorji_ppt_seasonal.csv

**Description:** Seasonal precipitation totals (mm) for 15 stations along the gradient, combining three field-collected sites with 12 additional weather station records (Dorji et al. 2016). Seasons are DJF, MAM, JJA, and SON. Used for precipitation interpolation in Stage 3 of the pipeline.

#### Dataset 4. Species-level RBA matrix

**File name:** moist_slope_rba_anonymous.csv

**Description:** Relative basal area (%) by growth form for 145 species across 28 sites, anonymized (e.g. species names replaced with spp01, spp02, etc…). Columns 3:31 represent the 28 plot, the sum of each column equals 1; the first half of each column name indicates plot identifier (e.g. WS.P07) and the last four digits representing altitude of samples, separated by a period. Used for vegetation zone dendrogram, dbRDA community analysis, and variance partitioning. For complete (non-anonymous) species RBA and plot-level information, contact Pema Wangda at.

#### Dataset 5. Species inventory list

**File name:** species_inventory.csv

**Description:** Species sampled along the moist slope transect, from 450 to 3370 m a.s.l., including details on Family, Genus, growth form, and size group. Two columns represent species names, the first column “original_moist” represents the name recorded in this study, often corresponding with historical nomenclature used in Flora of Bhutan; the second column “name_phylo” represents the more contemporary name or synonym used by the World Flora Online (https://wfoplantlist.org/) or for the creation of phylogenetic mega-trees.

## References

Aguirre-Gutiérrez, J., E. Berenguer, I. Oliveras Menor, D. Bauman, J. J. Corral-Rivas, M. G. Nava-Miranda, S. Both, J. E. Ndong, F. E. Ondo, N. N. Bengone, V. Mihinhou, J. W. Dalling, K. Heineman, A. Figueiredo, R. González-M, N. Norden, A. B. Hurtado-M, D. González, B. Salgado-Negret, S. M. Reis, M. M. Moraes de Seixas, W. Farfan-Rios, A. Shenkin, T. Riutta, C. A. J. Girardin, S. Moore, K. Abernethy, G. P. Asner, L. P. Bentley, D. F. R. P. Burslem, L. A. Cernusak, B. J. Enquist, R. M. Ewers, J. Ferreira, K. J. Jeffery, C. A. Joly, B. H. Marimon-Junior, R. E. Martin, P. S. Morandi, O. L. Phillips, A. C. Bennett, S. L. Lewis, C. A. Quesada, B. S. Marimon, W. D. Kissling, M. Silman, Y. A. Teh, L. J. T. White, N. Salinas, D. A. Coomes, J. Barlow, S. Adu-Bredu, and Y. Malhi. 2022. Functional susceptibility of tropical forests to climate change. Nature Ecology & Evolution 6:878–889.

Ashton, P. S. 2014. On the Forests of Tropical Asia Lest the Memory Fade. Royal Botanic Gardens, Kew, in association with the Arnold Arboretum of Harvard University., Kew, UK.

Ashton, P. S., S. I. Aiba, H. Zhu, and R. Pradhan. 2022. Further food for thought: higher tropical mountains revisited once again. Tropics 31:43–57.

Ashton, P. S., and H. Zhu. 2020. The tropical-subtropical evergreen forest transition in East Asia: An exploration. Plant Diversity 42:255–280.

Bader, M. Y., L. D. Llambí, B. S. Case, H. L. Buckley, J. M. Toivonen, J. J. Camarero, D. M. Cairns, C. D. Brown, T. Wiegand, and L. M. Resler. 2021. A global framework for linking alpine-treeline ecotone patterns to underlying processes. Ecography 44:265–292.

Bartlett, J. W., and R. H. Keogh. 2018. Bayesian correction for covariate measurement error: A frequentist evaluation and comparison with regression calibration. Statistical Methods in Medical Research 27:1695–1708.

Bhattarai, K. R., and O. R. Vetaas. 2006. Can Rapoport’s rule explain tree species richness along the Himalayan elevation gradient, Nepal? Diversity and Distributions 12:373–378.

Blomberg, S. P., and O. S. Todorov. 2025. The fallacy of single imputation for trait databases: Use multiple imputation instead. Methods in Ecology and Evolution 16:658–667.

Bracewell, S. A., E. L. Johnston, and G. F. Clark. 2024. Variation in Successional Dynamics Shape Biodiversity Patterns over a Tropical-Temperate Latitudinal Gradient. The American Naturalist 204:327–344.

Bray, J. R., and J. T. Curtis. 1957. An Ordination of the Upland Forest Communities of Southern Wisconsin. Ecological Monographs 27:325–349.

Brown, C. J., and S. Spillias. 2026. Prompting large language models for quality ecological statistics. Methods in Ecology and Evolution 17:1012–1021.

Brown, J. H., G. C. Stevens, and D. M. Kaufman. 1996. The geographic range: Size, shape, boundaries, and internal structure. Annual Review of Ecology and Systematics 27:597–623.

Bruijnzeel, L. A., and E. J. Veneklaas. 1998. Climatic Conditions and Tropical Montane Forest Productivity: The Fog Has Not Lifted Yet. Ecology 79:3–9.

Camarero, J. J., A. Gazol, R. Sánchez-Salguero, A. Fajardo, E. J. B. McIntire, E. Gutiérrez, E. Batllori, S. Boudreau, M. Carrer, J. Diez, G. Dufour-Tremblay, N. P. Gaire, A. Hofgaard, V. Jomelli, A. V. Kirdyanov, E. Lévesque, E. Liang, J. C. Linares, I. E. Mathisen, P. A. Moiseev, G. Sangüesa-Barreda, K. B. Shrestha, J. M. Toivonen, O. V. Tutubalina, and M. Wilmking. 2021. Global fading of the temperature–growth coupling at alpine and polar treelines. Global Change Biology 27:1879–1889.

Carpenter, B., A. Gelman, M. D. Hoffman, D. Lee, B. Goodrich, M. Betancourt, M. Brubaker, J. Guo, P. Li, and A. Riddell. 2017. Stan: A Probabilistic Programming Language. Journal of Statistical Software 76:1–32.

Champion, H. G. 1936. A preliminary survey of the forest types of India and Burma. Indian Forest Records 1:1–286.

Clark, J. S. 2005. Why environmental scientists are becoming Bayesians. Ecology Letters 8:2–14.

Colwell, R. K., and D. C. Lees. 2000. The mid-domain effect: Geometric constraints on the geography of species richness. Trends in Ecology and Evolution 15:70–76.

Damgaard, C. F., and K. M. Irvine. 2019. Using the beta distribution to analyse plant cover data. Journal of Ecology 107:2747–2759.

Davies, S. J., I. Abiem, K. Abu Salim, S. Aguilar, D. Allen, A. Alonso, K. Anderson-Teixeira, A. Andrade, G. Arellano, P. S. Ashton, P. J. Baker, M. E. Baker, J. L. Baltzer, Y. Basset, P. Bissiengou, S. Bohlman, N. A. Bourg, W. Y. Brockelman, S. Bunyavejchewin, D. F. R. P. Burslem, M. Cao, D. Cárdenas, L.-W. Chang, C.-H. Chang-Yang, K.-J. Chao, W.-C. Chao, H. Chapman, Y.-Y. Chen, R. A. Chisholm, C. Chu, G. Chuyong, K. Clay, L. S. Comita, R. Condit, S. Cordell, H. S. Dattaraja, A. A. de Oliveira, J. den Ouden, M. Detto, C. Dick, X. Du, Á. Duque, S. Ediriweera, E. C. Ellis, N. L. E. Obiang, S. Esufali, C. E. N. Ewango, E. S. Fernando, J. Filip, G. A. Fischer, R. Foster, T. Giambelluca, C. Giardina, G. S. Gilbert, E. Gonzalez-Akre, I. A. U. N. Gunatilleke, C. V. S. Gunatilleke, Z. Hao, B. C. H. Hau, F. He, H. Ni, R. W. Howe, S. P. Hubbell, A. Huth, F. Inman-Narahari, A. Itoh, D. Janík, P. A. Jansen, M. Jiang, D. J. Johnson, F. A. Jones, M. Kanzaki, D. Kenfack, S. Kiratiprayoon, K. Král, L. Krizel, S. Lao, A. J. Larson, Y. Li, X. Li, C. M. Litton, Y. Liu, S. Liu, S. K. Y. Lum, M. S. Luskin, J. A. Lutz, H. T. Luu, K. Ma, J.-R. Makana, Y. Malhi, A. Martin, C. McCarthy, S. M. McMahon, W. J. McShea, H. Memiaghe, X. Mi, D. Mitre, M. Mohamad, L. Monks, H. C. Muller-Landau, P. M. Musili, J. A. Myers, A. Nathalang, K. M. Ngo, N. Norden, V. Novotny, M. J. O’Brien, D. Orwig, R. Ostertag, K. Papathanassiou, G. G. Parker, R. Pérez, I. Perfecto, R. P. Phillips, N. Pongpattananurak, H. Pretzsch, H. Ren, G. Reynolds, L. J. Rodriguez, S. E. Russo, L. Sack, W. Sang, J. Shue, A. Singh, G.-Z. M. Song, R. Sukumar, I.-F. Sun, H. S. Suresh, N. G. Swenson, S. Tan, S. C. Thomas, D. Thomas, J. Thompson, B. L. Turner, A. Uowolo, M. Uriarte, R. Valencia, J. Vandermeer, A. Vicentini, M. Visser, T. Vrska, X. Wang, X. Wang, G. D. Weiblen, T. J. S. Whitfeld, A. Wolf, S. J. Wright, H. Xu, T. L. Yao, S. L. Yap, W. Ye, M. Yu, M. Zhang, D. Zhu, L. Zhu, J. K. Zimmerman, and D. Zuleta. 2021. ForestGEO: Understanding forest diversity and dynamics through a global observatory network. Biological Conservation 253:108907.

Delcourt, P. A., and H. R. Delcourt. 1992. Ecotone Dynamics in Space and Time. Pages 19–54 in A. J. Hansen and F. di Castri, editors. Landscape Boundaries: Consequences for Biotic Diversity and Ecological Flows. Springer, New York, NY.

Dorji, U., J. E. Olesen, P. K. Bøcher, and M. S. Seidenkrantz. 2016. Spatial Variation of Temperature and Precipitation in Bhutan and Links to Vegetation and Land Cover. Mountain Research and Development 36:66–79.

Doughty, C. E., J. M. Keany, B. C. Wiebe, C. Rey-Sanchez, K. R. Carter, K. B. Middleby, A. W. Cheesman, M. L. Goulden, H. R. da Rocha, S. D. Miller, Y. Malhi, S. Fauset, E. Gloor, M. Slot, I. Oliveras Menor, K. Y. Crous, G. R. Goldsmith, and J. B. Fisher. 2023. Tropical forests are approaching critical temperature thresholds. Nature 2023 621:7977 621:105–111.

Douma, J. C., and J. T. Weedon. 2019. Analysing continuous proportions in ecology and evolution: A practical introduction to beta and Dirichlet regression. Methods in Ecology and Evolution 10:1412–1430.

Dumelle, M., R. Trangucci, A. M. Nahlik, A. R. Olsen, K. M. Irvine, K. Blocksom, J. M. Ver Hoef, and C. Fuentes. 2025. Missing data in ecology: Syntheses, clarifications, and considerations. Ecological Monographs 95:e70037.

Edwards, P. J., and P. J. Grubb. 1977. Studies of mineral cycling in a montane rain forest in New Guinea: I. The distribution of organic matter in the vegetation and soil. The Journal of Ecology 65:943.

Ellison, A. M. 2004. Bayesian inference in ecology. Ecology Letters 7:509–520.

Feeley, K., J. Martinez-Villa, T. Perez, A. Silva Duque, D. Triviño Gonzalez, and A. Duque. 2020. The thermal tolerances, distributions, and performances of tropical montane tree species. Frontiers in Forests and Global Change 3:25.

Fisher, J. B., Y. Malhi, I. C. Torres, D. B. Metcalfe, M. J. van de Weg, P. Meir, J. E. Silva-Espejo, and W. H. Huasco. 2013. Nutrient limitation in rainforests and cloud forests along a 3,000-m elevation gradient in the Peruvian Andes. Oecologia 172:889–902.

Gabry, J., R. Češnovar, A. Johnson, and S. Bronder. 2026. Authors and Citation.

Gallou, A., A. S. Jump, J. S. Lynn, R. Field, S. D. H. Irl, M. J. Steinbauer, C. Beierkuhnlein, J.-C. Chen, C.-H. Chou, A. Hemp, Y. Kidane, C. König, H. Kreft, A. Naqinezhad, A. Nowak, J.-N. Nuppenau, P. Trigas, J. P. Price, C. A. Roland, A. H. Schweiger, P. Weigelt, S. G. A. Flantua, and J.-A. Grytnes. 2023. Diurnal temperature range as a key predictor of plants’ elevation ranges globally. Nature Communications 2023 14:1 14:1–8.

Givnish, T. J. 1999. On the causes of gradients in tropical tree diversity. Journal of Ecology 87:193–210.

Givnish, T. J. 2002. Adaptive significance of evergreen vs. deciduous leaves: solving the triple paradox. Silva Fennica 36:703–743.

Givnish, T. J. Submitted. Elevational gradients in plant form and function: study design and conceptual framework for predicting and testing trait shifts on Mount Washington, New Hampshire. Annals of Botany.

Goldsmith, G. R., N. J. Matzke, and T. E. Dawson. 2013. The incidence and implications of clouds for cloud forest plant water relations. Ecology Letters 16:307–314.

González-Caro, S., Á. Duque, K. J. Feeley, E. Cabrera, J. Phillips, S. Ramirez, and A. Yepes. 2020. The legacy of biogeographic history on the composition and structure of Andean forests. Ecology 101:e03131.

Grime, J. P. 1977. Evidence for the existence of three primary strategies in plants and its relevance to ecological and evolutionary theory. American Naturalist 111:1169–1194.

Grossiord, C., T. N. Buckley, L. A. Cernusak, K. A. Novick, B. Poulter, R. T. W. Siegwolf, J. S. Sperry, and N. G. McDowell. 2020. Plant responses to rising vapor pressure deficit. New Phytologist 226:1550–1566.

Grubb, P. J. 1971. Interpretation of the “Massenerhebung” effect on tropical mountains. Nature 229:44–45.

Grubb, P. J., P. J. Bellingham, T. S. Kohyama, F. I. Piper, and A. Valido. 2013. Disturbance regimes, gap-demanding trees and seed mass related to tree height in warm temperate rain forests worldwide. Biological Reviews 88:701–744.

Grytnes, J.-A., J. H. Beaman, T. S. Romdal, and C. Rahbek. 2008. The mid-domain effect matters: simulation analyses of range-size distribution data from Mount Kinabalu, Borneo. Journal of Biogeography 35:2138–2147.

Gyaltshen, D., P. Wangda, and B. Suberi. 2014. Structure and Composition of the Natural Sal (Shorea robusta Gaertner f.) Forest, Gomtu, Southern Bhutan. Bhutan Journal of Natural Resources and Development 1:1–10.

Holdridge, L. R. 1967. Life zone ecology. Tropical Science Center, San Jose, Costa Rica.

Janzen, D. H. 1967. Why mountain passes are higher in the tropics. American Naturalist 101:233–249.

Joyce, E. M., K. R. Thiele, J. W. F. Slik, and D. M. Crayn. 2021. Plants will cross the lines: climate and available land mass are the major determinants of phytogeographical patterns in the Sunda–Sahul Convergence Zone. Biological Journal of the Linnean Society 132:374–387.

Juvik, J. O., and D. Nullet. 1995. Relationships between rainfall, cloud-water interception, and canopy throughfall in a Hawaiian montane forest. Pages 165–182 Tropical montane cloud forests. Ecological studies (analysis and synthesis), vol 110. Springer, New York, NY.

Kaurila, K., S. Kuningas, A. Lappalainen, and J. Vanhatalo. 2026. Species distribution modeling with expert elicitation and Bayesian calibration. Ecography 2026:e08173.

Khandu, Y., A. Polthanee, and S. Isarangkool Na Ayutthaya. 2022. Ecological dynamics and regeneration expansion of treeline ecotones in response to climate change in Northern Bhutan Himalayas. Forests 13:1062.

Kidane, Y. O., M. J. Steinbauer, and C. Beierkuhnlein. 2019. Dead end for endemic plant species? A biodiversity hotspot under pressure. Global Ecology and Conservation 19:e00670.

Kira, T. 1945. A new classification of climate in eastern Asia as the basis for agricultural geography. Horticultural Institute Kyoto University, Kyoto, Japan.

Kira, T. 1991. Forest ecosystems of east and southeast Asia in a global perspective. Ecological Research 6:185–200.

Kitayama, K. 1992. An altitudinal transect study of the vegetation on Mount Kinabalu, Borneo. Vegetatio 102:149–171.

Kluge, J., S. Worm, S. Lange, D. Long, J. Böhner, R. Yangzom, and G. Miehe. 2017. Elevational seed plants richness patterns in Bhutan, Eastern Himalaya. Journal of Biogeography 44:1711–1722.

Korner, C., A. Allison, and H. Hilscher. 1983. Altitudinal variation of leaf diffusive conductance and leaf anatomy in heliophytes of montane New Guinea and their interrelation with microclimate. Flora, Jena 174:91–135.

Legendre, P., and L. Legendre. 2012. Numerical Ecology. Elsevier.

Leng, R., S. Harrison, D. Fawcett, A. Tiwari, M. Harrison, and K. Anderson. 2026. Vegetation on the move: elevational shifts and greening dynamics across the Himalayan alpine zone. Ecography e08259:e08259.

Li, C. F., M. Chytrý, D. Zelený, M. Y. Chen, T. Y. Chen, C. R. Chiou, Y. J. Hsia, H. Y. Liu, S. Z. Yang, C. L. Yeh, J. C. Wang, C. F. Yu, Y. J. Lai, W. C. Chao, and C. F. Hsieh. 2013. Classification of Taiwan forest vegetation. Applied Vegetation Science 16:698–719.

Lynn, J. S., M. R. Kazenel, S. N. Kivlin, and J. A. Rudgers. 2019. Context-dependent biotic interactions control plant abundance across altitudinal environmental gradients. Ecography 42:1600–1612.

Mangen, J.-M. 1993. Ecology and Vegetation of Mt Trikora New Guinea (lrian Jaya I Indonesia). Musée national d’histoire naturelle de Luxembourg, Luxembourg.

Manly, B. F. J., and B. F. J. Manly. 2018. Randomization, Bootstrap and Monte Carlo Methods in Biology. Third edition. Chapman and Hall/CRC, New York.

Mayor, J. R., N. J. Sanders, A. T. Classen, R. D. Bardgett, J.-C. Clément, A. Fajardo, S. Lavorel, M. K. Sundqvist, M. Bahn, C. Chisholm, E. Cieraad, Z. Gedalof, K. Grigulis, G. Kudo, D. L. Oberski, and D. A. Wardle. 2017. Elevation alters ecosystem properties across temperate treelines globally. Nature 542:91–95.

Morley, R. J. 2000. Origin and Evolution of Tropical Rain Forests. Wiley, Chichester.

Murtagh, F., and P. Legendre. 2014. Ward’s Hierarchical Agglomerative Clustering Method: Which Algorithms Implement Ward’s Criterion? Journal of Classification 31:274–295.

Norbu, L. 2000. Cattle grazing - an integral part of broadleaf forest management planning in Bhutan. ETH Zürich.

Ohsawa, M. 1984. Differentiation of vegetation zones and species strategies in the subalpine region of Mt. Fuji. Vegetatio 57:15–52.

Ohsawa, M. 1987. Vegetation zones in the Bhutan Himalaya. Pages 1–71 in M. Ohsawa, editor. Life Zone Ecology of Bhutan Himalaya. Chiba University, Japan.

Ohsawa, M. 1990. An interpretation of latitudinal patterns of forest limits in South and East Asian mountains. Journal of Ecology 78:326–339.

Ohsawa, M. 1991. Life Zone Ecology of the Bhutan Himalaya II. Chiba University, Japan.

Ohsawa, M. 1992. Altitudinal zonation and succession of forests in the eastern Himalaya. Braun-Blanquetia 8:92–98.

Ohsawa, M. 1993. Latitudinal pattern of mountain vegetation zonation in southern and eastern Asia. Journal of Vegetation Science 4:13–18.

Proctor, J., K. Haridasan, and Smith. 1998. How far north does Lowland Evergreen Tropical Rain Forest go? Global Ecology & Biogeography Letters 7:141–146.

R Core Team. 2026. R: A language and environment for statistical computing. R Foundation for Statistical Computing, Vienna, Austria.

Rahbek, C. 1995. The elevational gradient of species richness: a uniform pattern? Ecography 18:200–205.

Rehm, E. M., and K. J. Feeley. 2015. The inability of tropical cloud forest species to invade grasslands above treeline during climate change: potential explanations and consequences. Ecography 38:1167–1175.

Richardson, J. E., C. M. Costion, and A. N. Muellner. 2012. The Malesian floristic interchange: Pages 138–163 Biotic Evolution and Environmental Change in Southeast Asia. Cambridge University Press.

Rubin, D. B. 1987. Multiple Imputation for Nonresponse in Surveys.

Russo, S. E., P. Brown, S. Tan, and S. J. Davies. 2008. Interspecific demographic trade-offs and soil-related habitat associations of tree species along resource gradients. Journal of Ecology 96:192–203.

Serafini, C., F. Cosentino, G. Amori, and L. Maiorano. 2025. Modelling Species Distribution at the Boundaries of the Earth’s Climate. Global Ecology and Biogeography 34:e70082.

Sharma, N., M. D. Behera, A. P. Das, and R. M. Panda. 2019. Plant richness pattern in an elevation gradient in the Eastern Himalaya. Biodiversity and Conservation 28:2085–2104.

Shih, C.-H., Y.-S. Jang, T.-Y. Yang, C. Huang, J.-Y. Juang, and M.-H. Lo. 2025. Impact of diurnal temperature and relative humidity hysteresis on atmospheric dryness in changing climates. Science advances 11:eadu5713.

Simmonds, E. G., K. P. Adjei, B. Cretois, L. Dickel, R. González-Gil, J. H. Laverick, C. P. Mandeville, E. G. Mandeville, O. Ovaskainen, J. Sicacha-Parada, E. S. Skarstein, and B. O’Hara. 2024. Recommendations for quantitative uncertainty consideration in ecology and evolution. Trends in Ecology & Evolution 39:328–337.

Stark, J. R., and J. D. Fridley. 2022. Microclimate-based species distribution models in complex forested terrain indicate widespread cryptic refugia under climate change. Global Ecology and Biogeography 31:562–575.

Steinbauer, M. J., R. Field, J.-A. Grytnes, P. Trigas, C. Ah-Peng, F. Attorre, H. J. B. Birks, P. A. V. Borges, P. Cardoso, C.-H. Chou, M. De Sanctis, M. M. de Sequeira, M. C. Duarte, R. B. Elias, J. M. Fernández-Palacios, R. Gabriel, R. E. Gereau, R. G. Gillespie, J. Greimler, D. E. V. Harter, T.-J. Huang, S. D. H. Irl, D. Jeanmonod, A. Jentsch, A. S. Jump, C. Kueffer, S. Nogué, R. Otto, J. Price, M. M. Romeiras, D. Strasberg, T. Stuessy, J.-C. Svenning, O. R. Vetaas, and C. Beierkuhnlein. 2016. Topography-driven isolation, speciation and a global increase of endemism with elevation. Global Ecology and Biogeography 25:1097–1107.

Still, C., P. Foster, and S. Schneider. 1999. Simulating the Effects of Climate Change on Tropical Montane Cloud Forests. Nature 398.

Sundqvist, M. K., N. J. Sanders, and D. A. Wardle. 2013. Community and Ecosystem Responses to Elevational Gradients: Processes, Mechanisms, and Insights for Global Change. Annual Review of Ecology, Evolution, and Systematics 44:261–280.

Tang, C. Q., T. Matsui, H. Ohashi, Y.-F. Dong, A. Momohara, S. Herrando-Moraira, S. Qian, Y. Yang, M. Ohsawa, H. T. Luu, P. J. Grote, P. V. Krestov, Ben LePage, M. Werger, K. Robertson, C. Hobohm, C.-Y. Wang, M.-C. Peng, X. Chen, H.-C. Wang, W.-H. Su, R. Zhou, S. Li, L.-Y. He, K. Yan, M.-Y. Zhu, J. Hu, R.-H. Yang, W.-J. Li, M. Tomita, Z.-L. Wu, H.-Z. Yan, G.-F. Zhang, H. He, S.-R. Yi, H. Gong, K. Song, D. Song, X.-S. Li, Z.-Y. Zhang, P.-B. Han, L.-Q. Shen, D.-S. Huang, K. Luo, and J. López-Pujol. 2018. Identifying long-term stable refugia for relict plant species in East Asia. Nature Communications 9:4488.

Tang, C. Q., and M. Ohsawa. 1997. Zonal transition of evergreen, deciduous, and coniferous forests along the altitudinal gradient on a humid subtropical mountain, Mt. Emei, Sichuan, China. Plant Ecology 133:63–78.

Thorne, J. H., H. Choe, L. Dorji, K. Yangden, D. Wangdi, Y. Phuntsho, and K. Beardsley. 2022. Species richness and turnover patterns for tropical and temperate plants on the elevation gradient of the eastern Himalayan Mountains. Frontiers in Ecology and Evolution 10.

Tredennick, A. T., G. Hooker, S. P. Ellner, and P. B. Adler. 2021. A practical guide to selecting models for exploration, inference, and prediction in ecology. Ecology 102:e03336.

Tremblay, R. L., A. J. Tyre, M.-E. Pérez, and J. D. Ackerman. 2021. Population projections from holey matrices: Using prior information to estimate rare transition events. Ecological Modelling 447:109526.

Trethowan, L. A., B. E. Walker, S. P. Bachman, C. Heatubun, P. Puradyatmika, H. Rustiami, and T. M. A. Utteridge. 2023. Plant species biogeographic origin shapes their current and future distribution on the world’s highest island mountain. Journal of Ecology 111:372–379.

Trouvé, R., S. Bunyavejchewin, and P. J. Baker. 2020. Disentangling fire intensity and species’ susceptibility to fire in a species-rich seasonal tropical forest. Journal of Ecology 108:1664–1676.

Vehtari, A., A. Gelman, and J. Gabry. 2017. Practical Bayesian model evaluation using leave-one-out cross-validation and WAIC. Statistics and Computing 27:1413–1432.

Wang, Z., J. Fang, Z. Tang, and X. Lin. 2010. Patterns, determinants and models of woody plant diversity in China. Proceedings of the Royal Society B: Biological Sciences 278:2122–2132.

Wangda, P., D. Gyaltshen, and T. Norbu. 2010. Influence of slope-aspect on the species composition and structural traits along the altitudinal gradients of the inner dry valleys. RNR Journal of Bhutan 6:73–87.

Wangda, P., and M. Ohsawa. 2006a. Structure and regeneration dynamics of dominant tree species along altitudinal gradient in a dry valley slopes of the Bhutan Himalaya. Forest ecology and management 230:136–150.

Wangda, P., and M. Ohsawa. 2006b. Gradational forest change along the climatically dry valley slopes of Bhutan in the midst of humid eastern Himalaya. Plant Ecology 186:109–128.

Ward Jr., J. H. 1963. Hierarchical Grouping to Optimize an Objective Function. Journal of the American Statistical Association 58:236–244.

White, A. E., K. K. Dey, D. Mohan, M. Stephens, and T. D. Price. 2019. Regional influences on community structure across the tropical-temperate divide. Nature Communications 10:2646.

Whitman, M., R. S. Beaman, R. Repin, K. Kitayama, S. I. Aiba, and S. E. Russo. 2021. Edaphic specialization and vegetation zones define elevational range-sizes for Mt Kinabalu regional flora. Ecography 44:1698–1709.

Whitman, M., and S. E. Russo. 2024. Biogeographic affiliation and centers of richness as predictors of elevational range-size patterns for Malesian flora. Ecography:e07043.

Wilson, A. M., and J. A. Silander Jr. 2014. Estimating uncertainty in daily weather interpolations: a Bayesian framework for developing climate surfaces. International Journal of Climatology 34:2573–2584.

Zhang, A., Y. Chen, X. Pan, S. Chen, W. Li, and Y. Fu. 2022. Entangled impacts of large-scale monsoon flows and terrain circulations on the diurnal cycle of rainfall over the Himalayas. Journal of the Atmospheric Sciences 79:301–316.

Zhu, H., and P. S. Ashton. 2021. Ecotones in the tropical-subtropical vegetation transition at the tropical margin of southern China. Chinese Science Bulletin 66:3732–3743.

Zhu, H., P. S. Ashton, B. Gu, S. Zhou, and Y. Tan. 2021. Tropical deciduous forest in Yunnan, southwestern China: Implications for geological and climatic histories from a little-known forest formation. Plant Diversity 43:444–451.

Zu, K., Z. Wang, X. Zhu, J. Lenoir, N. Shrestha, T. Lyu, A. Luo, Y. Li, C. Ji, S. Peng, J. Meng, and J. Zhou. 2021. Upward shift and elevational range contractions of subtropical mountain plants in response to climate change. Science of The Total Environment 783:146896.

